# Dissecting the conformational complexity and flipping mechanism of a prokaryotic heme transporter

**DOI:** 10.1101/2022.04.07.487047

**Authors:** Di Wu, Ahmad R Mehdipour, Franziska Finke, Hojjat G Goojani, Roan R Groh, Tamara N Grund, Thomas MB Reichhart, Rita Zimmermann, Sonja Welsch, Dirk Bald, Mark Shepherd, Gerhard Hummer, Schara Safarian

## Abstract

Iron-bound cyclic tetrapyrroles (hemes) are key redox-active cofactors in membrane-integrated oxygen reductases and other bioenergetic enzymes. However, the mechanisms of heme transport and insertion into respiratory chain complexes remain unclear. Here, we used a combination of cellular, biochemical, structural and computational methods to resolve ongoing controversies around the function of the heterodimeric bacterial ABC transporter CydDC. We provide multi-level evidence that CydDC is a heme transporter required for assembly and functional maturation of cytochrome *bd*, a pharmaceutically relevant drug target. Our systematic single-particle cryo-EM approach combined with atomistic molecular dynamics simulations provides detailed insight into the conformational landscape of CydDC during substrate binding and occlusion. Our simulations reveal that heme binds laterally from the membrane space to the transmembrane region of CydDC, enabled by a highly asymmetrical inward-facing CydDC conformation. During the binding process, heme propionates interact with positively charged residues on the surface and later in the substrate-binding pocket of the transporter, causing the heme orientation to flip 180 degrees. The membrane-accessible heme entry site of CydDC is primarily controlled by the conformational plasticity of CydD transmembrane helix 4, the extended cytoplasmic segment of which also couples heme confinement to a rotational movement of the CydC nucleotide-binding domain. Our cryo-EM data highlight that this signal transduction mechanism is necessary to drive conformational transitions toward occluded and outward-facing states.

**One Sentence Summary:** The heterodimeric bacterial ABC transporter CydDC is a heme flippase essential for the functional maturation of cytochrome *bd*.

## Introduction

Iron is the second most abundant metal on our planet and an essential trace element for all domains of life (*1*–*6*). It is involved in a multitude of physiological processes such as photosynthesis, protein biosynthesis, nitrogen fixation, nucleic acid repair, and respiration. Cellular iron is found in the form of iron-sulfur clusters (Fe-S), iron-bound cyclic tetrapyrroles (hemes), or in its free ionic forms. In biological systems, iron mediates electron transfer by acting as electron acceptor or donor in various biochemical reactions (*7*). Terminal respiratory oxidases such as the mitochondrial cytochrome *c* oxidase and the bacterial cytochrome *bd* oxidase are metalloproteins that rely on their redox active heme cofactors for reduction of molecular oxygen to water (*8*–*14*). Knowledge about the mechanism of maturation and heme insertion of cytochrome *bd*-type oxidases remains unknown but is urgently required for the development of antimicrobial drugs that can specifically target the energy metabolism and respiratory re-wiring of human pathogenic bacteria upon infection and proliferation (*15, 16*).

The ABC transporter CydDC is conserved in most bacteria and plays a central role in the biogenesis of membrane-integrated and soluble cytochromes (*17*–*21*). However, its precise function and molecular mode of action have remained enigmatic for decades (*22, 23*). CydDC was initially thought to translocate heme across the bacterial cytoplasmic membrane but later studies instead concluded that CydDC is involved in the regulation of the periplasmic redox poise by transporting the reductants glutathione (GSH) and L-cysteine (L-Cys) to the periplasm (*18, 20, 24*–*27*). Despite this widely accepted substrate specificity of CydDC, it was later found that the CydDC complex co-purifies with bound heme (*28*). This observation motivated the assumption that heme represents a prosthetic group of a periplasmic redox sensor domain for regulation of transporter activity (*17, 22*). A recent work fundamentally challenged the transporter function of CydDC and proposed that it operates as a membrane-bound enzyme catalyzing the reduction of cytoplasmic cystine (CSSC) to L-Cys (*29*).

To resolve these points of controversy, we used *Escherichia coli* as a model system amenable to our integrative approach including growth-complementation studies, biochemical activity assays, systematic single-particle electron cryo-microscopy (cryo-EM), and atomistic molecular dynamics (MD) simulations to characterize the function of CydDC down to the atomic level and to dissect its molecular mode of action. For our structural biology approach, we established a matrix of sample conditions based on a combination of putative substrate molecules (GSH, GSSG, L-Cys, CSSC, and heme), nucleotides that are either susceptible or resistant to hydrolysis (ATP, ADP, ATP+Vi, and AMP-PNP), and rationally designed mutant variants of CydDC. We determined structures of CydDC under 23 different sample conditions at resolutions of 2.7 to 3.9 Å (table S1, and figs. S1 to S7). Based on these results, we conclude that the primary role of CydDC is the transport of heme for functional maturation of cytochrome *bd* oxidases.

## Results

### CydDC is required for the functional maturation of both cytochrome *bd* isoforms

We carried out growth complementation studies using the *E. coli* mutant strain MB43, which lacks all terminal oxidases and thus shows impaired respiratory activity (*30*). The growth of this strain is poor compared to wild-type controls, but was improved following the introduction of genes encoding cytochromes *bd*-I (p*cydABX*) or *bd*-II (p*appCBX*) (Fig. 1A and fig. S8). Next, we constructed an isogenic deletion mutant of the *cydDC* operon in the genetic background of MB43. This strain, designated MB43Δ*cydDC*, shows an indistinguishable phenotype to MB43 (Fig. 1A). Unlike MB43 however, the growth of MB43Δ*cydDC* was not restored by introducing p*cydABX* or p*appCBX*. A growth-active phenotype of MB43Δ*cydDC* was only achieved by double complementation with p*cydDC* and either p*cydABX* or p*appCBX* (Fig. 1A). Accordingly, UV-Vis spectroscopy revealed that the characteristic fingerprints of cytochrome *bd* cofactors (hemes *b* and *d*) were absent in membrane fractions of MB43 and MB43Δ*cydDC* but clearly detected in the growth-restored complementation strains (Fig. 1B) (*30*– *33*).

**Fig. 1.**
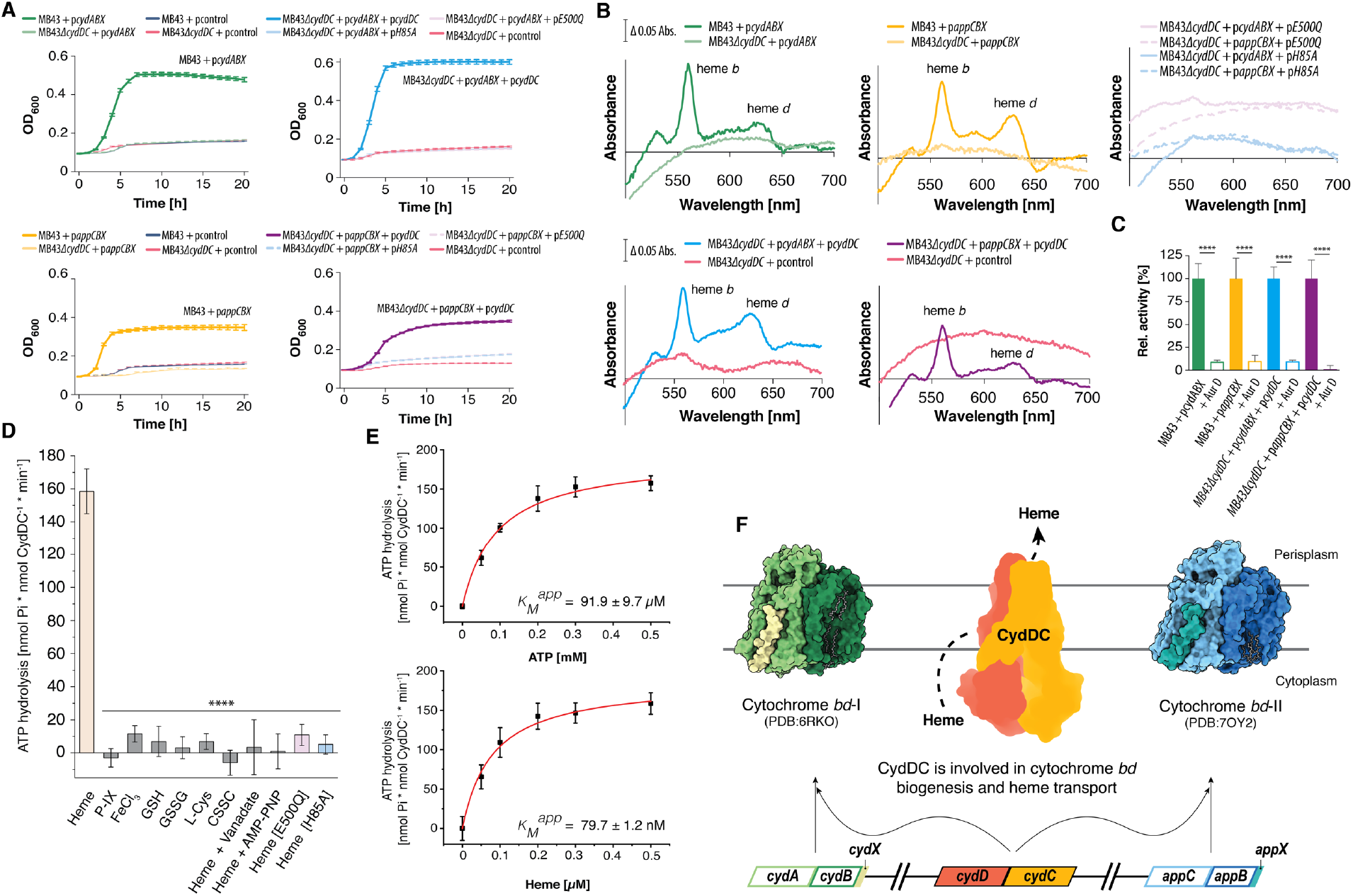
Physiological and biochemical links between CydDC activity and cytochrome *bd* biogenesis. **(A)** Growth complementation studies using *E. coli* MB43 and MB43*ΔcydDC*. Both strains lack NADH dehydrogenase I and all terminal oxidases. Cytochrome *bd* variants: p*cydABX* – wild-type cyt. *bd*-I; p*appCBX* – wild-type cyt. *bd*-II. CydDC variants: p*cydDC* – wild-type variant; p*E500Q* – glutamate to glutamine exchange at position 500 of CydC; p*H85A* – histidine to alanine exchange at position 85 of CydC. Control plasmid: pcontrol – empty pET17 vector. **(B)** Reduced-minus-oxidized UV-Vis spectra (450 - 700 nm) of membrane fractions from strains with restored/impaired growth phenotypes. Peaks corresponding to *b*-type and *d*-type hemes are indicated. **(C)** Oxygen reductase activity of membranes fractions from strains with growth-active phenotypes following complementation with structural genes of cytochrome *bd*-I, *bd*-II, and CydDC, respectively. Aurachin D (Aur. D) induced inhibition of oxygen consumption indicates for cytochrome *bd* specific oxygen reductase activity. Data are given as mean of n=3 ± SD. Significance was assessed based on a paired two-tailed Student’s t-test (**** *p* < 0.0001). **(D)** ATP hydrolysis activity of CydDC in the presence of putative substrate molecules. Data are given as mean of n=3 ± SD. Significance was assessed based on a paired two-tailed Student’s t-test (***, *p* < 0.001; ****, p < 0.0001). **(E)** Titration experiments for the determination of Michaelis constants (*K*_*M*_^*app*^) for ATP (91.9 ± 9.7 µM) and heme (79.7 ± 1.2 nM). All presented ATP hydrolysis data are corrected for background activity in the absence of substrate candidates. **(F)** Working model for substrate specificity and physiological role of CydDC based on independent physiological and biochemical data generated in this study.

Further, we confirmed oxygen reductase activity of membrane fractions from growth-restored MB43 and MB43Δ*cydDC* variants (Fig. 1C and fig. S9A). Finally, we used the highly specific inhibitor aurachin D to demonstrate that restored oxygen consumption was due to the activity of cytochrome *bd* variants (Fig. 1C). In summary, growth complementation studies, UV-Vis spectroscopy and oxygen consumption assays confirmed that CydDC is essential for the assembly of both known variants of cytochrome *bd* (Fig. 1F).

### ATPase activity of CydDC is stimulated by heme

We next tested affinity-purified CydDC for ATPase activity in the presence of various potential substrates using the Malachite green phosphate assay (fig. S1) (*25, 28, 29, 34*). In contrast to previous reports, we found that GSH and L-Cys (and their oxidized counterparts GSSG and CSSC) failed to induce hydrolysis, whereas heme stimulated the ATPase activity of CydDC in a concentration-dependent manner (Fig. 1, D and E, and fig. S10). This effect was only observed for heme itself, but not for its macrocycle scaffold Protoporphyrin IX, or free iron, suggesting that the complexed iron molecule might play a critical role in binding and coordination (Fig. 1D). For verification of our results, we employed a second conceptually different ATP hydrolysis assay based on an enzyme-coupled approach utilizing pyruvate kinase and lactate dehydrogenase (fig. S11A). Both assays confirm the ATPase stimulating effect of heme while showing that the tested reductants do not show typical effects of ABC transporter substrates. To bolster our finding of heme as a primary candidate for a substrate molecule of CydDC, we determined kinetic parameters for heme and ATP. We obtained *K*_*M*_^*app*^ and *V*_*max*_ values of 79.7 nM and 187.04 nmol [Pi] × nmol [CydDC]^-1^ × min^-1^ for heme, and 91.9 *µ*M and 191.58 nmol [Pi] π-π nmol [CydDC]^-1^ × min^-1^ for ATP (Fig. 1E, and fig. S11, B and C). These values are reasonable regarding the physiological occurrence of both substrates (*35, 36*).

Next, we generated glutamate to glutamine mutants of the Walker B domains of CydD and CydC to determine whether CydDC possesses a functionally degenerate nucleotide-binding domain (NBD) as found in other ABC transporters (*37*). In case of the CydD^E511Q^C mutant, we were not able obtain pure and homogeneous preparations of sufficient yield upon recombinant production. In contrast, the CydDC^E500Q^ mutant was stably purified and did not show any reduction in its T_M_ as compared to the wild-type variant (fig. S1). Using heme as substrate, we showed that the conserved Walker B glutamate of CydC is essential for ATPase activity of CydDC (Fig. 1D, and figs. S12A and S13). In line with this observed lack of enzymatic activity, transformation of MB43Δ*cydDC* with *pE500Q* (CydDC^E500Q^) in combination with either p*cydABX* or p*appCBX* did not restore bacterial growth (Fig. 1, A and B, and fig. S9, B and C). We thus conclude that NBD^C^ represents the canonical hydrolysis site and that CydD most presumably contains a degenerate NBD.

To verify our findings on the role of heme as a potential substrate molecule and to dissect the molecular basis of CydDC function, we employed a systematic single-particle cryo-EM approach for structure determination of CydDC in the presence of different putative substrate molecules, nucleotides, and inhibitors (table S1, and figs. S2 to S7). By solving CydDC structures in biochemically defined states and under turnover conditions, we gained an in-depth view of the conformational landscape and substrate binding modes of this transporter. This allows us to describe key events in substrate recognition, gating, and translocation across the cytoplasmic membrane. In the following, we will focus on the major conformations obtained and their roles in substrate binding, ATP hydrolysis, and substrate release.

### General architecture of CydDC

The nucleotide and substrate-free, inward-facing conformation of CydDC 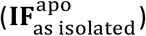, which was obtained in conditions lacking any externally added compounds, exhibits canonical features of type IV ABC transporters (Fig. 2, A and B) (*38*–*40*). Each subunit is composed of a membrane domain formed by six transmembrane α-helices (TMH), an N-terminal cytoplasmic elbow helix (EL) oriented parallel to the membrane plane, and a cytoplasmic nucleotide-binding domain (Fig. 2, A and B). Both NBDs feature a typical RecA-type fold and two peripheral ABC specific subdomains ABCα and ABCβ (Fig. 2A) (*41*). In this conformation the heterodimer adopts an asymmetrical structure with an overall Cα root-mean-square deviation (RMSD) of 4.17 Å (TMH: 3.28 Å; NBD: 2.5 Å) (fig. S14). Particularly TMs 4 and 5 show a pronounced conformational difference between CydD and CydC.

**Fig. 2.**
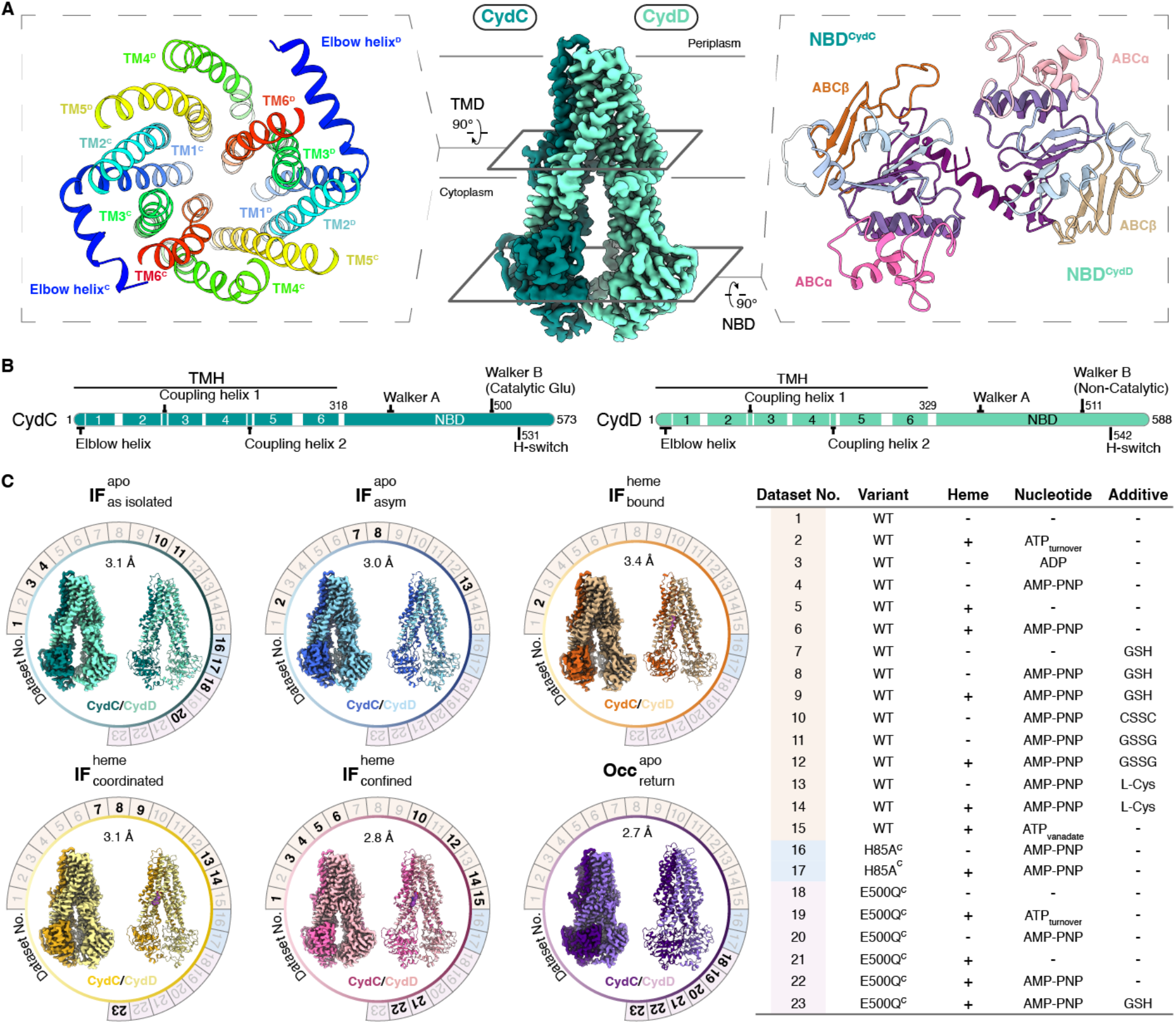
Systematic cryo-EM approach. **(A)** Volume map representation of the 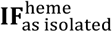 structure representing the general architecture of CydDC. Dark green, CydC; light green, CydD. Left: cross-section of the membrane domain composed of 12 TMs showing arrangement and geometry typical for type IV ABC transporters. Right: cross-section of nucleotide-binding domains (NBDs). The subdomains ABCα and ABCβ of each NBD are highlighted. **(B)** Schematic organization of CydC and CydD subunits. TMs are indicated by numbers. Residues important for ATP hydrolysis and signal transduction between TMH and NB domains are highlighted. **(C)** Summary of systematic single-particle cryo-EM studies. Left: volume maps and corresponding ribbon models within each circle represent a distinct conformation of CydDC. Numbers around circles refer to sample condition of CydDC analyzed by electron microscopy. Bold numbers indicate that the given conformation was present under that condition. Right: summarized information about the analyzed CydDC variants, presence and absence of heme, nucleotide, and additional putative substrate molecules. CydDC^wt^, beige; CydDC^H85A^, light blue; CydDC^E500Q^, purple. IF, inward-facing; Occ, occluded.

### Structural insights into substrate specificity

The successful determination of the 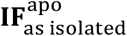 conformation was followed by a systematic screen for substrate molecules that locate within a specific binding pocket. We solved cryo-EM structures of CydDC in the presence of the putative substrate molecules GSH, GSSG, L-Cys, CSSC, and heme individually (Fig. 2C). Distinct densities within the TMH domain were neither obtained in the presence of reduced nor of oxidized forms of glutathione and cysteine. By contrast, we were able to unambiguously identify a bound heme molecule within the TMH domain (fig. S15). This finding is consistent with the ATP hydrolysis inducing effect of heme (Fig. 1D and fig. S11). In addition, we determined structures of CydDC in the presence of a combination of heme and thiols (GSH, GSSG, L-Cys) in order to assess whether these molecules may be heme-dependent co-substrates of CydDC. This possibility was, however, ruled out because we could identify ligand density only for heme but did not observe signal for any other added molecule. It is important to note that we obtained heme bound CydDC conformations not only through exogenous heme addition but consistently also from subsets of CydDC particles that natively co-purified with heme in sample compositions free of external heme (Fig. 2C). This observation further reinforces our working hypothesis that heme is the physiological substrate of CydDC (Fig. 1F).

### Heme binding and occlusion requires sequential conformational transitions

The binding of heme and the formation of an inward-facing confined state requires distinct conformational changes within the TMH domain of CydDC. From the sum of analyzed sample conditions, we obtained three heme-bound states of CydDC that provide comprehensive insight into this process and allow us to delineate conformational transitions during this process (Figs. 2C and 3, and figs. S15 and S16). Based on these structural insights, we can assign that heme initially binds to a cavity formed by transmembrane helices TM2^C^, TM3^C^, TM5^D^, and TM6^D^ that is in the vicinity of the lateral membrane plane (Fig. 3A, and figs. S15 and S16). In this 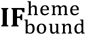 conformation, which was obtained exclusively under active turnover, the heme molecule primarily interacts with the invariant H85^C^ residue which functions as an axial ligand for the heme iron at a distance of 3.4 Å (Fig. 2C). Additionally, the two heme propionate groups are located in a reasonable distance to form electrostatic interactions with R81^C^ of TM2^C^ and R136^C^ of TM3^C^, respectively (Fig. 3A, figs. S15 and S16, and movie S1). Despite these key interactions, heme is not tightly bound since the distinct density features of the porphyrin macrocycle are less pronounced than the overall local resolution consistent with a dynamically bound substrate (fig. S15). We assign the 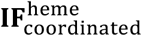 state as the second step in the binding cascade. Here, the interaction between CydDC and heme is strengthened by movement of the C-terminal segment of TM6^D^ toward the heme and concomitant sidechain rearrangements of H312^D^, which acts as a second axial ligand of the heme iron (Fe - H312^D^ distance 2.6 Å), and Y311^D^, which moves in front of the porphyrin plane and thereby prevents heme from escaping to the membrane space (Fig. 3A, figs. S15 and S16, and movie S2 and S3). The interaction between TM6^D^ and EL^D^ further causes positioning of the elbow helix closer towards TM4^D^, which narrows the membrane accessible gate to the heme binding site (Fig. 3D, figs. S15 and S16, and movie S3). Overall, the TMH region adopts a more compact state upon heme binding and coordination. This compaction is the result of small-scale movements of all transmembrane helices (except TM4^C^ and TM5^C^) towards the symmetry axis of the TMH domain (movie S2).

**Fig. 3.**
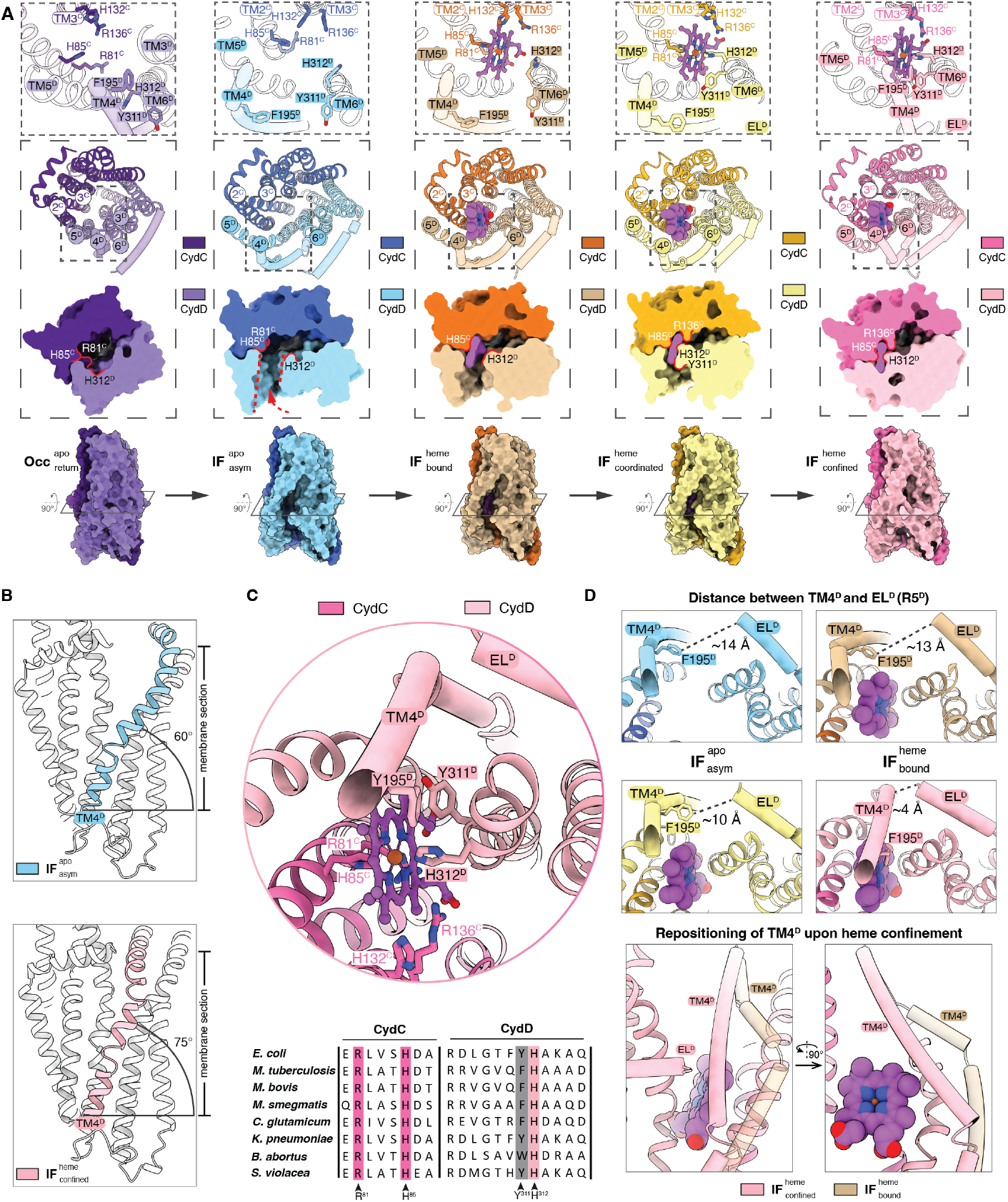
Conformational landscape of heme binding and translocation. (**A**) Closeup views show residues involved in heme binding and occlusion. Surface model cross sections illustrate the changing shape and size of the internal cavity in different conformational states. Surface model side views highlight the change in overall shape of the TMH region in different conformational states. Heme is shown as purple ball-and-stick model. EL1^D^ and TM4^D^ are shown as tubes. (**B**) Upon heme entry and coordination, the membrane-embedded segment TM4^D^ helix moves towards EL^D^ and thus closes the lateral gate of CydDC. **(C)** Top: closeup view of the heme binding site in the 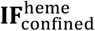 state. Heme is axially coordinated by two histidine residues (H85^C^ and H312^D^). Propionate groups of the heme molecule form electrostatic interactions with invariant R81^C^ and R136^C^. The heme macrocycle is further held in place by F195^D^ and Y311^D^, which restrict lateral movement towards the membrane and act as internal gate-forming residues. Bottom: sequence alignments of CydD and CydC heme binding-site forming regions of representative and disease related bacteria. Conserved residues are highlighted in magenta. Conserved amino acid groups are highlighted in gray. **(D)** Top: changing EL^D^ - TM4^D^ distance during the transition from 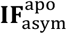 to 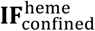 states shown by ribbon models viewed from the cytoplasmic angle. Distances are indicated by dashed lines. Bottom: closeup view of the membrane-exposed and sealed heme entry site in 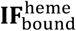 (beige) and 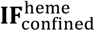 (magenta) states, respectively. During the transition from 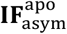 via 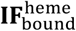 to 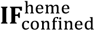 we observe that the cytoplasmic portion of TM4^D^ approaches the heme molecule and is positioned above it, contributing to substrate confinement. IF, inward-facing; OF, outward-facing; Occ, occluded; asym, asymmetrical.

The most abundant heme-bound conformation obtained (11 individual structures) is termed 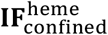 and represents the confined inward-facing TMH conformation (Figs. 2C and 3A). Upon binding and tight coordination of heme, a crucial conformational change of TM4^D^ is induced (movie S4). Movement of this 7.2 nm long extended transmembrane helix is occurring on two levels (movie S5 and S6). The membrane embedded segment of TM4^D^ (segment 164-195) changes its angle from ca. 60° to ca. 75° relative to the membrane plane and thus moves in front of the heme, blocking the lateral membrane gate (Fig. 3, B and C). The kinked C-terminal half of TM4^D^ that is exposed to the cytoplasm concomitantly moves towards EL^D^ and adopts a conformation that runs diagonally above the bound heme molecule (Fig. 3D). Accordingly, the side chain of F195^D^ moves in front of the heme and aligns in parallel with the porphyrin plane (Fig 3, A and C, and fig. S16). Together with Y311 of TM6^D^, these aromatic residues generate π-π stacking interactions with the porphyrin plane of the heme. An electrostatic interaction network formed between EL_D_ and TM4 further contributes to the formation of a tightly-sealed substrate access gate (Fig. 3, A and C, figs. S16 and S17, and movie S5). As a consequence, the central solvent-filled cavity of the TMH domain changes in size and position (Fig. 3A). The formation of a distinct heme-binding pocket narrows the unoccupied space within the TMH domain and establishes a lateral pathway perpendicular to the membrane plane that ends at the tightly interlocked periplasmic substrate exit gate (Fig. 3A). Furthermore, we observed that even in the absence of any nucleotide, if heme was added to CydDC we exclusively obtained the 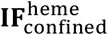 conformation, which provides evidence that binding and confinement of heme occurs independent of nucleotide occupancy of the NBDs (Fig. 2C).

Based on the deduced sequence of binding events and the distinct interaction patterns between heme and CydDC, we reasoned that the interaction between H85^C^ and the heme iron marks the most crucial step of substrate recognition and initiation of all subsequent events required to facilitate heme translocation across the membrane. To test this hypothesis, we generated a CydDC H85A^C^ mutant (p*H85A*) and performed growth-restoring complementation studies and oxygen consumption measurements using the MB43Δ*cydDC* strain (Fig. 1, A and B, and fig. S9, B and C). In addition, we investigated the effect of the histidine to alanine replacement on ATP hydrolysis activity of CydDC^H85A^ (Fig. 1D). Results of these experiments revealed that lack of this highly conserved histidine residue abrogates the functionality of CydDC (Fig. 1, A, B and D, and figs. S9 and 12B). Furthermore, in contrast to wild-type CydDC, CydDC^H85A^ could not be co-purified with bound heme and could not be loaded by the exogenous addition of heme (fig. S18). These observations were confirmed by cryo-EM structures of CydDC^H85A^, exclusively adopting the 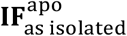 conformation, irrespective of presence or absence of exogenously added heme to the samples (Fig. 2C).

By analyzing the relationship between sample conditions and obtained CydDC conformations we recognized that the 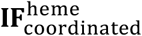 conformation was obtained only in the presence of the reductants GSH and L-Cys. The fact that their oxidized counterparts do not have a similar effect on CydDC could suggest that the reductive properties of GSH and L-Cys are responsible for stabilizing the 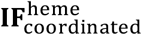 state. Further, we reason that the systematic absence of 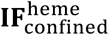 state in the presence of GSH and L-Cys together with the consistent observation of 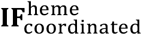 as the major heme-bound class indicates that these reductants cause opening of the closed lateral heme entry gate. Lack of clearly distinguishable ligand density in the cryo-EM maps of the respective datasets leaves open the question about the precise interaction sites of these thiols and the underlying molecular mechanisms that result in the stabilization of the 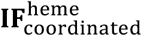 conformation. Further, it remains elusive whether these effects indicate for a physiological link between CydDC function and the two reductants or whether they have to be attributed to the high excess of thiols resulting unspecific reductive effects.

### MD simulations suggests that heme enters CydDC via the membrane space and is flipped by 180°

To map possible routes of heme entry, we carried out multiple atomistic MD simulations (n=10), focusing on the behavior of heme near lipid bilayer surfaces and the substrate binding pocket of CydDC (Fig. 4 and fig. S19). As a starting point we placed heme near a heterogeneous membrane composed of 70% POPE, 25% POPG and 5% cardiolipin. In 7 out of 10 of our repeated simulations, heme rapidly partitioned into the lipid bilayer, which is consistent with earlier experimental reports about interactions of heme with biomembranes (fig. S19) (*42, 43*). The porphyrin scaffold immersed deeply into the membrane core and interacted with lipid acyl chains, while the two propionate groups orientated towards the lipid head groups and interacted mostly with the ethanolamine groups of POPE molecules (fig. S19). Next, we analyzed the dynamics of heme placed near the lateral entry site of CydDC in the 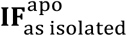 state. Our simulations show that the heme propionates interact with positively charged residues first on the protein surface and then the inside of the binding cavity. These latter interactions induce a rotation of the heme by 90° (Fig. 4). The observation of spontaneous heme binding and rotation suggests this pathway as the primary route for heme entry.

**Fig. 4.**
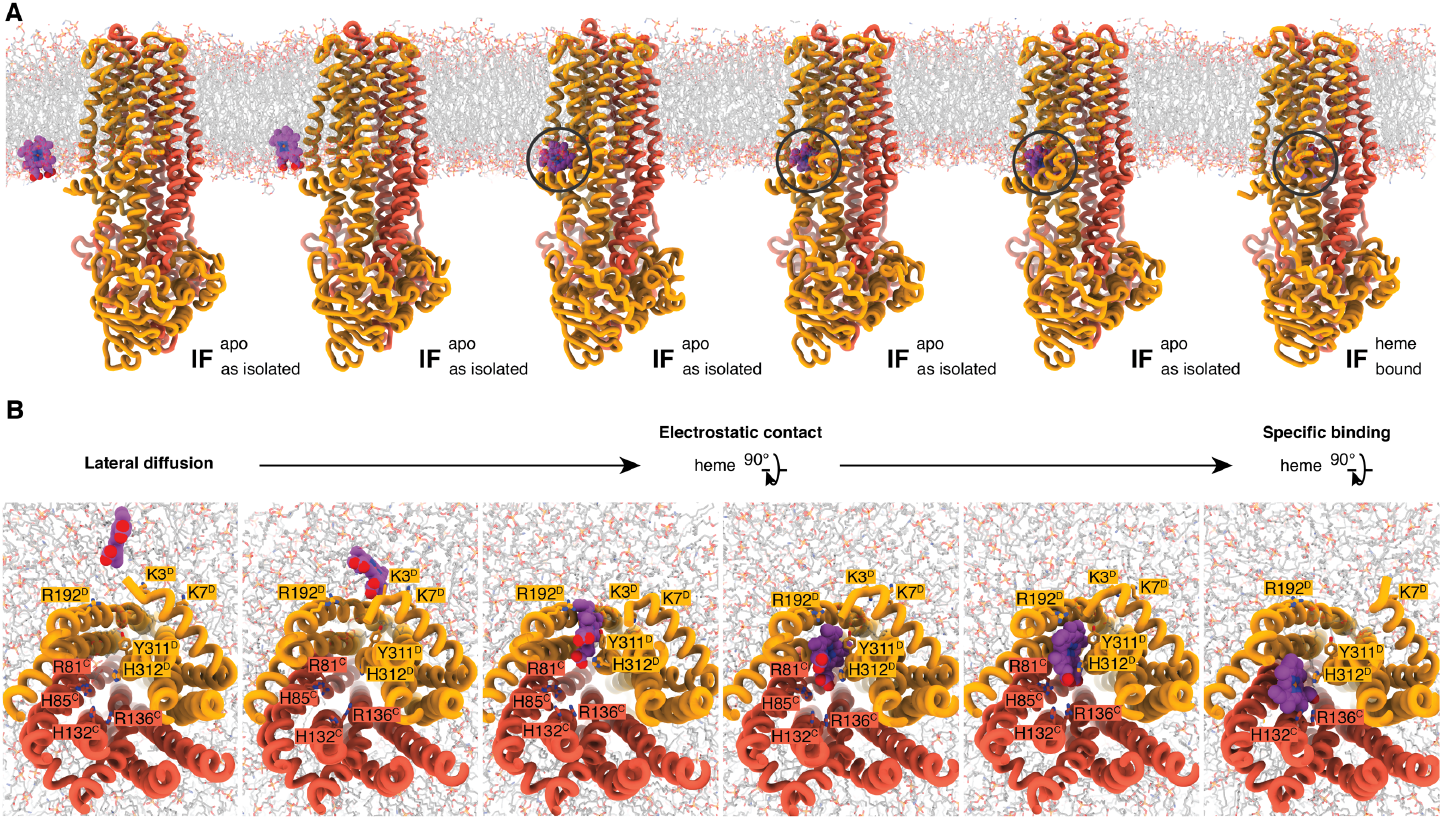
Structural dynamics of the heme binding and flipping mechanism. **(A)** Representative snapshots of MD simulations showing the lateral diffusion and entry of heme to the membrane-accessible binding cavity of CydDC. **(B)** Closeup views show that once heme interacts with the electrostatic surface and the interior of the binding cavity of CydDC, a 90° rotation occurs that orients the propionate groups of the porphyrin macrocycle horizontally. A second 90° rotation takes place upon bilateral axial coordination via H85^C^ and H312^D^, which results in a complete flip of the heme molecule compared to its conformation when immersed freely within the lipid bilayer.

We carried out further MD simulations using models of 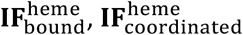, and 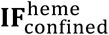 When heme is placed within the binding pocket without H85^C^-Fe coordination, it relaxes toward the entry conformation observed after spontaneous heme binding to 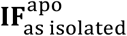 and shows significant mobility within the binding site. However, when heme is axially ligated to H85^C^ in 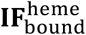, or both H85^C^ and H312^D^ in 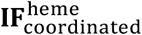 and 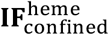, it maintains the conformation we observed in our cryo-EM structures. We conclude that heme is rotating 90° as it enters the binding site and a further 90° upon binding and axial ligation. This reorientation mechanism may prime heme for directed release toward the periplasmic space or the outer membrane leaflet once the transporter adopts its outward-facing release conformation (Fig. 4).

These results suggest that CydDC operates via a trap-and-flip mechanism similar to that described for the MFS type proton-dependent lipid transporter LtaA and other primary and secondary active lipid transporters (*44*–*48*). In MsbA and PfMATE, lipid flipping is primarily driven by electrostatic interactions between lipid head groups and side chains of the respective transporters (*45, 48*). This functional analogy between lipid and heme flipping indicates that the translocation of amphipathic molecules across biological membranes is presumably based on a common underlying mechanism.

### Substrate release is ATP dependent but not hydrolysis dependent

Next, we aimed to characterize the mechanistic relationship between substate transport and ATP hydrolysis. Active turnover conditions of wild-type CydDC promote two catalytic states as captured by cryo-EM: 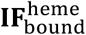 and a highly asymmetrical heme-free inward-facing conformation with nucleotide bound to both NBDs 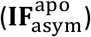 (Fig. 2C). Lack of additional conformations suggests that formation of 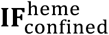 and subsequent translocation of heme to the periplasmic space via outward-facing conformations occurs rapidly and features short-lived intermediate states. To capture further transient conformations during the transport cycle, we performed an additional turnover cryo-EM experiment using CydDC^E500Q^ that has a significantly reduced hydrolysis activity (Figs. 1 and 2C). This cryo-EM experiment revealed an additional conformation closely matching the TmrAB return state (RMSD = 1.32) characterized by Hofmann *et al*. (*49*). Due to the lack of heme in the characterized binding site despite its presence in the sample, we assign this fully occluded conformation as the pre-hydrolysis return state of CydDC 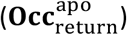 following substrate release (Fig. 3A and fig. S16). In contrast to the turnover dataset of CydDC^E500Q^, cryo-EM experiments of biochemically defined CydDC^E500Q^ samples revealed conformational heterogeneity reflected by structures of 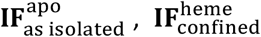, and 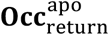 states (Fig. 2C). In the presence of heme and ATP (turnover conditions) this heterogeneity becomes resolved by fully driving CydDC^E500Q^ into the 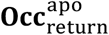 state (Fig. 2C). As this effect is not achievable by the addition of heme alone and given the substantially diminished ATPase activity of the CydDC^E500Q^ mutant, we conclude that ATP binding but not its hydrolysis is required for the transition from 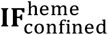 to 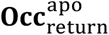 state via outward-facing (**OF**) heme-release states (Fig. 1D).

### Signal transduction and conformational coupling between TMH and NB domains

The transport of heme involves changes in the inter- and intra-subunit symmetry of CydDC. The 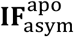 conformation features the greatest structural asymmetry with an overall Cα RMSD of 5.4 Å (TMH: 3.69 Å; NBD: 1.91 Å) (fig. S14). Loss of symmetry between CydD and CydC in the 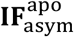 state is caused by the rotational movement of each subunit along the transporter’s central axis perpendicular to the membrane plane (movie S7). In this wedge-like conformation, the distal side (relative to the heme binding domain) of CydDC moves closer together than the proximal half whereby ABCα^C^ and ABCβ^D^ form a semi-interlocked NBD dimer (Fig. 5, A and B, and movie S7). This rearrangement increases the distance between the analogous NBDs of the proximal side, and dilates the heme-accessible membrane cavity (Figs. 3A and 5B). A central question concerning the functionality of ABC transporters is how the information about binding of a substrate molecule to the membrane domain is transduced to the soluble NBDs to induce a fully occluded state for subsequent substrate release and ATP hydrolysis. In CydDC, TM4^D^ has a dual function during the transition from 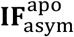 Via 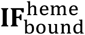 to 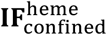 state. As described above, the membrane-integrated segment of TM4^D^ strongly contributes to sealing the bound heme molecule from the membrane environment (Fig. 3, and figs. S15 and S16). Beyond this function, we found that movement and rearrangement of the cytoplasmic segment of TM4^D^ is associated with a 25° rotation of the horizontal coupling helix 2 (CH2^D^) connecting TM5^D^ and TM4^D^. The contact of CH2^D^ with NBD^C^ then establishes a conformational coupling that leads to an almost parallel alignment of the two NBDs, which remain separated by ca. 6-8 Å until ATP is bound to the catalytic nucleotide-binding domain of CydC (Fig. 5A and movie S8).

**Fig. 5.**
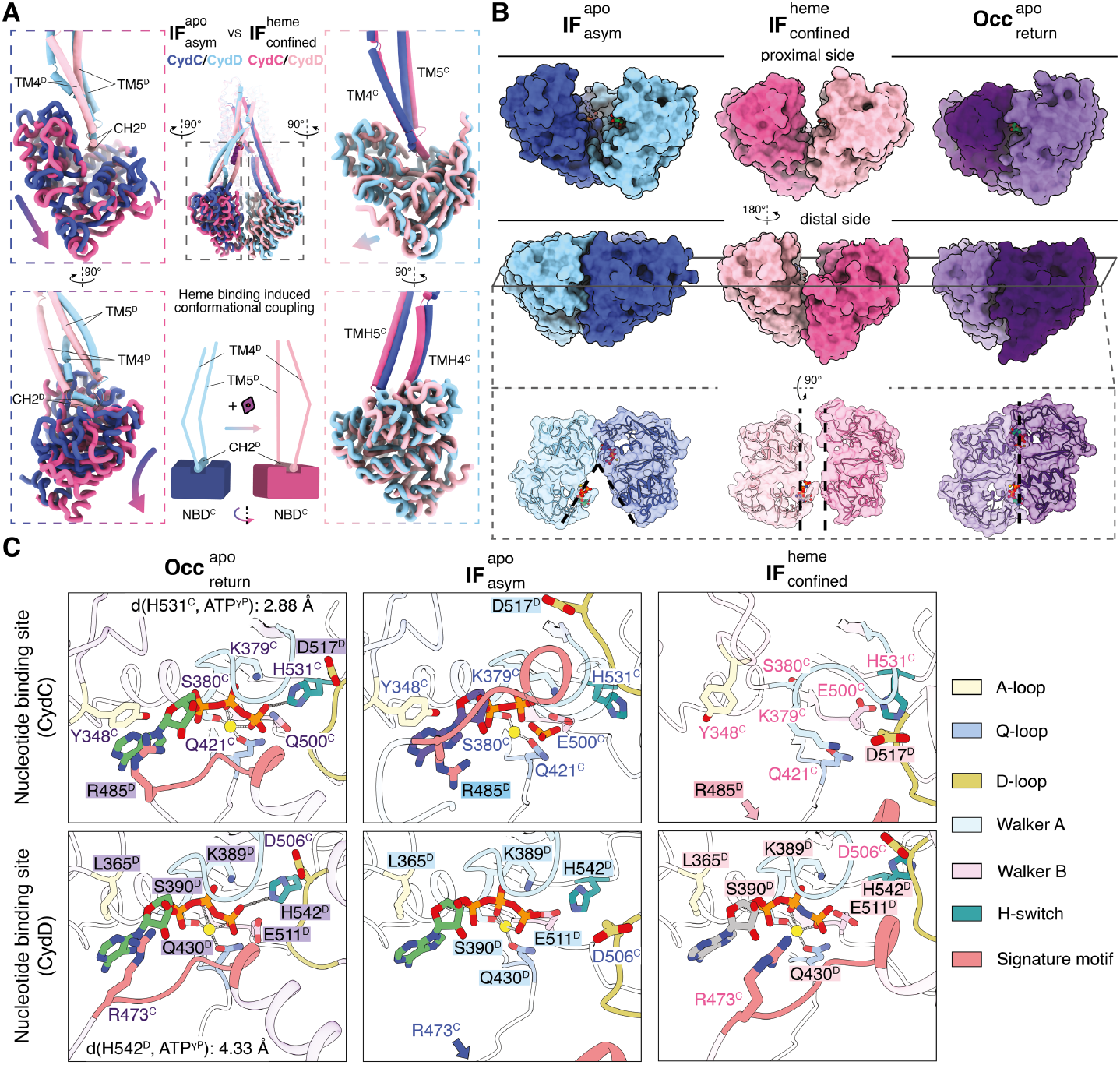
Signal transduction between TMH and NB domains. **(A)** Mechanism of conformational coupling between TMH and NB domains. Occlusion of the lateral substrate gate induces conformational changes of TMs 4 & 5 of CydD causing a rotational movement of NBD^C^ via coupling helix 2 (CH2^D^). No conformational coupling between TMs of CydC and NBD^D^ is observable. Dark blue, CydC; light blue, CydD; dark pink, CydC; light pink, CydD. **(B)** Surface representation of NBD-NBD interactions in proximal and distal view. Cross section top view on the level of the nucleotide-binding sites (NBS) shows distinctly different NBD conformations. 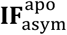, blue; 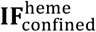, magenta; 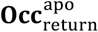, purple. **(C)** Closeup view of nucleotide-binding pockets in different conformational states and nucleotide occupancies. Conserved structural motifs of the nucleotide-binding and hydrolysis sites are highlighted. ATP, green; ADP, purple; AMP-PNP, grey; phosphate, orange; Mg^2+^, yellow.

### CydDC features a unique nucleotide exchange mechanism

In 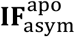 we discovered a previously unknown mode of interaction between the CydD signature loop and nucleotide bound to the canonical nucleotide-binding site (NBS) of CydC (Fig. 5C). During the asymmetrical interaction of the two NBDs on the distal side, the nucleotide is bound to NBS^C^ and its adenine base is coordinated by Y348^C^ of the A-loop via a π-π interaction on one side, and by a π-cation interaction via R485^D^ of the signature loop from CydD on the opposite side. However, in contrast to previously described fully-interlocked NBD dimers, as for instance from TmrAB^46^, the signature loop is separated from the γ-phosphate of the nucleotide by 12.6 Å (Fig. 5C and fig. S20). The degenerate NBS^D^ has a similar geometry in the 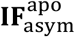 state, with the exception that adenine does not participate in π-π stacking and π-cation interactions. This is because of the uncharged L365^D^ residue at the analogous position of Y348^C^ in the Walker A motif of CydD, and the fact that the signature loop of CydC is placed at a distance of 20 Å to NBS^D^ (fig. S20).

A peculiar and thus far unreported NBS conformation was identified in the 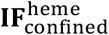 state. The conformational coupling between substrate occlusion and signal transduction (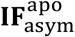 to 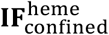) to the NBDs results in the collapse of the NBS^C^ region and the migration of the Walker A motif into the space originally occupied by the nucleotide phosphate groups (Fig. 5C and movie S9). During this process, the conformation of NBS^D^ remains mostly unchanged while the signature loop of CydC moves closer towards the nucleotide, although still separated from it by 10 Å. It is noteworthy that in none of the determined 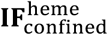 structures, we were able to observe nucleotide bound to NBS^C^, even in conditions where nucleotide had been added. In contrast, in all cases where externally added nucleotides were present in the sample, a clear density was identified at NBS^D^. This observation suggests that the collapsed conformation of NBS^C^ lacks affinity for the tested nucleotides. A second structural effect of the reductants GSH and L-Cys on CydDC was observed on the level of the NBDs. By a mechanism unknown, both GSH and L-Cys cause stabilization of the distal NBD-NBD interaction and allow for nucleotide binding at NBS^C^. While this NBD conformation was otherwise only observed during active turnover, GSH and L-Cys appear to stably induce this NBD state even in the presence of heme which otherwise induces the 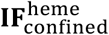 conformation and causes the dissociation of the NBDs and collapsing of NBS^C^.

Sequence analysis of critical structural motifs required for ATP hydrolysis did not reveal known mutations that would abolish ATP hydrolysis by NBD^D^. However, the tightly interlocked conformation of 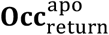 provides structural insights into differences between NBDs that may explain hydrolysis deficiency of NBD^D^ (Figs. 5C and 6, and figs. S20 and S21). The primary difference between ATP bound NBD^D^ and NBD^C^ of this pre-hydrolysis state is the respective distance between the switch histidine and the γ-phosphate of ATP. While H531^C^ is in close distance (2.9 Å) to act as catalytic acid/base during hydrolysis, H542^D^ is positioned in a less favorable distance of 4.4 Å to participate in polar interactions (Fig. 5C). In addition to this, the weak coordination of the adenosine base, also found in other heterodimeric ABC transporters, might influence the binding stability of the nucleotide at this position (Figs. 5C and 6, and fig. S20) (*49*). Here, it is important to mention that when we determined the 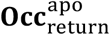 state structure from a sample that did not include exogenously added ATP or AMP-PNP, we observed lack of bound nucleotide at NBS^D^ but not at NBS^C^ (Figs. 2C and 6A). Loss of the NBS^D^ nucleotide might be possible by the presence of a solvent accessible cavity that remains open towards NBS^D^ even with interlocked NBDs in this conformation, whereas the tight interaction between the A-loop of CydC and ABCα of CydD forms an enclosed inner environment for ATP around NBS^C^ (Fig. 6B and fig. S21). Given the hydrolysis deficiency of CydDC^E500Q^ and the associated conformational differences observed between turnover datasets of the wild-type and CydDC^E500Q^ variants, we assign the consistently co-purified nucleotide bound to NBS^C^ as ATP.

**Fig. 6.**
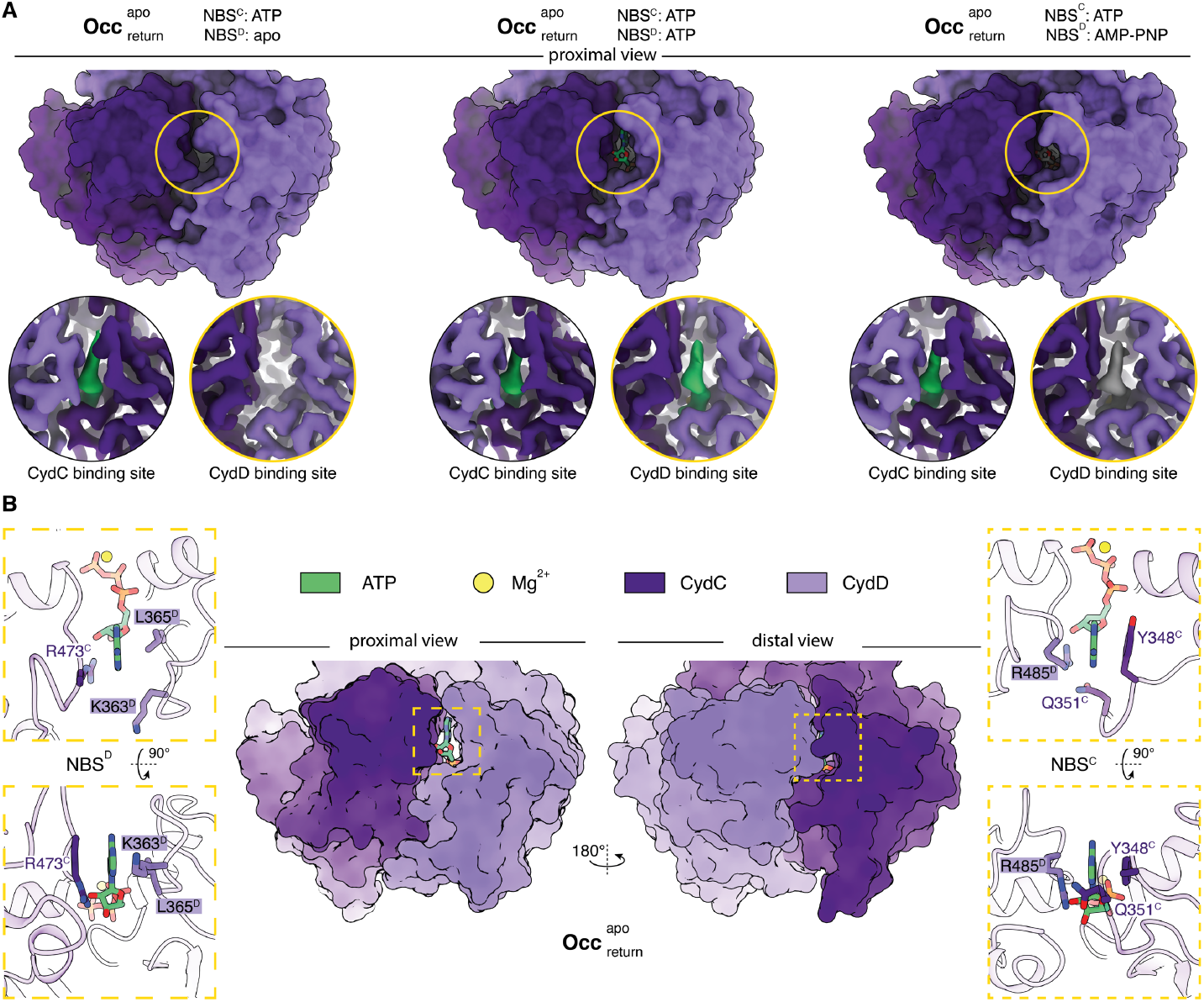
Local environment and accessibility of NBS^D^ and NBS^C^. **(A)** Top panel: proximal view of tightly-interlocked NBDs in the 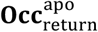 state from CydDC^E500Q^ shown as surface models (datasets 17-19). Bottom panel: closeup views of the NBS regions of CydD and CydC show sample-condition specific differences in nucleotide occupancy. ATP densities are shown in green, AMP-PNP densities are shown in grey. Densities of CydD and CydC are shown in dark and light purple, respectively. Bound ATP at NBS^C^ is enclosed by tight interactions between the CydC A-loop and the ABCα region of CydD in adjacency of the adenine group. The corresponding loops at NBS^D^ are separated widely and expose ATP bound at this site to the solvent. **(B)** Center panel: surface model of the interlocked NBD dimer in the 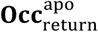 state from the turnover dataset of CydDC^E500Q^. Left panel: closeup view of ATP bound to NBS^D^ in top and side view orientation. Residues forming the local protein environment close to the adenine head group are shown. Right panel: closeup view of ATP bound to NBS^C^ in top and side view orientation. Residues forming the local protein environment close to the adenine head group are shown. ATP, green; Mg^2+^, yellow; CydC, light purple; CydD, dark purple.

### Modelling of outward-facing substrate-release conformations of CydDC

To complement our cryo-EM data, we modeled important but experimentally inaccessible short-lived conformational states during the transition from 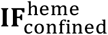 to 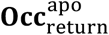. Our simulation-based modeling allowed us to construct three transient CydDC conformations representing distinct states of the transport cycle (fig. S22). The 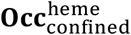 state represents heme-loaded CydDC upon ATP binding with tightly interlocked NBDs. To obtain this conformation, we performed an MD simulation for 400 ns of the fully occluded state of CydDC without heme 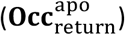. Further simulations were carried out to generate the 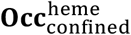 conformation by inserting heme into the binding site using the slow-growth procedure. After generating the initial model, we performed 300 ns of simulation and analyzed the structural stability by RMSD evolution and conformation dynamics. Our final model shows that the transition from 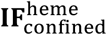 to 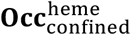 entails concentered movements of TM3^C^ and TM4^D^. This seals the cytoplasmic gate from solvent access and results in a pseudo-symmetric heterodimer conformation (fig. S22 and movie S10).

Starting from the 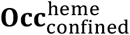 and 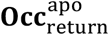 states we obtained two outward-open states (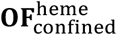 and **OF**^apo^) by performing steered MD simulations on the two periplasmic halves of CydDC. Having generated initial models, we checked the stability and the degree of opening after 300 ns of unrestrained simulations. Our outward facing models suggest that heme is released in a sequential process. First, the periplasmic gate formed by all twelve TMs is opened via separation of two lobes composed of TMs 1-2 of CydC and 4-6 of CydD, and vice versa of TMs 1-2 of CydD and 4-6 of CydC 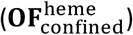 (fig. S23 and movie S11). These movements cause the lateral cavity next to the heme binding site to enlarge and form an exit pathway. During periplasmic gate-opening we found that the overall interaction of heme with the residues in the binding site remained unchanged. Accordingly, we conclude that substrate release is initiated by retraction of H312 of TM6^D^ and H85 of TM2^C^ (figs. S22 and S23, and movie S12). These events most likely change the local affinity and trigger the release towards the periplasmic side of the membrane. The post-release **OF**^apo^ conformation features a narrower periplasmic gate than 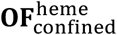. With the complete closing of the periplasmic gate, the transporter adopts its pre-hydrolysis 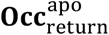 conformation as observed in the cryo-EM structures of CydDC^E500Q^ (Fig. 2C and fig. S23). In this state, TM6^D^ and TM2^C^ already adopt the 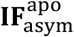 conformation, which would ensure rapid binding of another heme molecule upon ATP hydrolysis and lateral opening of the substrate entry gate (figs. S22 and S23, and movie S13).

## Conclusions

The molecular basis of CydDC function has been contested for 30 years. The results from various *in vitro* and *in vivo* studies led to conflicting conclusions about its role in bacterial physiology, and its function as a heme transporter was previously ruled out (*20, 25, 26*). In this study, we provided evidence that translocation of heme from the cytoplasmic space to the periplasm is the primary function of this ABC transporter.

Based on our multi-layered approach, we suggest a model for the transport cycle that integrates the results of the employed complementary methods (fig. S24). We assign the ATP-bound 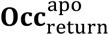 state as the starting conformation. We biochemically confirmed that NBD^C^ is essential for the ATPase activity of CydDC (Fig. 1D). In line with this, our structural data indicate that hydrolysis of ATP at this site is required for a mechanochemically-induced return to the substrate-free 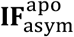 state (Figs. 3A and 5C, and fig. S16). By this rationale, we assign the densities at NBS^C^ of this post-hydrolysis state obtained from the turnover dataset of CydDC to ADP and Pi (Fig. 5C and fig. S20). We further conclude that heme binding to the TMH domain and ATP to NBD^C^ occurs sequentially rather than simultaneously, given that the binding and occlusion of heme 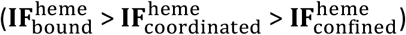 is needed to induce dissociation of the distal NBD-NBD interaction, causing NBS^C^ to collapse and release of ADP and Pi (Figs. 3 to 5, and movies S7 to S9). Given the fact that heme binding itself is not sufficient to drive the conformational transition toward the return state conformation but merely results in enrichment of the 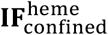 population, we conclude that binding of ATP to NBS^C^ is essential for the transition to the fully occluded 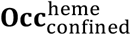 state. In our simulations, this state is followed by the opening of the extracellular gate, the retraction of TM6^C^, including the axial heme ligand H312^D^, to adopt the inward facing conformation of this helix, and the collapsing of the heme binding site (**OF**^apo^) (figs. S22 and S23, and movies S10 to S13). After release of the heme to the periplasmic space, the extracellular gate closes again, reverting back to the 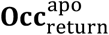 conformation. This is in agreement with the observation that under turnover conditions of CydDC^E500Q^ in which heme and ATP are present, we exclusively captured the occluded return state conformation. Notably, this is the sole analyzed condition, that does not result in a heme bound conformation of CydDC. Lack of ATP hydrolysis activity of the CydDC^E500Q^ variant traps the transporter in the post-substrate-release state which suggests that the key role of ATP hydrolysis is to convert CydDC to the 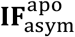, as observed in the turnover dataset of CydDC^wt^.

The transitional cycle of asymmetrical, symmetry-approaching and pseudo-symmetrical conformations during heme transport is unique to CydDC and highlights the mechanistic possibilities for the specialized substrate transport facilitated by heterodimeric ABC transporters and their diverse conformational space (fig. S25). This and previous structural studies of heme-transporting membrane proteins reveal mechanistic and architectural differences reflecting an evolutionary diversity that may be associated with specific physiological pathways and processes these transporters are involved in (*50*–*54*).

While we cannot fully exclude the possibility that CydDC might be substrate promiscuous and that the previously suggested substrates GSH and L-Cys might tightly bind to CydDC and induce ATP hydrolysis under conditions not met in this work, we can conclude that the heme-induced ATPase stimulation, the highly specific heme-binding pocket, the abrogation of ATPase activity upon mutation of a key heme-binding residue within the TMH domain, native heme remaining bound to this site after solubilization and extraction of CydDC, and the large-scale conformational responses to addition of external heme all indicate for a primary role as a heme transporter. The structural effects of GSH and L-Cys on CydDC observed in our study further suggest that it may be possible that these reductants could hold a regulatory role for the heme transporter function of CydDC. At present, this latter possibility remains a working hypothesis whose confirmation requires in-depth physiological, biochemical, and structural work.

Lack of CydDC is associated with extensive changes in bacterial metabolism, including an over-oxidized periplasm and perturbed respiratory activity (*17*). A crucial link is found between CydDC and the bacterial terminal oxygen reductase cytochrome *bd*, which relies on the function of CydDC for the insertion of its heme cofactors and functional maturation. This dependency is also reflected in their genomic organization as in most bacteria the structural genes of cytochrome *bd* and CydDC are operonically encoded in the same gene cluster (*55*). In infection studies, Δ*cydC* mutants of *Mycobacterium tuberculosis and Brucella abortus* displayed diminished survival in mouse models (*56, 57*). Inactivation of CydDC in *M. tuberculosis* also interfered with bacterial persistence during treatment with the front-line drug isoniazid and caused hypersusceptibility to compounds targeting the mycobacterial cytochrome *bc*_1_ complex, with bioenergetic profiles indistinguishable from cytochrome *bd* knock-out strains (*58, 59*). Hence, CydDC represents a viable target for the development of antibacterial drugs attacking bacterial respiration on the level of assembly.

While our current work provides evidence on a molecular level for the functional link between the heme transport function of CydDC and functional maturation of bacterial cytochromes, two central questions for future research arise. First, are additional and yet unknown periplasmic proteins involved in the process of cytochrome *bd* assembly and cofactor insertion? A putative candidate for a periplasmic chaperone is the heme-binding protein NikA, which is more abundant in *E. coli* strains overexpressing CydDC (*60*). This co-regulation suggests a functional link between CydDC and NikA. However, given the localization of cytochrome *bd* prosthetic heme groups close to the periplasmic membrane surface, it is also conceivable that CydDC could directly interact with the nascent CydA polypeptide and thus might be actively involved in loading heme cofactors to the catalytic subunit of cytochrome *bd*. A first evidence in favor of this scenario is found in *Francisella tularensis*, where a molecular interaction between CydC and CydA was reported based on co-migration patterns in BN-PAGE experiments (*61*). Second, does CydDC functions as a heme scavenger by shifting the equilibrium of membrane-dissolved heme towards the periplasmic leaflet? As shown in previous experimental studies and our MD simulations, partitioning of heme from the aqueous phase into membranes occurs rapidly. However, the transmembrane movement of heme from one bilayer leaflet to the other occurs very slowly (< 1 s^-1^) (*42*). The kinetic properties of CydDC suggest that heme translocation occurs at much higher rates, indicating that CydDC can shift the equilibrium of membrane-integrated heme towards the periplasmic leaflet. It remains unclear, whether CydDC releases heme to the periplasmic bilayer leaflet, the periplasmic membrane space, or whether it might deliver it directly to another protein. Based on its flippase function, we infer that CydDC could efficiently release heme towards the periplasmic membrane space because the exiting orientation of the heme macrocycle would readily promote electrostatic interactions between the propionate side chains and the periplasmic lipid head groups upon release.

## Acknowledgements

We thank Alexander Hahn and Gregory Cook for valuable discussions. We thank Hartmut Michel for support and providing infrastructural resources.

## Author contributions

D.W purified CydDC, prepared grids, collected cryo-EM data, processed cryo-EM data, refined the structure, built the model, co-drafted the manuscript, and prepared figures. F.F purified CydDC, prepared grids, collected cryo-EM data, performed biochemical activity assays, analyzed data, and drafted figures. H.G.G performed bacterial growth assays, determined oxygen consumption rates, and collected UV-Vis spectra. A.R.M performed MD simulations, co-drafted the manuscript, and prepared figures. T.N.G performed biochemical activity assays and analyzed data. R.G optimized purification conditions and performed initial cryo-EM screening experiments. T.R implemented purification and optimized CydDC buffer conditions. R.Z assisted in cell culturing and protein purification. M.S provided p*cydDC* and p*H85A*, and established the enzyme coupled ATPase assay. S.W calibrated and aligned the microscope. M.S and S.S initiated the project. G.H, D.B, M.S, and S.S supervised the project. S.S designed research, evaluated data, funded the project, drafted the manuscript, and generated figures.

## Data availability

Cryo-EM maps are deposited at the Electron Microscopy Data Bank under accession numbers: <u>EMD-14636</u>, <u>EMD-14638</u>, <u>EMD-14639</u>, <u>EMD-14640</u>, <u>EMD-14641</u>, <u>EMD-14642</u>, <u>EMD-14643</u>, <u>EMD-14644</u>, <u>EMD-14645</u>, <u>EMD-14646</u>, <u>EMD-14647</u>, <u>EMD-14649</u>, <u>EMD-14652</u>, <u>EMD-14653</u>, <u>EMD-14654</u>, <u>EMD-14655</u>, <u>EMD-14656</u>, <u>EMD-14657</u>, <u>EMD-14659</u>, <u>EMD-14660</u>, <u>EMD-14662</u>, <u>EMD-14663</u>, <u>EMD-14665</u>, <u>EMD-14667</u>, <u>EMD-14668</u>, <u>EMD-14669</u>, <u>EMD-14670</u>, <u>EMD-14671</u>, <u>EMD-14672</u>, <u>EMD-14673</u>, <u>EMD-14674</u>, <u>EMD-14675</u>, <u>EMD-14676</u>, <u>EMD-14684</u>, <u>EMD-14689</u>, <u>EMD-15264</u>, <u>EMD-15265</u>. Atomic models of CydDC have been deposited to the Protein Data Bank under accession numbers: <u>7ZD5</u>, <u>7ZDA</u>, <u>7ZDB</u>, <u>7ZDC</u>, <u>7ZDE</u>, <u>7ZDF</u>, <u>7ZDG</u>, <u>7ZDK</u>, <u>7ZDL</u>, <u>7ZDR</u>, <u>7ZDS</u>, <u>7ZDT</u>, <u>7ZDU</u>, <u>7ZDV</u>, <u>7ZDW</u>, <u>7ZE5</u>, <u>7ZEC</u>. All other data is presented in the main text or supplementary materials. Source data are provided with this paper.

