## Supplementary materials for "Dissecting the conformational complexity and flipping mechanism of a prokaryotic heme transporter"

This PDF file includes:

Materials and Methods
Tables S1 to S3
Figures S1 to S25

Other Supplementary Material for this manuscript includes the following:
Movies S1-S13

### Materials and Methods

#### Production of CydDC from *Escherichia coli*

CydDC from *E. coli* was produced in *E. coli* BL21-pLysS (DE3) cells transformed with a pTTQ18 plasmid carrying structural genes of the CydDC heterodimer (*cydD + cydC)* and encoding a C-terminal hexahistidine modification at CydC for downstream IMAC purification (p*cydDC*) (*28*). Mutant variants $CydDC^{E500Q}$ (p*E500Q*), $CydD^{E511Q}C$ (p*E511Q*), and $CydDC^{H85A}$ (p*H85A*) were produced analogously. Further details are found in sequence data provided as supplementary information (tables S2 and S3).

For an initial pre-culture, a volume of 0.1 mL 50% glycerol stock of each respective strain was added to 200 mL M9-Carb (50 µg/mL carbenicillin) growth medium. Cells were incubated at 37 °C while shaking (185 rpm) for 16-20 h. The production culture was started by inoculating 2 L M9-Carb (50 µg/mL carbenicillin) growth medium with the pre-culture adjusted to a starting $OD_{600}$ of 0.1. Cells were grown while shaking (185 rpm) at 37 °C until an $OD_{600}$ of 0.6 before recombinant production was induced by addition of IPTG at a final concentration of 0.6 mM. Gene expression was carried out at 30 °C for 16-18 h. After harvest, cells were disrupted using a French-press cell disruptor (Thermo Fisher Scientific) via double-pass at a pressure of 700-1,000 psi. The cell lysate was centrifuged at 5,000x *g* at 4 °C for 30 minutes. Subsequently, the low-velocity supernatant was centrifuged at 220,000x *g* at 4 °C for 90 minutes. Pelleted membranes were resuspended and stored at a protein concentration of 10 mg/mL in a storage buffer containing 50 mM Tris-HCl (pH 7.4), 100 mM KCl.

#### Purification of CydDC from *Escherichia coli*

Isolated membranes were solubilized with 1% dodecyl-ß-D-maltoside (DDM) in a mass ratio of 1 mg detergent per 5 mg of membrane protein for 60 minutes at 4 °C. CydDC was purified via $His_6$-tag affinity chromatography using a Talon (Co-IMAC) affinity matrix (Takara Bio Inc.). For column conditioning and washing step I (10 column volumes [CV]) a buffer containing 50 mM Tris-HCl (pH 8), 500 mM KCl, 10% (v/v) glycerol, 20 mM imidazole, and 0.02% DDM was used. For samples intended to be used for cryo-EM studies, the resin was additionally washed with 10 CV of washing buffer II containing 50 mM Tris-HCl (pH 8), 500 mM KCl, 10% (v/v) glycerol, 20 mM imidazole, and 0.05% LMNG to perform detergent exchange from DDM to LMNG. For step elution (5 x 1 CV), buffers I and II were adjusted to a final imidazole concentration of 300 mM, respectively. For sample polishing, size exclusion chromatography (SEC) was performed using a Superdex 200 10/300 Increase column. The SEC running buffer contained 20 mM MES (pH 6), 50 mM KCl and either 0.02% DDM or 0.001% LMNG. All steps of the purification process were carried out at 4 °C.

**Growth complementation studies**

The bacterial growth assays were carried out as described previously (*30*). Pre-cultures of *E. coli* MB43 and MB43Δ*cydDC* (transformed with empty pET17 control vector or with plasmids encoding one of the cytochrome *bd* variants and/or CydDC) were incubated in Luria Bertani (LB) medium with 100 µg/mL ampicillin overnight at 37 °C while shaking at 200 rpm. Bacteria were then diluted to $OD_{600}$ of 0.01 with 200 µL LB medium containing 100 µg/mL ampicillin and each sample was subsequently distributed over 8 wells (technical replicates) of a 96-well microtiter plate. $OD_{600}$ was measured every 5 minutes at 37 °C for 20 hours (shaking at 200 rpm) in a SpectraMax Plus 384 Microplate reader (Molecular Devices).

**ATP hydrolysis assays**

*Malachite green phosphate assay*

ATPase activity of CydDC was determined colorimetrically as described previously (*62*). In brief, measurements were performed at a final concentration of 50 nM CydDC (in DDM) in a volume of 25 µL and carried out for 5 min at 37 °C. Reactions were stopped by adding 175 µL ice cold stopping buffer (20 mM $H_2SO_4$) and stored on ice. For detection, 175 µL of a stopped reaction sample was transferred into a 96 well microtiter plate and incubated for 8 min with 50 µL malachite green solution (2.7 mM Malachite Green chloride, 0.17% Tween, 1.5% $Na_2MoO_4$) at room temperature. The absorbance change at 620 nm was measured by a SpectraMax® M2 Microplate Reader (Molecular Deviced, U.S.A). For heme titration experiments a constant ATP concentration of 1 mM was used. ATP titration experiments were performed at a constant heme concentration of 0.3 µM. Substate screening experiments were performed using following final concentrations of the respective reaction components: ATP (1 mM), ferrous heme (0.5 µM), ferric heme (0.5 µM), protoporphyrin IX (0.5 µM), $FeCl_3$ (0.5 µM), GSH (1 mM), GSSG (1 mM), L-Cys (0.5 mM), CSSC (1 µM), orthovanadate (1 mM), AMP-PNP (1 mM). All measurements were performed in a reaction buffer containing 20 mM Tris-HCl (pH 7.0), 50 mM KCl, 0.02% DDM, and 3 mM $MgCl_2$. Samples without substrate were used as negative control and subtracted as background. All data are presented as mean ± SD (n = 3). Statistical significance was analyzed via paired two-tailed Student's t-tests using the GraphPad t-Test calculator (https://www.graphpad.com/quickcalcs/ttest1/).

*Enzyme-coupled ATPase assay*

The pyruvate kinase (PK)/lactate dehydrogenase (LDH) coupled ATP hydrolysis assay was performed as described previously (*63*). In brief, a volume of 100 µL activity buffer [20 mM Tris-HCl (pH 7.0), 50 mM KCl, 4 mM $MgCl_2$, 350 µM PEP, 350 µM ATP, 350 µM NADH, 0.02% DDM] was mixed with 2.5 µL PK/LDH enzyme solution (Sigma Aldrich - P0294) and incubated for 5 min at 37 °C in a 96-well microtiter plate. Subsequently, putative substrate candidates and CydDC (0.6 µM final concentration) were added. Kinetic turnover was monitored at 340 nm for 45 min every 10 s using the SpectraMax® M2 Microplate Reader (Molecular Devices, U.S.A). All data are presented as mean ± SD (n = 3). Statistical significance was analyzed via paired two-tailed Student's t-tests using the GraphPad t-Test calculator (https://www.graphpad.com/quickcalcs/ttest1/).

**Determination of thermal stability by microscale thermophoresis**

Thermal stabilities of purified CydDC variants were investigated with a Prometheus NT.48 instrument (Nanotemper). Excitation at 280 nm (20 nm bandwidth) was set to a power of 10% yielding emission intensities of 6,000 to 25,000 at 333-380 nm. A temperature ramp of 1 °C/min between 20 and 95 °C was applied in all experiments. Measurements were performed at 2 µM CydDC concentration. Heme was added in equimolar concentrations to CydDC. Unfolding transitions were monitored from changes in the emission of tryptophan and tyrosine fluorescence at 350/330 nm. Melting temperatures were determined at the inflection points (free energy change $\Delta G$ is equal to zero) from the raw data (measured in triplicates) after baseline correction and normalization.

**Cryo-EM sample preparation**

In order to collect cryo-EM data of CydDC in different sample conditions, different combinations of CydDC variants, nucleotides, inhibitors, and putative substrate molecules were prepared. See Fig. 2 and fig. S5 for details of specific sample conditions. For all non-turnover datasets, the protein concentration was adjusted to approximately 1.5 mg/mL before other components were added. Heme loading was performed prior to sample vitrification. Heme was adjusted to a final concentration of 17 µM in all heme containing samples. Nucleotides and inhibitors (ADP, AMP-PNP, and ATP-orthovanadate) were used at a final concentration of 1 mM (+ 2 mM $MgCl_2$). Putative substrate molecules were screened at final concentrations of 1 mM. Due to poor aqueous solubility cystine was adjusted to a final concentration of 0.2 mM. Samples were incubated for 2 minutes at room temperature before plunge freezing.

For preparation of turnover samples, the protein concentration was adjusted to 3 mg/mL. Subsequently, heme was added at a final concentration of 34 µM. The sample was then mixed with a freshly prepared ATPase buffer [20 mM MES (pH 6), 50 mM KCl, 0.001% LMNG, 10 mM ATP and 20 mM $MgCl_2$) in a ratio of 1:1, incubated for 30 seconds at 37 °C, and immediately subjected to plunge freezing. Identical plunge freezing conditions were applied for all samples: Quantifoil R1.2/1.3 copper grids (mesh 300) were washed in chloroform and subsequently glow discharged with a PELCO easiGlow device at 15 mA for 90 seconds. A volume of 4 µL sample was applied to a grid and blotting was performed for 4 seconds at 4 °C, 100% humidity with nominal blot force 20 immediately before freezing in liquid ethane, using a Vitrobot Mark IV device (Thermo Scientific).

**Cryo-EM image recording**

For each cryo-EM sample, a dataset was recorded in Energy-Filtered Transmission Electron Microscopy (EF-TEM) mode using either a Titan Krios G2 or a Krios G3i microscope (Thermo Scientific), both operated at 300 kV. Electron-optical alignments were adjusted with EPU 2.9 - 2.11 (Thermo Scientific). Images were recorded using automation strategies of EPU 2.9 - 2.11 in electron counting mode with either a Gatan K2 (installed on Krios G2) or a Gatan K3 (installed on Krios G3i) direct electron detector at a nominal magnification of 105,000, corresponding to a calibrated pixel size of 0.831 Å and 0.837 Å, respectively. Dose fractionated movies (40 frames) were recorded at an electron flux of approximately 15 $e^-$ x $pixel^{-1}$ x $s^{-1}$ for 2 s, corresponding to a total dose of ~40 $e^-/A^2$. Images were recorded between -1.1 and -2.1 µm nominal defocus.

**Cryo-EM image processing**

For each acquired dataset, the same cryo-EM image processing approach was applied: MotionCor2 was used to correct for beam-induced motion and to generate dose-weighted images (*64*). Gctf was used to determine the contrast transfer function (CTF) parameters and perform correction steps (*65*). Images with estimated poor resolution (>4 Å) and severe astigmatism (>400 Å) were removed at this step. Particles were picked by crYOLO and used for all further processing steps (*66*). 2D classification, initial model generation, 3D classification, CTF refinement, Bayesian polishing, 3D sorting, and final map reconstructions were performed using RELION-3.1 (*67*). Fourier shell correlation (FSC) curves were generated in RELION-3.1. Local-resolution estimation was performed in RELION-3.1 for all final maps. A schematic overview of our processing workflow, and a summary of map qualities are shown in figs. S4 to S6.

**Model building and geometry refinement**

The first atomic model of CydDC was built *de novo* into the EM density map of the $\mathbf{IF}^{\mathrm{apo}}_{\mathrm{as\ isolated}}$ state in Coot (version 0.8) (*68*). After manual backbone tracing and docking of side chains in the respective map densities real-space refinement in Phenix was performed (version 1.18) (*69*). Refinement results were manually inspected and corrected if required. This model was used as a template to build all subsequent atomic models. In total, 17 models were built, refined, inspected and corrected. The finalized models were validated by MolProbity implemented in Phenix (*70*). Map-to-model cross validation was performed in Phenix (version 1.18). $FSC_{0.5}$ was used as cut-off to define resolution. A summary of model parameters and the corresponding cryo-EM map statistics is found in table S1. The finalized models were visualized using ChimeraX (*71*). Highest resolution models of $\mathbf{IF}^{\mathrm{apo}}_{\mathrm{as\ isolated}}$, $\mathbf{IF}^{\mathrm{apo}}_{\mathrm{asym}}$, $\mathbf{IF}^{\mathrm{heme}}_{\mathrm{bound}}$, $\mathbf{IF}^{\mathrm{heme}}_{\mathrm{coordinated}}$, $\mathbf{IF}^{\mathrm{heme}}_{\mathrm{confined}}$, and $\mathbf{Occ}^{\mathrm{apo}}_{\mathrm{return}}$ states were used as starting structures for MD simulations.

**Tunnels and interior cavities**

Tunnels and cavities were mapped with MOLE 2.5 (bottleneck radius: 1.2 Å, bottleneck tolerance 3 Å, origin radius 5 Å, surface radius 10 Å, probe radius 5 Å, interior threshold 1.1 Å) (*72*).

**Structural alignments**

The overall folds of the CydDC subunits CydC and CydD were compared with each other using the structural alignment program TM-align (*73*). Subunits, individual transmembrane regions, and NBDs were aligned and respective Cα R.M.S.D values were calculated.

**Molecular dynamics simulations**

The CydDC structures were placed in a heterogenous bilayer composed of POPE (70%), POPG (25%), and CL (5%) using CHARMM-GUI (*74*). All systems were hydrated with 150 mM NaCl electrolyte. The all-atom CHARMM36m force field was used for lipids, ions, cofactors, and protein with TIP3P water (*75*, *76*). MD trajectories were analyzed using Visual Molecular Dynamics (VMD) (*77*) and MDAnalysis (*78*).

All simulations were performed using GROMACS 2021.2 (*79*). Starting systems were energy-minimized for 5,000 steepest descent steps and equilibrated initially for 500 ps of MD in a canonical (NVT) ensemble and later for 7.5 ns in an isothermal-isobaric (NPT) ensemble under periodic boundary conditions. During equilibration, the restraints on the positions of nonhydrogen protein atoms of initially 4,000 $\mathrm{kJ\cdot mol^{-1}\cdot nm^{2}}$ were gradually released. Particle-mesh Ewald summation (*80*) with cubic interpolation and a 0.12-nm grid spacing was used to treat long-range electrostatic interactions. The time step was initially 1 fs, and was increased to 2 fs during the NPT equilibration. The LINCS algorithm was used to fix all bond lengths (*81*). Constant temperature was established with a Berendsen thermostat, combined with a coupling constant of 1.0 ps. A semi-isotropic Berendsen barostat was used to maintain a pressure of 1 bar (*82*). During production runs, the Berendsen thermostat and barostat were replaced by a Nosé–Hoover thermostat and a Parrinello–Rahman barostat (*83*, *84*). All simulations were performed at 310 K. Analysis was carried out on unconstrained simulations. Simulations with liganded and unliganded heme were performed for 400 ns for each of $\mathbf{IF}^{\mathrm{heme}}_{\mathrm{bound}}$, $\mathbf{IF}^{\mathrm{heme}}_{\mathrm{coordinated}}$, and $\mathbf{IF}^{\mathrm{heme}}_{\mathrm{confined}}$ states.

For heme entry simulations a heterogenous bilayer composed of POPE (70%), POPG (25%), and CL(5%) was built using CHARMM-GUI (*74*). The system was hydrated with 150 mM NaCl electrolyte. After 7 ns of equilibration, a production run was performed for 100 ns. Then, a heme molecule (ferrous state) was placed 1 nm away from the membrane. Ten separate MD simulations were initiated with independent initial velocities drawn according to the Boltzmann distribution of the targeted temperature. Each replicate was run for additional 100 ns. In seven out of the ten simulations, the heme molecule had partitioned into the membrane bilayer at the end of 100ns simulation. The $\mathbf{IF}^{\mathrm{apo}}_{\mathrm{as\ isolated}}$ CydDC structure was placed in a heterogenous bilayer composed of POPE (70%), POPG (25%), and CL(5%) using CHARMM-GUI (*74*). Then, a heme molecule was placed near the lateral opening between TM4$^{D}$ and TM6$^{D}$ in 5 different positions. After equilibration, a simulation of 100 ns duration was performed for each setup.

We modeled the $\mathbf{Occ}^{\mathrm{heme}}_{\mathrm{confined}}$ state, which represents the heme-loaded conformation of CydDC upon ATP binding, with tightly interlocked NBDs. To obtain this conformation, we first ran an MD simulation for the fully occluded state of CydDC without heme ($\mathbf{Occ}^{\mathrm{apo}}_{\mathrm{return}}$) for 400 ns to get an equilibrated structure in the membrane bilayer. Then, the end structure of this simulation was aligned with the $\mathbf{IF}^{\mathrm{heme}}_{\mathrm{confined}}$ conformation and a virtual heme molecule was placed in the approximate position of the binding site taken from the aligned $\mathbf{IF}^{\mathrm{heme}}_{\mathrm{confined}}$ conformation. In the next step, three independent simulations were carried out to generate the $\mathbf{Occ}^{\mathrm{heme}}_{\mathrm{confined}}$ conformation by gradually turning on the interactions of the virtual heme with the protein and the rest of the system. Increasing a lambda parameter scaling these interactions slowly from 0 to 1, over 1 ns with a 0.5 fs timestep, results in the insertion of heme by slow growth. This method allows for smooth adaptation of amino acids in the binding site to the presence of heme. Extra position restraints were added to the C-alpha atoms of the NBDs to stabilize the conformation of protein during the slow-growth process. After the slow-growth heme insertion, we added the axial ligation bonds between heme and H85$^{C}$ and H312$^{D}$ and energy minimized the system. Then, we performed 300 ns of simulation. Structural stability during simulations was checked by RMSD evolution and conformation dynamics.

To obtain structural models of the two elusive outward-facing ($\mathbf{OF}^{\mathrm{heme}}_{\mathrm{confined}}$ and $\mathbf{OF}^{\mathrm{apo}}$) states, we performed steered MD simulations starting from $\mathbf{Occ}^{\mathrm{heme}}_{\mathrm{confined}}$ and $\mathbf{Occ}^{\mathrm{apo}}_{\mathrm{return}}$, respectively. A bias was applied on the distance between the center of mass of the two periplasmic halves of CydDC. The periplasmic halves are defined as (i) TM 1-2 of CydC and TM 3-6 of CydD and (ii) TM 1-2 of CydD and TM 3-6 of CydC. The initial value of the distance was 19 Å and the target value was 28 Å (based on the outward-open structure of TmrAB PBD ID 6RAJ). The steered MD was performed in PLUMED-patched

Gromacs with an approximate velocity of 0.01 nm/ns. The steered MD simulation was performed for 100 ns. After the simulation and full opening of the periplasmic side, we ran a further 50 ns of restrained simulation which allowed the lipid molecules around the periplasmic site to equilibrate around the new conformation. Later, we performed a further 200 ns of unrestrained simulation for each of the two conformations. Finally, we checked the stability and the degree of opening after 300 ns of unrestrained simulations.

**Preparation of membrane fractions for heme spectra and oxygen consumption assays**

Membrane fractions were isolated as described previously (*85*). In brief, *E. coli* strains were grown in 800 mL LB medium (2-liter baffled flasks) from a starting $OD_{600}$ of 0.01 until late exponential phase. After washing with phosphate buffer saline, batches of cells (5 g) were resuspended in a buffer composed of 50 mM MOPS (pH 7), 100 mM NaCl, and cOmplete protease inhibitor (Roche) in a ratio of 5:1. Cells were disrupted using a Stansted homogenizer at 1.2 kbar. Cell debris was removed centrifugation at 9,500× *g* for 20 min at 4 °C. Membrane fractions were collected by ultracentrifugation at 250,000× *g* for 75 min at 4 °C. Membranes were resuspended in the above buffer containing 0.025% n-dodecyl-β-D-maltoside (DDM), and used for downstream oxygen consumption activity assays and spectroscopic analysis of heme cofactors.

**Ultraviolet-visible (UV-Vis) absorption spectroscopy for heme identification**

The heme composition of membrane fractions was analyzed by reduced minus oxidized UV-Vis spectra based on Goojani *et al.* (*30*). Membrane fractions were diluted to a protein concentration of 2.6 mg/mL in a buffer composed of 10 mM Tris (pH 7.4) and 16 mM sodium cholate. Samples were first oxidized with 100 µM potassium ferricyanide and a spectrum was recorded at room temperature using a Varian Cary 50 UV-Vis Spectrophotometer. Subsequently, a few grains of solid sodium hydrosulfite were dissolved in the sample to measure the spectrum in the fully reduced state. The difference spectrum (reduced-oxidized) was calculated using Origin Lab Pro 9.5 (Additive GmbH, Germany).

**Oxygen consumption measurements**

The oxygen consumption activity of membrane fractions was measured using a Clark-type electrode based on Goojani *et al.* (*30*). The membrane fractions were adjusted to a concentration of 0.01 - 0.03 mg/mL using a buffer composed of 50 mM MOPS (pH 7.0), 100 mM NaCl, and 0.025% DDM. Subsequently, the membrane fractions were pre-incubated with either aurachin D (final concentration 400 nM) or with DMSO as control for 3 min in the electrode chamber. Ubiquinone-1 and dithiothreitol (DTT) were pre-incubated separately for 3 min and then injected into the electrode chamber to start the reaction (final conc.: 200 µM ubiquinone-1 and 10 mM DTT). The reaction rate was determined for the period between 90 s and 150 s after addition of the substrate mixture. Statistical significance was analyzed via paired two-tailed Student's t-tests using the GraphPad t-test calculator (https://www.graphpad.com/quickcalcs/ttest1/).

**Multiple sequence alignments**

Multiple sequence alignments of CydD and CydC from *Escherichia coli* (strain K12), *Mycobacterium tuberculosis* (strain ATCC 25618 / H37Rv), *Mycobacterium smegmatis* MC2-155, *Mycobacterium bovis* (strain ATCC BAA-935/AF2122/97), *Corynebacterium glutamicum*, *Brucella abortus biovar*, *Shewanella violacea* (strain JCM 10179/CIP 106290/LMG 19151/DSS12), *Geobacillus stearothermophilus*, and *Klebsiella pneumoniae* were performed using Clustal Omega (*86*) and visualized using Jalview v. 2.11.2.0.

**Table S1 – Cryo-EM and model data statistics.**

| | **Dataset 1** IF(apo/as isolated) | **Dataset 1** IF(heme/confined) | **Dataset 2** IF(apo/asym) | **Dataset 2** IF(heme/bound) | **Dataset 3** IF(apo/as isolated) | **Dataset 3** IF(heme/confined) | **Dataset 4** IF(apo/as isolated) | **Dataset 4** IF(heme/confined) | **Dataset 5** IF(heme/confined) |
|---|---|---|---|---|---|---|---|---|---|
| **Data collection** | | | | | | | | | |
| Accession number | EMDB-14636 | EMDB-14638 | EMDB-14639 | EMDB-14640 | EMDB-14641 | EMDB-14642 | EMDB-14643 | EMDB-14644 | EMDB-14645 |
| Magnification | 105,000 | 105,000 | 105,000 | 105,000 | 105,000 | 105,000 | 105,000 | 105,000 | 105,000 |
| Voltage / kV | 300 | 300 | 300 | 300 | 300 | 300 | 300 | 300 | 300 |
| Dose / $e^{-}Å^{-2}$ | 42 | 42 | 43 | 43 | 42 | 42 | 41 | 41 | 41 |
| Pixel size / Å | 0.837 | 0.837 | 0.837 | 0.837 | 0.837 | 0.837 | 0.837 | 0.837 | 0.837 |
| Defocus range / µm | -1.1 to -2.1 | -1.1 to -2.1 | -1.1 to -2.1 | -1.1 to -2.1 | -1.1 to -2.1 | -1.1 to -2.1 | -1.1 to -2.1 | -1.1 to -2.1 | -1.1 to -2.1 |
| Recorded movies | 7,952 | 7,952 | 24,110 | 24,110 | 7,508 | 7,508 | 6,072 | 6,072 | 7,814 |
| Final particle images | 98,317 | 86,512 | 39592 | 87,939 | 71,865 | 89,336 | 57,308 | 89,870 | 644,226 |
| Camera | Gatan K3 | Gatan K3 | Gatan K3 | Gatan K3 | Gatan K3 | Gatan K3 | Gatan K3 | Gatan K3 | Gatan K3 |
| Energy filter | BioQuantum K3 | BioQuantum K3 | BioQuantum K3 | BioQuantum K3 | BioQuantum K3 | BioQuantum K3 | BioQuantum K3 | BioQuantum K3 | BioQuantum K3 |
| Microscope | Titan Krios G3i | Titan Krios G3i | Titan Krios G3i | Titan Krios G3i | Titan Krios G3i | Titan Krios G3i | Titan Krios G3i | Titan Krios G3i | Titan Krios G3i |
| **Image processing** | | | | | | | | | |
| Initial model | De novo generated with RELION 3.1 | | | | | | | | |
| Resolution ($FSC_{0.143}$) / Å | 3.17 | 3.05 | 3.17 | 3.35 | 3.65 | 3.13 | 3.17 | 2.94 | 2.77 |
| Applied B-factor / $Å^2$ | -73 | -64 | -57 | -75 | -102 | -80 | -59 | -58 | -70 |
| **Model refinement** | | | | | | | | | |
| PDB accession | 7ZD5 | - | 7ZDA | 7ZDB | - | 7ZDC | 7ZDE | 7ZDF | 7ZDG |
| Validation | | | | | | | | | |
| $FSC^{map\text{-}to\text{-}model}_{(0.5)}$ / Å | 3.1 | - | 3.2 | 3.3 | - | 3.1 | 3.0 | 2.9 | 2.7 |
| MolProbity score | 1.49 | - | 1.47 | 1.58 | - | 1.45 | 1.70 | 1.46 | 1.54 |
| Composition | | | | | | | | | |
| Atoms | 8,968 | - | 9,052 | 8,804 | - | 8,961 | 9,015 | 9,039 | 9,007 |
| Protein residues | 1,157 | - | 1,159 | 1,122 | - | 1,147 | 1,159 | 1,159 | 1,157 |
| Ligands | - | - | MG: 2, ADP: 1, PO3: 1, ATP: 1 | HEB: 1, MG: 2, ADP:1, PO3: 1, ATP: 1 | - | HEB: 1, MG: 1, ADP: 1 | MG: 1, ANP: 1 | HEB: 1, MG: 1, ANP: 1 | HEB: 1 |
| Bonds (R.M.S.D.) | | | | | | | | | |
| Length (Å) | 0.006 | - | 0.006 | 0.004 | - | 0.005 | 0.007 | 0.004 | 0.008 |
| Angles (°) | 0.709 | - | 0.706 | 0.740 | - | 0.727 | 0.770 | 0.651 | 0.811 |
| B-factors (min/max/mean) | | | | | | | | | |
| Protein | 13.98/105.03/46.35 | - | 3.35/79.70/25.61 | 14.56/96.07/47.96 | - | 8.65/79.13/33.20 | 15.71/126.14/51.59 | 13.99/81.56/38.33 | 12.90/93.23/48.08 |
| Ligand | - | - | 23.80/42.74/34.93 | 46.02/82.62/62.76 | - | 17.11/32.25/22.99 | 20.00/72.06/21.63 | 28.94/36.65/34.86 | 41.36/41.36/41.36 |
| Clash score | 9.26 | - | 8.71 | 11.21 | - | 7.57 | 10.71 | 7.61 | 9.39 |
| Ramachandran plot (%) | | | | | | | | | |
| Favored | 98.26 | - | 98.27 | 97.94 | - | 97.90 | 97.14 | 97.84 | 97.83 |
| Allowed | 1.74 | - | 1.73 | 2.06 | - | 2.10 | 2.86 | 2.16 | 2.17 |
| Outliers | 0 | - | 0 | 0 | - | 0 | 0 | 0 | 0 |
| Rotamer outliers (%) | 0.32 | - | 0.11 | 0 | - | 0 | 0 | 0 | 0 |

**Table S1 –** Continued

| | **Dataset 6** IF(heme/confined) | **Dataset 7** IF(apo/asym) | **Dataset 7** IF(heme/coordinated) | **Dataset 8** IF(apo/asym) | **Dataset8** IF(heme/coordinated) | **Dataset 9** IF(heme/confined) | **Dataset 10** IF(apo/as isolated) | **Dataset 10** IF(heme/confined) | **Dataset 11** IF(apo/as isolated) |
|---|---|---|---|---|---|---|---|---|---|
| **Data collection** | | | | | | | | | |
| Accession number | EMDB-14646 | EMDB-14647 | EMDB-14649 | EMDB-14652 | EMDB-14653 | EMDB-14654 | EMDB-14655 | EMDB-14656 | EMDB-14657 |
| Magnification | 105,000 | 105,000 | 105,000 | 105,000 | 105,000 | 105,000 | 105,000 | 105,000 | 105,000 |
| Voltage / kV | 300 | 300 | 300 | 300 | 300 | 300 | 300 | 300 | 300 |
| Dose / $e^{-}Å^{-2}$ | 41 | 41 | 41 | 41 | 41 | 48 | 41 | 41 | 41 |
| Pixel size / Å | 0.837 | 0.837 | 0.837 | 0.837 | 0.837 | 0.831 | 0.837 | 0.837 | 0.837 |
| Defocus range / µm | -1.1 to -2.1 | -1.1 to -2.1 | -1.1 to -2.1 | -1.1 to -2.1 | -1.1 to -2.1 | -1.1 to -2.1 | -1.1 to -2.1 | -1.1 to -2.1 | -1.1 to -2.1 |
| Recorded movies | 8,538 | 9,920 | 9,920 | 7,552 | 7,552 | 13,299 | 7,436 | 7,436 | 8,392 |
| Final particle images | 132,480 | 73,885 | 47,521 | 96,066 | 120,630 | 152,983 | 81,388 | 72,194 | 99,304 |
| Camera | Gatan K3 | Gatan K3 | Gatan K3 | Gatan K3 | Gatan K3 | Gatan K2 Summit | Gatan K3 | Gatan K3 | Gatan K3 |
| Energy filter | BioQuantum K3 | BioQuantum K3 | BioQuantum K3 | BioQuantum K3 | BioQuantum K3 | Quantum K2 | BioQuantum K3 | BioQuantum K3 | BioQuantum K3 |
| Microscope | Titan Krios G3i | Titan Krios G3i | Titan Krios G3i | Titan Krios G3i | Titan Krios G3i | Titan Krios G2 | Titan Krios G3i | Titan Krios G3i | Titan Krios G3i |
| **Image processing** | | | | | | | | | |
| Initial model | De novo generated with RELION 3.1 | | | | | | | | |
| Resolution ($FSC_{0.143}$) / Å | 3.09 | 3.49 | 3.49 | 3.01 | 3.35 | 3.42 | 3.26 | 3.35 | 3.44 |
| Applied B-factor / $Å^2$ | -72 | -73 | -88 | -68 | -89 | -98 | -88 | -70 | -95 |
| **Model refinement** | | | | | | | | | |
| PDB accession | - | - | - | 7ZDK | 7ZDL | - | - | - | - |
| Validation | | | | | | | | | |
| $FSC^{map\text{-}to\text{-}model}_{(0.5)}$ / Å | - | - | - | 3.0 | 3.3 | - | - | - | - |
| MolProbity score | - | - | - | 1.45 | 1.80 | - | - | - | - |
| Composition | | | | | | | | | |
| Atoms | - | - | - | 9,089 | 9,026 | - | - | - | - |
| Protein residues | - | - | - | 1,161 | 1,152 | - | - | - | - |
| Ligands | - | - | - | MG: 2, ANP:2 | HEB: 1, MG: 2, ANP: 2 | - | - | - | - |
| Bonds (R.M.S.D.) | | | | | | | | | |
| Length (Å) | - | - | - | 0.006 | 0.007 | - | - | - | - |
| Angles (°) | - | - | - | 0.775 | 0.844 | - | - | - | - |
| B-factors (min/max/mean) | | | | | | | | | |
| Protein | - | - | - | 8.26/63.36/24.44 | 19.33/101.69/48.81 | - | - | - | - |
| Ligand | - | - | - | 21.12/33.68/29.60 | 37.77/73.77/54.51 | - | - | - | - |
| Clash score | - | - | - | 8.29 | 11.54 | - | - | - | - |
| Ramachandran plot (%) | | | | | | | | | |
| Favored | - | - | - | 98.27 | 96.60 | - | - | - | - |
| Allowed | - | - | - | 1.73 | 3.40 | - | - | - | - |
| Outliers | - | - | - | 0 | 0 | - | - | - | - |
| Rotamer outliers (%) | - | - | - | 0 | 0 | - | - | - | - |

**Table S1** – Continued

| | **Dataset 12** IF(heme/confined) | **Dataset 13** IF(apo/asym) | **Dataset 13** IF(heme/coordinated) | **Dataset 14** IF(heme/coordinated) | **Dataset 14** IF(heme/confined) | **Dataset 15** IF(heme/confined) | **Dataset 16** IF(apo/as isolated) | **Dataset 17** IF(apo/as isolated) | **Dataset 18** IF(apo/as isolated) |
|---|---|---|---|---|---|---|---|---|---|
| **Data collection** | | | | | | | | | |
| Accession number | EMDB-14659 | EMDB-14660 | EMDB-14662 | EMDB-14663 | EMDB-14665 | EMDB-14689 | EMDB-14667 | EMDB-15264 | EMDB-14668 |
| Magnification | 105,000 | 105,000 | 105,000 | 105,000 | 105,000 | 105,000 | 105,000 | 105,000 | 105,000 |
| Voltage / kV | 300 | 300 | 300 | 300 | 300 | 300 | 300 | 300 | 300 |
| Dose / $e^{-}Å^{-2}$ | 41 | 41 | 41 | 41 | 41 | 41 | 41 | 41 | 41 |
| Pixel size / Å | 0.837 | 0.837 | 0.837 | 0.837 | 0.837 | 0.837 | 0.837 | 0.837 | 0.837 |
| Defocus range / µm | -1.1 to -2.1 | -1.1 to -2.1 | -1.1 to -2.1 | -1.1 to -2.1 | -1.1 to -2.1 | -1.1 to -2.1 | -1.1 to -2.1 | -1.1 to -2.1 | -1.1 to -2.1 |
| Recorded movies | 8,936 | 10,092 | 10,092 | 9,576 | 9,576 | 8,009 | 5,372 | 9138 | 7,646 |
| Final particle images | 87,143 | 148,055 | 81,009 | 231,551 | 131,737 | 83,501 | 107,130 | 204256 | 88,567 |
| Camera | Gatan K3 | Gatan K3 | Gatan K3 | Gatan K3 | Gatan K3 | Gatan K3 | Gatan K3 | Gatan K3 | Gatan K3 |
| Energy filter | BioQuantum K3 | BioQuantum K3 | BioQuantum K3 | BioQuantum K3 | BioQuantum K3 | BioQuantum K3 | BioQuantum K3 | BioQuantum K3 | BioQuantum K3 |
| Microscope | Titan Krios G3i | Titan Krios G3i | Titan Krios G3i | Titan Krios G3i | Titan Krios G3i | Titan Krios G3i | Titan Krios G3i | Titan Krios G3i | Titan Krios G3i |
| **Image processing** | | | | | | | | | |
| Initial model | De novo generated with RELION 3.1 | | | | | | | | |
| Resolution ($FSC_{0.143}$) / Å | 3.44 | 3.26 | 3.17 | 3.05 | 2.87 | 3.05 | 3.05 | 3.26 | 3.26 |
| Applied B-factor / $Å^2$ | -91 | -80 | -81 | -75 | -60 | -71 | -65 | -86 | -86 |
| **Model refinement** | | | | | | | | | |
| PDB accession | - | - | - | - | - | 7ZEC | 7ZDR | - | 7ZDS |
| Validation | | | | | | | | | |
| $FSC^{map\text{-}to\text{-}model}_{(0.5)}$ / Å | - | - | - | - | - | 3.0 | 3.0 | - | 3.2 |
| MolProbity score | - | - | - | - | - | 1.55 | 1.44 | - | 1.56 |
| Composition | | | | | | | | | |
| Atoms | - | - | - | - | - | 8,997 | 9,014 | - | 8,737 |
| Protein residues | - | - | - | - | - | 1,149 | 1,158 | - | 1,129 |
| Ligands | - | - | - | - | - | HEB:1, MG: 1, ATP: 1 | MG: 1, ANP: 1 | - | - |
| Bonds (R.M.S.D.) | | | | | | | | | |
| Length (Å) | - | - | - | - | - | 0.008 | 0.006 | - | 0.005 |
| Angles (°) | - | - | - | - | - | 0.800 | 0.641 | - | 0.724 |
| B-factors (min/max/mean) | | | | | | | | | |
| Protein | - | - | - | - | - | 11.45/83.58/39.04 | 16.22/100.95/42.68 | - | 7.26/96.17/38.30 |
| Ligand | - | - | - | - | - | 14.14/46.22/27.73 | 54.79/62.82/42.68 | - | - |
| Clash score | - | - | - | - | - | 8.15 | 8.13 | - | 10.98 |
| Ramachandran plot (%) | | | | | | | | | |
| Favored | - | - | - | - | - | 97.46 | 98.09 | - | 98.13 |
| Allowed | - | - | - | - | - | 2.54 | 1.91 | - | 1.87 |
| Outliers | - | - | - | - | - | 0 | 0 | - | 0 |
| Rotamer outliers (%) | - | - | - | - | - | 0 | 0 | - | 0 |

**Table S1** – Continued

| | **Dataset 18** Occ(apo/return) | **Dataset 19** Occ(apo/return) | **Dataset 20** IF(apo/as isolated) | **Dataset 20** Occ(apo/return) | **Dataset 21** IF(heme/confined) | **Dataset 21** Occ(apo/return) | **Dataset 22** IF(heme/confined) | **Dataset 22** Occ(apo/return) | **Dataset 23** IF(heme/coordinated) |
|---|---|---|---|---|---|---|---|---|---|
| **Data collection** | | | | | | | | | |
| Accession number | EMDB-14669 | EMDB-14670 | EMDB-14671 | EMDB-14672 | EMDB-14673 | EMDB-14674 | EMDB-14675 | EMDB-14676 | EMDB-15265 |
| Magnification | 105,000 | 105,000 | 105,000 | 105,000 | 105,000 | 105,000 | 105,000 | 105,000 | 105,000 |
| Voltage / kV | 300 | 300 | 300 | 300 | 300 | 300 | 300 | 300 | 300 |
| Dose / $e^{-}Å^{-2}$ | 41 | 41 | 41 | 41 | 41 | 41 | 41 | 41 | 41 |
| Pixel size / Å | 0.837 | 0.837 | 0.837 | 0.837 | 0.837 | 0.837 | 0.837 | 0.837 | 0.837 |
| Defocus range / µm | -1.1 to -2.1 | -1.1 to -2.1 | -1.1 to -2.1 | -1.1 to -2.1 | -1.1 to -2.1 | -1.1 to -2.1 | -1.1 to -2.1 | -1.1 to -2.1 | -1.1 to -2.1 |
| Recorded movies | 7,646 | 15,860 | 8,856 | 8,856 | 8,090 | 8,090 | 2,294 | 2,294 | 8,270 |
| Final particle images | 96,900 | 102,009 | 114,003 | 132,215 | 77,061 | 130,996 | 44,248 | 44,418 | 50,573 |
| Camera | Gatan K3 | Gatan K3 | Gatan K3 | Gatan K3 | Gatan K3 | Gatan K2 Summit | Gatan K3 | Gatan K3 | Gatan K3 |
| Energy filter | BioQuantum K3 | BioQuantum K3 | BioQuantum K3 | BioQuantum K3 | BioQuantum K3 | Quantum K2 | BioQuantum K3 | BioQuantum K3 | BioQuantum K3 |
| Microscope | Titan Krios G3i | Titan Krios G3i | Titan Krios G3i | Titan Krios G3i | Titan Krios G3i | Titan Krios G2 | Titan Krios G3i | Titan Krios G3i | Titan Krios G3i |
| **Image processing** | | | | | | | | | |
| Initial model | De novo generated with RELION 3.1 | | | | | | | | |
| Resolution ($FSC_{0.143}$) / Å | 2.71 | 2.98 | 3.05 | 3.89 | 3.77 | 3.17 | 3.35 | 3.35 | 3.65 |
| Applied B-factor / $Å^2$ | -48 | -46 | -74 | -114 | -128 | -70 | -55 | -61 | -92 |
| **Model refinement** | | | | | | | | | |
| PDB accession | 7ZDT | 7ZDU | 7ZDV | - | - | - | 7ZDW | - | - |
| Validation | | | | - | | | | | - |
| $FSC^{map\text{-}to\text{-}model}_{(0.5)}$ / Å | 2.7 | 3.0 | 3.0 | | - | - | 3.3 | - | |
| MolProbity score | 1.35 | 1.45 | 1.50 | - | - | - | 1.72 | - | - |
| Composition | | | | - | | | | | - |
| Atoms | 9,004 | 8,975 | 8,878 | - | - | - | 9,039 | - | - |
| Protein residues | 1,155 | 1,148 | 1,141 | | - | - | 1,159 | - | |
| Ligands | MG: 1, ATP: 1 | MG: 2, ATP: 2 | MG: 1, ANP: 1 | - | - | - | HEB: 1, MG: 1, ANP: 1 | - | - |
| Bonds (R.M.S.D.) | | | | - | | | | | - |
| Length (Å) | 0.005 | 0.004 | 0.005 | | - | - | 0.007 | - | |
| Angles (°) | 0.701 | 0.626 | 0.659 | - | - | - | 0.818 | - | - |
| B-factors (min/max/mean) | | | | - | | | | | - |
| Protein | 11.59/83.41/34.04 | 22.88/110.11/51.52 | 9.90/100.91/44.51 | - | - | - | 14.73/100.56/48.49 | - | - |
| Ligand | 24.76/27.64/27.55 | 42.80/48.91/45.86 | 66.48/69.69/69.59 | | - | - | 24.16/61.09/39.79 | - | |
| Clash score | 6.21 | 8.23 | 8.86 | - | - | - | 9.91 | - | - |
| Ramachandran plot (%) | | | | - | | | | | - |
| Favored | 98.70 | 98.25 | 97.88 | - | - | - | 96.80 | - | - |
| Allowed | 1.30 | 1.75 | 2.12 | - | - | - | 3.20 | - | - |
| Outliers | 0 | 0 | 0 | - | - | - | 0 | - | - |
| Rotamer outliers (%) | 0 | 0 | 0.11 | - | - | - | 0 | - | - |

**Table S1** – Continued

| | **Dataset 23**<br>Occ(apo/return) |
|---|---|
| **Data collection** | |
| Accession number | EMDB-14684 |
| Magnification | 105,000 |
| Voltage / kV | 300 |
| Dose / $e^{-}$Å$^{-2}$ | 41 |
| Pixel size / Å | 0.837 |
| Defocus range / µm | -1.1 to -2.1 |
| Recorded movies | 8,270 |
| Final particle images | 74,444 |
| Camera | Gatan K3 |
| Energy filter | BioQuantum K3 |
| Microscope | Titan Krios G3i |
| **Image processing** | |
| Initial model | De novo generated with RELION 3.1 |
| Resolution ($FSC_{0.143}$) / Å | 2.94 |
| Applied B-factor / Å$^2$ | -48 |
| **Model refinement** | |
| PDB accession | 7ZE5 |
| Validation | |
| $FSC^{map\text{-}to\text{-}model}{}_{(0.5)}$ / Å | 2.9 |
| MolProbity score | 1.42 |
| Composition | |
| Atoms | 9,077 |
| Protein residues | 1,159 |
| Ligands | MG: 2, ATP:1, ANP: 1 |
| Bonds (R.M.S.D.) | |
| Length (Å) | 0.004 |
| Angles (°) | 0.589 |
| B-factors (min/max/mean) | |
| Protein | 26.14/94.30/48.12 |
| Ligand | 37.64/42.78/40.93 |
| Clash score | 7.65 |
| Ramachandran plot (%) | |
| Favored | 98.27 |
| Allowed | 1.73 |
| Outliers | 0 |
| Rotamer outliers (%) | 0 |

**Table S2 – Nucleotide sequences of CydDC variants used in this study.**

| Variant | Nucleotide sequence |
|---|---|
| CydDC (wild-type) | atgaattcgaataaatctcgtcaaaaagagttaacccgctggttaaaacagcaaagcgtcatctcccaacgttggctgaatatttctcgtctgctgggctttgtgag<br>cggcatattgatcattgcccaggcctggttcatggcgcgtattctgcaacatatgattatggagaatattccccgtgaagccctgctgcttccctttacgttactggtt<br>ctgacctttgtactgcgcgcatgggtggtctggttacgcgaacgggtgggttatcacgccgggcagcatatccgctttgccatccgccgtcaggttctcgaccgtct<br>gcaacaagcagggccagcgtggattcagggtaaacctgcggggagctgggcgacgctggtactcgagcaaattgacgatatgcatgattactatgcacgctatc<br>tgccgcaaatggcgctggcagtgtcggtgccgttgctgattgtggtggcaatcttcccctctaactgggctgcggcgctcattctgctgggcactgcaccgttaattc<br>cgttgtttatggcgctggttggaatgggggctgccgatgctaaccgacgtaactttctcgctcttgctcgcttaagtgggcatttcctcgatcgcctgcgcggcatgg<br>aaacattgcgtatttttggtcgtggtgaagctgaaattgaaagtattcgttctgcttcggaagatttccgccaacggacaatggaagtgctacggctggcgtttttat<br>cctccggcattctcgaatttttacctcgctgtcaattgctctggtggcggtctactttggtttttcctatctcggcgagctggattttggtcactacgataccggtgtga<br>cgctggctgcgggttttctggccctgatccttgcgccagagtttttccagccattacgcgatctcggtacgttttatcatgctaaagcccaggctgttggcgcagctg<br>acagtctgaaaacgtttatggaaaccccgctcgcccatccgcaacgtggtgaggcggaattagcatcgaccgatccggtgaccattgaggccgaggagctgttta<br>tcacgtcgccggaaggtaaaacgctggccggaccgctgaactttactttgccagcaggccaacgtgcggtgttggttggtcgcagcggttcaggtaaaagctcac<br>tgctgaacgcgctttctggttttctctcatatcagggatcgctacgaatcaacgggatagaattacgcgatttatcaccagaatcatggcgtaaacatctctcctcgg<br>ttgggcaaaacccacaattaccggcagcaacattgcgggataacgtactactggcgcgacctgatgccagcgaacaagaattacaagcagcgctggataacgc<br>ctgggtcagcgagtttctaccgctcctcccacaaggcgttgatacgcctgttggcgaccaggctgcccgcctttccgtggggcaggcgcagcgcgtggcggtggcc<br>cgtgcgttactaaatccctgttcgctattactgttggatgaacccgctgccagccttgatgctcacagtgaacagcgcgtaatggaggcgctgaatgccgcctctct<br>gcgccagacaacgttaatggtcacccaccagttagaagatcttgctgactgggatgtcatttgggttatgcaggatggccggattattgagcaaggacgttacgcg<br>gaattaagtgtggctggtggcccattcgccacattactggcccatcgtcaggaggagatttaaatgcgcgctttgctaccctatctggcactgtataaacgtcataa<br>atggatgttaagtcttggtattgtgctggcaattgtgacgctgctcgccagtatcggtctgttgacactttccggctggttcctctcggcctcagcggttgcggggtt<br>gccggactgtacagcttcaactatatgctacccgctgcgggcgtgcgtggcgcagcaatcacccgtactgccgggcgctattttgaacgtctggtaagtcacgacg<br>cgactttccgcgtgttgcagcatctgcgcatttacaccttcagcaaattgctgcccctctcccctgccggactggcgcgctatcgtcagggcgaattgctcaatcgcg<br>tggtggcggatgttgatacgctcgatcatctttacctgcgcgttatctcgccgctggtgggcgcttttgtggtgattatggtggtgacaatcgggttaagtttccttga<br>tttcaccctcgcctttacgctgggcggcattatgttactgacgcttttcctgatgccaccgctgttttatcgtgcgggaaaaagcaccgggcaaaatctgactcatctt<br>cgcggacagtatcgccaacaactgacggcctggctgcaagggcaagctgagctgaccatttttggtgccagcgatcgttatcgcacgcaactagagaatacaga<br>aattcaatggctggaagcgcaacgccgtcaatctgaactgaccgcattgtcgcaagcgataatgctgctcattggcgcgttagcggtgatcctgatgctgtggatg<br>gcgtctggcggcgttggcggcaatgctcaacccggcgcgttaattgccctgtttgtcttctgcgcgttagccgcgtttgaagcactggcaccagtaacgggtgcatt<br>tcagcatctggggcaagtcattgcctctgccgtacgtatctctgacttaacggatcaaaaaccggaggtcacctttcctgatacccaaactcgtgttgccgatcgcg<br>tttcgctgacgttacgggatgttcagttcacttatccggagcaatctcaacaggcacttaaagggatttctcttcaggtaaacgccggggaacatatagcgattctc<br>gggcgaaccggatgcggcaaatcaacactgttacaacagctgacccgcgcatgggacccgcaacagggcgagattttgcttaacgatagccccatagccagcct<br>gaatgaagcggctctacgacagaccatcagcgttgttcctcagcgagtgcatctgtttagcgccacgctgcgtgataatcttttactcgcctcgcctggcagtagtg<br>atgaggctctgtcggagatcttgcgtcgcgttggcctggaaaagctgctcgaggatgcaggtctcaacagttggttaggtgaaggcggacgccagctctccggtg<br>gtgaactgcgccgtctggctatcgcccgtgcgctgttacatgatgcgccactggtgttgctggatgaacctaccgaaggcttagatgccacaaccgaaagccagat<br>ccttgaattgcttgcagaaatgatgcgtgagaaaacggtgttaatggtcacccatcgacttcgcggactctctcgtttccaacaaataatagtgatggacaacggg<br>caaattattgagcaaggtactcacgcagaactgcttgccagacaggggcgttattaccagttcaagcagggtttgtctgcaggcggtcgtggcagccaccatcac<br>catcaccattaa |
| CydD$^{E511Q}$C | atgaattcgaataaatctcgtcaaaaagagttaacccgctggttaaaacagcaaagcgtcatctcccaacgttggctgaatatttctcgtctgctgggctttgtgag<br>cggcatattgatcattgcccaggcctggttcatggcgcgtattctgcaacatatgattatggagaatattccccgtgaagccctgctgcttccctttacgttactggtt<br>ctgacctttgtactgcgcgcatgggtggtctggttacgcgaacgggtgggttatcacgccgggcagcatatccgctttgccatccgccgtcaggttctcgaccgtct<br>gcaacaagcagggccagcgtggattcagggtaaacctgcggggagctgggcgacgctggtactcgagcaaattgacgatatgcatgattactatgcacgctatc<br>tgccgcaaatggcgctggcagtgtcggtgccgttgctgattgtggtggcaatcttcccctctaactgggctgcggcgctcattctgctgggcactgcaccgttaattc<br>cgttgtttatggcgctggttggaatgggggctgccgatgctaaccgacgtaactttctcgctcttgctcgcttaagtgggcatttcctcgatcgcctgcgcggcatgg<br>aaacattgcgtatttttggtcgtggtgaagctgaaattgaaagtattcgttctgcttcggaagatttccgccaacggacaatggaagtgctacggctggcgtttttat<br>cctccggcattctcgaatttttacctcgctgtcaattgctctggtggcggtctactttggtttttcctatctcggcgagctggattttggtcactacgataccggtgtga<br>cgctggctgcgggttttctggccctgatccttgcgccagagtttttccagccattacgcgatctcggtacgttttatcatgctaaagcccaggctgttggcgcagctg<br>acagtctgaaaacgtttatggaaaccccgctcgcccatccgcaacgtggtgaggcggaattagcatcgaccgatccggtgaccattgaggccgaggagctgttta<br>tcacgtcgccggaaggtaaaacgctggccggaccgctgaactttactttgccagcaggccaacgtgcggtgttggttggtcgcagcggttcaggtaaaagctcac<br>tgctgaacgcgctttctggttttctctcatatcagggatcgctacgaatcaacgggatagaattacgcgatttatcaccagaatcatggcgtaaacatctctcctcgg<br>ttgggcaaaacccacaattaccggcagcaacattgcgggataacgtactactggcgcgacctgatgccagcgaacaagaattacaagcagcgctggataacgc<br>ctgggtcagcgagtttctaccgctcctcccacaaggcgttgatacgcctgttggcgaccaggctgcccgcctttccgtggggcaggcgcagcgcgtggcggtggcc<br>cgtgcgttactaaatccctgttcgctattactgttggatCAGcccgctgccagccttgatgctcacagtgaacagcgcgtaatggaggcgctgaatgccgcctctc<br>tgcgccagacaacgttaatggtcacccaccagttagaagatcttgctgactgggatgtcatttgggttatgcaggatggccggattattgagcaaggacgttacgc<br>ggaattaagtgtggctggtggcccattcgccacattactggcccatcgtcaggaggagatttaaatgcgcgctttgctaccctatctggcactgtataaacgtcata<br>aatggatgttaagtcttggtattgtgctggcaattgtgacgctgctcgccagtatcggtctgttgacactttccggctggttcctctcggcctcagcggttgcggggt<br>tgccggactgtacagcttcaactatatgctacccgctgcgggcgtgcgtggcgcagcaatcacccgtactgccgggcgctattttgaacgtctggtaagtcacgac<br>gcgactttccgcgtgttgcagcatctgcgcatttacaccttcagcaaattgctgcccctctcccctgccggactggcgcgctatcgtcagggcgaattgctcaatcgc<br>gtggtggcggatgttgatacgctcgatcatctttacctgcgcgttatctcgccgctggtgggcgcttttgtggtgattatggtggtgacaatcgggttaagtttccttg<br>atttcaccctcgcctttacgctgggcggcattatgttactgacgcttttcctgatgccaccgctgttttatcgtgcgggaaaaagcaccgggcaaaatctgactcatc<br>ttcgcggacagtatcgccaacaactgacggcctggctgcaagggcaagctgagctgaccatttttggtgccagcgatcgttatcgcacgcaactagagaatacag<br>aaattcaatggctggaagcgcaacgccgtcaatctgaactgaccgcattgtcgcaagcgataatgctgctcattggcgcgttagcggtgatcctgatgctgtggat<br>ggcgtctggcggcgttggcggcaatgctcaacccggcgcgttaattgccctgtttgtcttctgcgcgttagccgcgtttgaagcactggcaccagtaacgggtgcat<br>ttcagcatctggggcaagtcattgcctctgccgtacgtatctctgacttaacggatcaaaaaccggaggtcacctttcctgatacccaaactcgtgttgccgatcgc<br>gtttcgctgacgttacgggatgttcagttcacttatccggagcaatctcaacaggcacttaaagggatttctcttcaggtaaacgccggggaacatatagcgattct<br>cgggcgaaccggatgcggcaaatcaacactgttacaacagctgacccgcgcatgggacccgcaacagggcgagattttgcttaacgatagccccatagccagc<br>ctgaatgaagcggctctacgacagaccatcagcgttgttcctcagcgagtgcatctgtttagcgccacgctgcgtgataatcttttactcgcctcgcctggcagtag<br>tgatgaggctctgtcggagatcttgcgtcgcgttggcctggaaaagctgctcgaggatgcaggtctcaacagttggttaggtgaaggcggacgccagctctccggt<br>ggtgaactgcgccgtctggctatcgcccgtgcgctgttacatgatgcgccactggtgttgctggatgaacctaccgaaggcttagatgccacaaccgaaagccag<br>atccttgaattgcttgcagaaatgatgcgtgagaaaacggtgttaatggtcacccatcgacttcgcggactctctcgtttccaacaaataatagtgatggacaacg<br>ggcaaattattgagcaaggtactcacgcagaactgcttgccagacaggggcgttattaccagttcaagcagggtttgtctgcaggcggtcgtggcagccaccatc<br>accatcaccattaa |

| | |
|---|---|
| CydDC$^{E500Q}$ | atgaattcgaataaatctcgtcaaaaagagttaacccgctggttaaaacagcaaagcgtcatctcccaacgttggctgaatatttctcgtctgctgggctttgtgag<br>cggcatattgatcattgcccaggcctggttcatggcgcgtattctgcaacatatgattatggagaatattccccgtgaagccctgctgcttccctttacgttactggtt<br>ctgacctttgtactgcgcgcatgggtggtctggttacgcgaacgggtgggttatcacgccgggcagcatatccgctttgccatccgccgtcaggttctcgaccgtct<br>gcaacaagcagggccagcgtggattcagggtaaacctgcggggagctgggcgacgctggtactcgagcaaattgacgatatgcatgattactatgcacgctatc<br>tgccgcaaatggcgctggcagtgtcggtgccgttgctgattgtggtggcaatcttcccctctaactgggctgcggcgctcattctgctgggcactgcaccgttaattc<br>cgttgtttatggcgctggttggaatgggggctgccgatgctaaccgacgtaactttctcgctcttgctcgcttaagtgggcatttcctcgatcgcctgcgcggcatgg<br>aaacattgcgtatttttggtcgtggtgaagctgaaattgaaagtattcgttctgcttcggaagatttccgccaacggacaatggaagtgctacggctggcgtttttat<br>cctccggcattctcgaatttttacctcgctgtcaattgctctggtggcggtctactttggtttttcctatctcggcgagctggattttggtcactacgataccggtgtga<br>cgctggctgcgggttttctggccctgatccttgcgccagagtttttccagccattacgcgatctcggtacgttttatcatgctaaagcccaggctgttggcgcagctg<br>acagtctgaaaacgtttatggaaaccccgctcgcccatccgcaacgtggtgaggcggaattagcatcgaccgatccggtgaccattgaggccgaggagctgttta<br>tcacgtcgccggaaggtaaaacgctggccggaccgctgaactttactttgccagcaggccaacgtgcggtgttggttggtcgcagcggttcaggtaaaagctcac<br>tgctgaacgcgctttctggttttctctcatatcagggatcgctacgaatcaacgggatagaattacgcgatttatcaccagaatcatggcgtaaacatctctcctcgg<br>ttgggcaaaacccacaattaccggcagcaacattgcgggataacgtactactggcgcgacctgatgccagcgaacaagaattacaagcagcgctggataacgc<br>ctgggtcagcgagtttctaccgctcctcccacaaggcgttgatacgcctgttggcgaccaggctgcccgcctttccgtggggcaggcgcagcgcgtggcggtggcc<br>cgtgcgttactaaatccctgttcgctattactgttggatgaacccgctgccagccttgatgctcacagtgaacagcgcgtaatggaggcgctgaatgccgcctctct<br>gcgccagacaacgttaatggtcacccaccagttagaagatcttgctgactgggatgtcatttgggttatgcaggatggccggattattgagcaaggacgttacgcg<br>gaattaagtgtggctggtggcccattcgccacattactggcccatcgtcaggaggagatttaaatgcgcgctttgctaccctatctggcactgtataaacgtcataa<br>atggatgttaagtcttggtattgtgctggcaattgtgacgctgctcgccagtatcggtctgttgacactttccggctggttcctctcggcctcagcggttgcgggggtt<br>gccggactgtacagcttcaactatatgctacccgctgcgggcgtgcgtggcgcagcaatcacccgtactgccgggcgctattttgaacgtctggtaagtcacgacg<br>cgactttccgcgtgttgcagcatctgcgcatttacaccttcagcaaattgctgcccctctcccctgccggactggcgcgctatcgtcagggcgaattgctcaatcgcg<br>tggtggcggatgttgatacgctcgatcatctttacctgcgcgttatctcgccgctggtgggcgcttttgtggtgattatggtggtgacaatcgggttaagtttccttga<br>tttcaccctcgcctttacgctgggcggcattatgttactgacgcttttcctgatgccaccgctgttttatcgtgcgggaaaaagcaccgggcaaaatctgactcatctt<br>cgcggacagtatcgccaacaactgacggcctggctgcaagggcaagctgagctgaccatttttggtgccagcgatcgttatcgcacgcaactagagaatacaga<br>aattcaatggctggaagcgcaacgccgtcaatctgaactgaccgcattgtcgcaagcgataatgctgctcattggcgcgttagcggtgatcctgatgctgtggatg<br>gcgtctggcggcgttggcggcaatgctcaacccggcgcgttaattgccctgtttgtcttctgcgcgttagccgcgtttgaagcactggcaccagtaacgggtgcatt<br>tcagcatctggggcaagtcattgcctctgccgtacgtatctctgacttaacggatcaaaaaccggaggtcacctttcctgatacccaaactcgtgttgccgatcgcg<br>tttcgctgacgttacgggatgttcagttcacttatccggagcaatctcaacaggcacttaaagggatttctcttcaggtaaacgccggggaacatatagcgattctc<br>gggcgaaccggatgcggcaaatcaacactgttacaacagctgacccgcgcatgggacccgcaacagggcgagattttgcttaacgatagccccatagccagcct<br>gaatgaagcggctctacgacagaccatcagcgttgttcctcagcgagtgcatctgtttagcgccacgctgcgtgataatcttttactcgcctcgcctggcagtagtg<br>atgaggctctgtcggagatcttgcgtcgcgttggcctggaaaagctgctcgaggatgcaggtctcaacagttggttaggtgaaggcggacgccagctctccggtg<br>gtgaactgcgccgtctggctatcgcccgtgcgctgttacatgatgcgccactggtgttgctggatCAGcctaccgaaggcttagatgccacaaccgaaagccaga<br>tccttgaattgcttgcagaaatgatgcgtgagaaaacggtgttaatggtcacccatcgacttcgcggactctctcgtttccaacaaataatagtgatggacaacgg<br>gcaaattattgagcaaggtactcacgcagaactgcttgccagacaggggcgttattaccagttcaagcagggtttgtctgcaggcggtcgtggcagccaccatca<br>ccatcaccattaa |
| CydDC$^{H85A}$ | atgaattcgaataaatctcgtcaaaaagagttaacccgctggttaaaacagcaaagcgtcatctcccaacgttggctgaatatttctcgtctgctgggctttgtgag<br>cggcatattgatcattgcccaggcctggttcatggcgcgtattctgcaacatatgattatggagaatattccccgtgaagccctgctgcttccctttacgttactggtt<br>ctgacctttgtactgcgcgcatgggtggtctggttacgcgaacgggtgggttatcacgccgggcagcatatccgctttgccatccgccgtcaggttctcgaccgtct<br>gcaacaagcagggccagcgtggattcagggtaaacctgcggggagctgggcgacgctggtactcgagcaaattgacgatatgcatgattactatgcacgctatc<br>tgccgcaaatggcgctggcagtgtcggtgccgttgctgattgtggtggcaatcttcccctctaactgggctgcggcgctcattctgctgggcactgcaccgttaattc<br>cgttgtttatggcgctggttggaatgggggctgccgatgctaaccgacgtaactttctcgctcttgctcgcttaagtgggcatttcctcgatcgcctgcgcggcatgg<br>aaacattgcgtatttttggtcgtggtgaagctgaaattgaaagtattcgttctgcttcggaagatttccgccaacggacaatggaagtgctacggctggcgtttttat<br>cctccggcattctcgaatttttacctcgctgtcaattgctctggtggcggtctactttggtttttcctatctcggcgagctggattttggtcactacgataccggtgtga<br>cgctggctgcgggttttctggccctgatccttgcgccagagtttttccagccattacgcgatctcggtacgttttatcatgctaaagcccaggctgttggcgcagctg<br>acagtctgaaaacgtttatggaaaccccgctcgcccatccgcaacgtggtgaggcggaattagcatcgaccgatccggtgaccattgaggccgaggagctgttta<br>tcacgtcgccggaaggtaaaacgctggccggaccgctgaactttactttgccagcaggccaacgtgcggtgttggttggtcgcagcggttcaggtaaaagctcac<br>tgctgaacgcgctttctggttttctctcatatcagggatcgctacgaatcaacgggatagaattacgcgatttatcaccagaatcatggcgtaaacatctctcctcgg<br>ttgggcaaaacccacaattaccggcagcaacattgcgggataacgtactactggcgcgacctgatgccagcgaacaagaattacaagcagcgctggataacgc<br>ctgggtcagcgagtttctaccgctcctcccacaaggcgttgatacgcctgttggcgaccaggctgcccgcctttccgtggggcaggcgcagcgcgtggcggtggcc<br>cgtgcgttactaaatccctgttcgctattactgttggatgaacccgctgccagccttgatgctcacagtgaacagcgcgtaatggaggcgctgaatgccgcctctct<br>gcgccagacaacgttaatggtcacccaccagttagaagatcttgctgactgggatgtcatttgggttatgcaggatggccggattattgagcaaggacgttacgcg<br>gaattaagtgtggctggtggcccattcgccacattactggcccatcgtcaggaggagatttaaatgcgcgctttgctaccctatctggcactgtataaacgtcataa<br>atggatgttaagtcttggtattgtgctggcaattgtgacgctgctcgccagtatcggtctgttgacactttccggctggttcctctcggcctcagcggttgcgggggtt<br>gccggactgtacagcttcaactatatgctacccgctgcgggcgtgcgtggcgcagcaatcacccgtactgccgggcgctattttgaacgtctggtaagtgcagacg<br>cgactttccgcgtgttgcagcatctgcgcatttacaccttcagcaaattgctgcccctctcccctgccggactggcgcgctatcgtcagggcgaattgctcaatcgcg<br>tggtggcggatgttgatacgctcgatcatctttacctgcgcgttatctcgccgctggtgggcgcttttgtggtgattatggtggtgacaatcgggttaagtttccttga<br>tttcaccctcgcctttacgctgggcggcattatgttactgacgcttttcctgatgccaccgctgttttatcgtgcgggaaaaagcaccgggcaaaatctgactcatctt<br>cgcggacagtatcgccaacaactgacggcctggctgcaagggcaagctgagctgaccatttttggtgccagcgatcgttatcgcacgcaactagagaatacaga<br>aattcaatggctggaagcgcaacgccgtcaatctgaactgaccgcattgtcgcaagcgataatgctgctcattggcgcgttagcggtgatcctgatgctgtggatg<br>gcgtctggcggcgttggcggcaatgctcaacccggcgcgttaattgccctgtttgtcttctgcgcgttagccgcgtttgaagcactggcaccagtaacgggtgcatt<br>tcagcatctggggcaagtcattgcctctgccgtacgtatctctgacttaacggatcaaaaaccggaggtcacctttcctgatacccaaactcgtgttgccgatcgcg<br>tttcgctgacgttacgggatgttcagttcacttatccggagcaatctcaacaggcacttaaagggatttctcttcaggtaaacgccggggaacatatagcgattctc<br>gggcgaaccggatgcggcaaatcaacactgttacaacagctgacccgcgcatgggacccgcaacagggcgagattttgcttaacgatagccccatagccagcct<br>gaatgaagcggctctacgacagaccatcagcgttgttcctcagcgagtgcatctgtttagcgccacgctgcgtgataatcttttactcgcctcgcctggcagtagtg<br>atgaggctctgtcggagatcttgcgtcgcgttggcctggaaaagctgctcgaggatgcaggtctcaacagttggttaggtgaaggcggacgccagctctccggtg<br>gtgaactgcgccgtctggctatcgcccgtgcgctgttacatgatgcgccactggtgttgctggatgaacctaccgaaggcttagatgccacaaccgaaagccagat<br>ccttgaattgcttgcagaaatgatgcgtgagaaaacggtgttaatggtcacccatcgacttcgcggactctctcgtttccaacaaataatagtgatggacaacggg<br>caaattattgagcaaggtactcacgcagaactgcttgccagacaggggcgttattaccagttcaagcagggtttgtctgcaggcggtcgtggcagccaccatcac<br>catcaccattaa |

**Table S3 – Amino acid sequences of CydDC variants used in this study.**

| Variant | Amino acid sequence – CydD | Amino acid sequence – CydC |
|---|---|---|
| CydDC (wild-type) | MNSNKSRQKELTRWLKQQSVISQRWLNISRLLGFVSGILIIA<br>QAWFMARILQHMIMENIPREALLLPFTLLVLTFVLRAWVV<br>WLRERVGYHAGQHIRFAIRRQVLDRLQQAGPAWIQGKPA<br>GSWATLVLEQIDDMHDYYARYLPQMALAVSVPLLIVVAIFP<br>SNWAAALILLGTAPLIPLFMALVGMGAADANRRNFLALAR<br>LSGHFLDRLRGMETLRIFGRGEAEIESIRSASEDFRQRTMEV<br>LRLAFLSSGILEFFTSLSIALVAVYFGFSYLGELDFGHYDTGVT<br>LAAGFLALILAPEFFQPLRDLGTFYHAKAQAVGAADSLKTF<br>METPLAHPQRGEAELASTDPVTIEAEELFITSPEGKTLAGPL<br>NFTLPAGQRAVLVGRSGSGKSSLLNALSGFLSYQGSLRINGI<br>ELRDLSPESWRKHLSSVGQNPQLPAATLRDNVLLARPDASE<br>QELQAALDNAWVSEFLPLLPQGVDTPVGDQAARLSVGQA<br>QRVAVARALLNPCSLLLLDEPAASLDAHSEQRVMEALNAA<br>SLRQTTLMVTHQLEDLADWDVIWVMQDGRIIEQGRYAEL<br>SVAGGPFATLLAHRQEEI | MRALLPYLALYKRHKWMLSLGIVLAIVTLLASIGLLTLSGWF<br>LSASAVAGVAGLYSFNYMLPAAGVRGAAITRTAGRYFERLV<br>SHDATFRVLQHLRIYTFSKLLPLSPAGLARYRQGELLNRVVA<br>DVDTLDHLYLRVISPLVGAFVVIMVVTIGLSFLDFTLAFTLGG<br>IMLLTLFLMPPLFYRAGKSTGQNLTHLRGQYRQQLTAWLQ<br>GQAELTIFGASDRYRTQLENTEIQWLEAQRRQSELTALSQAI<br>MLLIGALAVILMLWMASGGVGGNAQPGALIALFVFCALAA<br>FEALAPVTGAFQHLGQVIASAVRISDLTDQKPEVTFPDTQT<br>RVADRVSLTLRDVQFTYPEQSQQALKGISLQVNAGEHIAIL<br>GRTGCGKSTLLQQLTRAWDPQQGEILLNDSPIASLNEAALR<br>QTISVVPQRVHLFSATLRDNLLLASPGSSDEALSEILRRVGLE<br>KLLEDAGLNSWLGEGGRQLSGGELRRLAIARALLHDAPLVL<br>LDEPTEGLDATTESQILELLAEMMREKTVLMVTHRLRGLSR<br>FQQIIVMDNGQIIEQGTHAELLARQGRYYQFKQGLSAGGR<br>GSHHHHHH |
| CydD$^{E511Q}$C | MNSNKSRQKELTRWLKQQSVISQRWLNISRLLGFVSGILIIA<br>QAWFMARILQHMIMENIPREALLLPFTLLVLTFVLRAWVV<br>WLRERVGYHAGQHIRFAIRRQVLDRLQQAGPAWIQGKPA<br>GSWATLVLEQIDDMHDYYARYLPQMALAVSVPLLIVVAIFP<br>SNWAAALILLGTAPLIPLFMALVGMGAADANRRNFLALAR<br>LSGHFLDRLRGMETLRIFGRGEAEIESIRSASEDFRQRTMEV<br>LRLAFLSSGILEFFTSLSIALVAVYFGFSYLGELDFGHYDTGVT<br>LAAGFLALILAPEFFQPLRDLGTFYHAKAQAVGAADSLKTF<br>METPLAHPQRGEAELASTDPVTIEAEELFITSPEGKTLAGPL<br>NFTLPAGQRAVLVGRSGSGKSSLLNALSGFLSYQGSLRINGI<br>ELRDLSPESWRKHLSSVGQNPQLPAATLRDNVLLARPDASE<br>QELQAALDNAWVSEFLPLLPQGVDTPVGDQAARLSVGQA<br>QRVAVARALLNPCSLLLLDQPAASLDAHSEQRVMEALNAA<br>SLRQTTLMVTHQLEDLADWDVIWVMQDGRIIEQGRYAEL<br>SVAGGPFATLLAHRQEEI | MRALLPYLALYKRHKWMLSLGIVLAIVTLLASIGLLTLSGWF<br>LSASAVAGVAGLYSFNYMLPAAGVRGAAITRTAGRYFERLV<br>SHDATFRVLQHLRIYTFSKLLPLSPAGLARYRQGELLNRVVA<br>DVDTLDHLYLRVISPLVGAFVVIMVVTIGLSFLDFTLAFTLGG<br>IMLLTLFLMPPLFYRAGKSTGQNLTHLRGQYRQQLTAWLQ<br>GQAELTIFGASDRYRTQLENTEIQWLEAQRRQSELTALSQAI<br>MLLIGALAVILMLWMASGGVGGNAQPGALIALFVFCALAA<br>FEALAPVTGAFQHLGQVIASAVRISDLTDQKPEVTFPDTQT<br>RVADRVSLTLRDVQFTYPEQSQQALKGISLQVNAGEHIAIL<br>GRTGCGKSTLLQQLTRAWDPQQGEILLNDSPIASLNEAALR<br>QTISVVPQRVHLFSATLRDNLLLASPGSSDEALSEILRRVGLE<br>KLLEDAGLNSWLGEGGRQLSGGELRRLAIARALLHDAPLVL<br>LDEPTEGLDATTESQILELLAEMMREKTVLMVTHRLRGLSR<br>FQQIIVMDNGQIIEQGTHAELLARQGRYYQFKQGLSAGGR<br>GSHHHHHH |
| CydDC$^{E500Q}$ | MNSNKSRQKELTRWLKQQSVISQRWLNISRLLGFVSGILIIA<br>QAWFMARILQHMIMENIPREALLLPFTLLVLTFVLRAWVV<br>WLRERVGYHAGQHIRFAIRRQVLDRLQQAGPAWIQGKPA<br>GSWATLVLEQIDDMHDYYARYLPQMALAVSVPLLIVVAIFP<br>SNWAAALILLGTAPLIPLFMALVGMGAADANRRNFLALAR<br>LSGHFLDRLRGMETLRIFGRGEAEIESIRSASEDFRQRTMEV<br>LRLAFLSSGILEFFTSLSIALVAVYFGFSYLGELDFGHYDTGVT<br>LAAGFLALILAPEFFQPLRDLGTFYHAKAQAVGAADSLKTF<br>METPLAHPQRGEAELASTDPVTIEAEELFITSPEGKTLAGPL<br>NFTLPAGQRAVLVGRSGSGKSSLLNALSGFLSYQGSLRINGI<br>ELRDLSPESWRKHLSSVGQNPQLPAATLRDNVLLARPDASE<br>QELQAALDNAWVSEFLPLLPQGVDTPVGDQAARLSVGQA<br>QRVAVARALLNPCSLLLLDEPAASLDAHSEQRVMEALNAA<br>SLRQTTLMVTHQLEDLADWDVIWVMQDGRIIEQGRYAEL<br>SVAGGPFATLLAHRQEEI | MRALLPYLALYKRHKWMLSLGIVLAIVTLLASIGLLTLSGWF<br>LSASAVAGVAGLYSFNYMLPAAGVRGAAITRTAGRYFERLV<br>SHDATFRVLQHLRIYTFSKLLPLSPAGLARYRQGELLNRVVA<br>DVDTLDHLYLRVISPLVGAFVVIMVVTIGLSFLDFTLAFTLGG<br>IMLLTLFLMPPLFYRAGKSTGQNLTHLRGQYRQQLTAWLQ<br>GQAELTIFGASDRYRTQLENTEIQWLEAQRRQSELTALSQAI<br>MLLIGALAVILMLWMASGGVGGNAQPGALIALFVFCALAA<br>FEALAPVTGAFQHLGQVIASAVRISDLTDQKPEVTFPDTQT<br>RVADRVSLTLRDVQFTYPEQSQQALKGISLQVNAGEHIAIL<br>GRTGCGKSTLLQQLTRAWDPQQGEILLNDSPIASLNEAALR<br>QTISVVPQRVHLFSATLRDNLLLASPGSSDEALSEILRRVGLE<br>KLLEDAGLNSWLGEGGRQLSGGELRRLAIARALLHDAPLVL<br>LDQPTEGLDATTESQILELLAEMMREKTVLMVTHRLRGLSR<br>FQQIIVMDNGQIIEQGTHAELLARQGRYYQFKQGLSAGGR<br>GSHHHHHH |
| CydDC$^{H85A}$ | MNSNKSRQKELTRWLKQQSVISQRWLNISRLLGFVSGILIIA<br>QAWFMARILQHMIMENIPREALLLPFTLLVLTFVLRAWVV<br>WLRERVGYHAGQHIRFAIRRQVLDRLQQAGPAWIQGKPA<br>GSWATLVLEQIDDMHDYYARYLPQMALAVSVPLLIVVAIFP<br>SNWAAALILLGTAPLIPLFMALVGMGAADANRRNFLALAR<br>LSGHFLDRLRGMETLRIFGRGEAEIESIRSASEDFRQRTMEV<br>LRLAFLSSGILEFFTSLSIALVAVYFGFSYLGELDFGHYDTGVT<br>LAAGFLALILAPEFFQPLRDLGTFYHAKAQAVGAADSLKTF<br>METPLAHPQRGEAELASTDPVTIEAEELFITSPEGKTLAGPL<br>NFTLPAGQRAVLVGRSGSGKSSLLNALSGFLSYQGSLRINGI<br>ELRDLSPESWRKHLSSVGQNPQLPAATLRDNVLLARPDASE<br>QELQAALDNAWVSEFLPLLPQGVDTPVGDQAARLSVGQA<br>QRVAVARALLNPCSLLLLDEPAASLDAHSEQRVMEALNAA<br>SLRQTTLMVTHQLEDLADWDVIWVMQDGRIIEQGRYAEL<br>SVAGGPFATLLAHRQEEI | MRALLPYLALYKRHKWMLSLGIVLAIVTLLASIGLLTLSGWF<br>LSASAVAGVAGLYSFNYMLPAAGVRGAAITRTAGRYFERLV<br>SADATFRVLQHLRIYTFSKLLPLSPAGLARYRQGELLNRVVA<br>DVDTLDHLYLRVISPLVGAFVVIMVVTIGLSFLDFTLAFTLGG<br>IMLLTLFLMPPLFYRAGKSTGQNLTHLRGQYRQQLTAWLQ<br>GQAELTIFGASDRYRTQLENTEIQWLEAQRRQSELTALSQAI<br>MLLIGALAVILMLWMASGGVGGNAQPGALIALFVFCALAA<br>FEALAPVTGAFQHLGQVIASAVRISDLTDQKPEVTFPDTQT<br>RVADRVSLTLRDVQFTYPEQSQQALKGISLQVNAGEHIAIL<br>GRTGCGKSTLLQQLTRAWDPQQGEILLNDSPIASLNEAALR<br>QTISVVPQRVHLFSATLRDNLLLASPGSSDEALSEILRRVGLE<br>KLLEDAGLNSWLGEGGRQLSGGELRRLAIARALLHDAPLVL<br>LDEPTEGLDATTESQILELLAEMMREKTVLMVTHRLRGLSR<br>FQQIIVMDNGQIIEQGTHAELLARQGRYYQFKQGLSAGGR<br>GSHHHHHH |

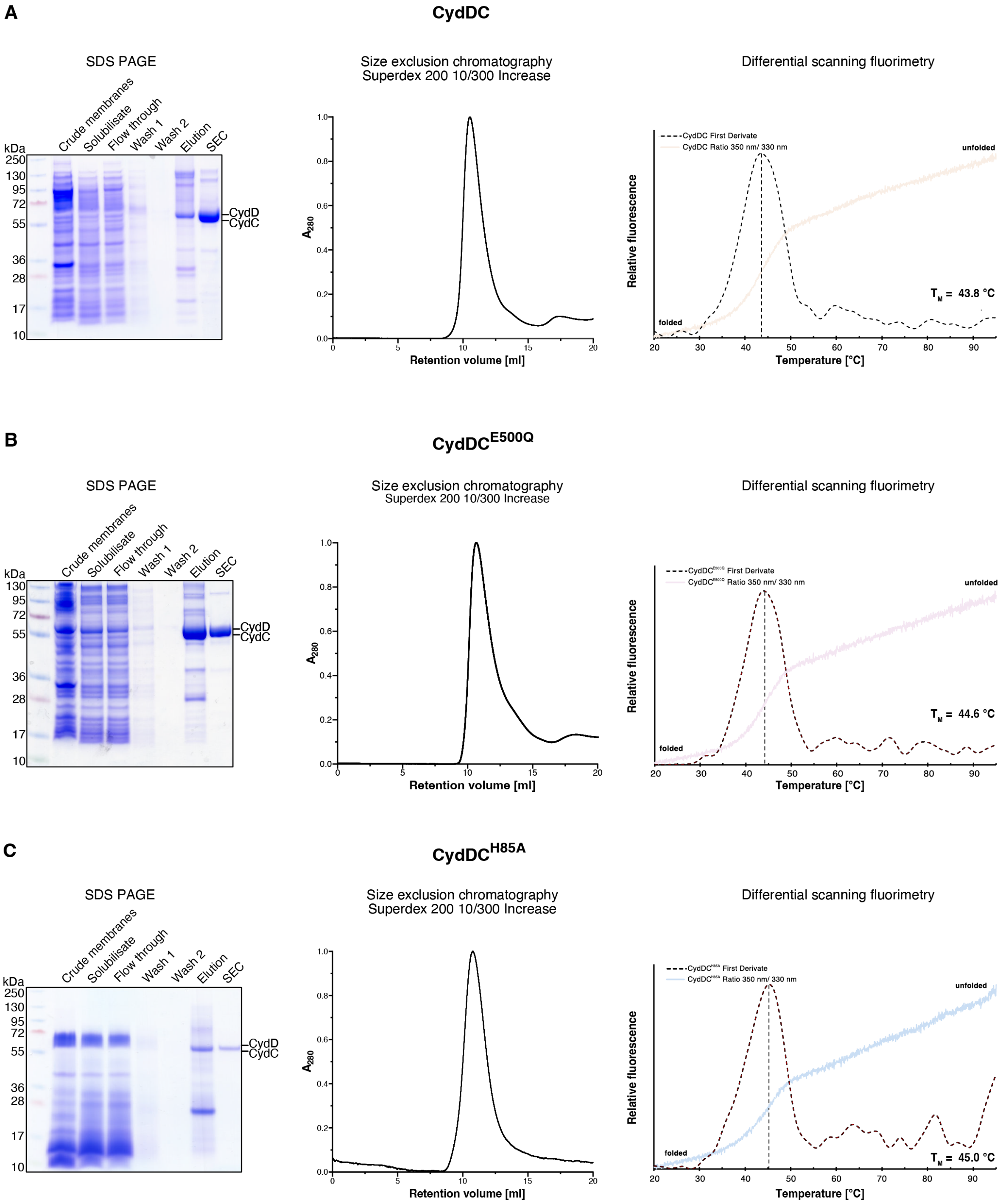

**Fig. S1 – Purification and characterization of CydDC variants. (A)** Wild-type, **(B)** E500Q$^{C}$, and **(C)** H85A$^{C}$ variants of CydDC were produced in *Escherichia coli* BL21 (DE3) cells and purified by immobilized metal ion affinity chromatography (Co-IMAC). Peak fractions were collected and analyzed by SDS-PAGE, size exclusion chromatography (SEC), nanoscale differential scanning fluorimetry (nanoDSF) and used for downstream activity assays and cryo-EM specimen preparation.

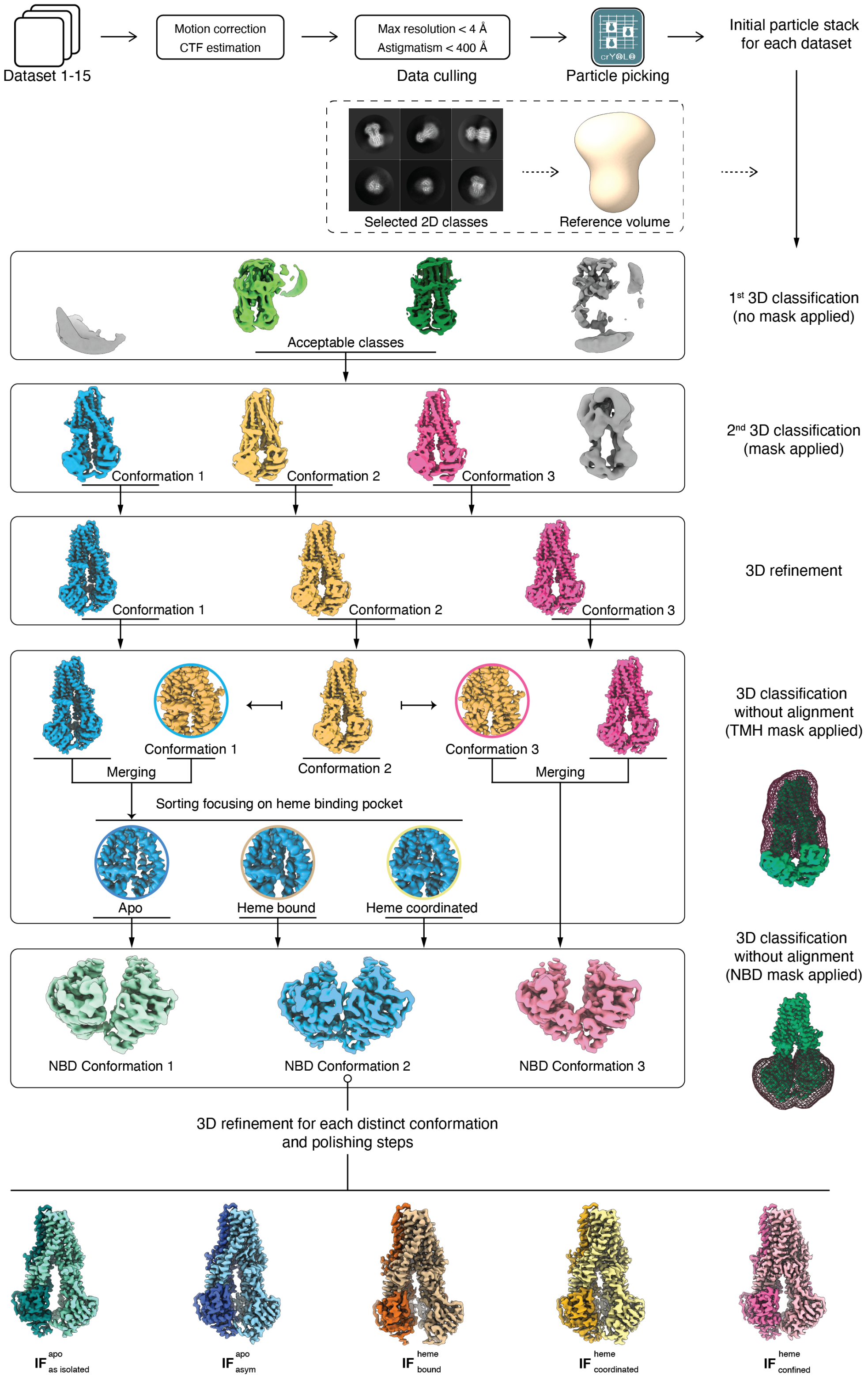

**Fig. S2 – Exemplary cryo-EM data processing workflow of wild-type CydDC datasets.** Datasets 1 - 16 were processed according to the depicted workflow scheme. Statistical values of datasets 1 - 16 are summarized in table S1. Initial full-frame motion correction was performed with MotionCorr2 (RELION-3.1). CTF estimation was performed using Gctf (version 1.06). Particles were picked using crYOLO and subsequently extracted in the RELION-3.1 suite. Particles contributing to 2D classes with distinct features were selected for further processing. A subset of these particles was used for initial model generation. Three-dimensional classification, CTF refinement, Bayesian polishing, and consensus 3D refinement were performed in RELION-3.1.

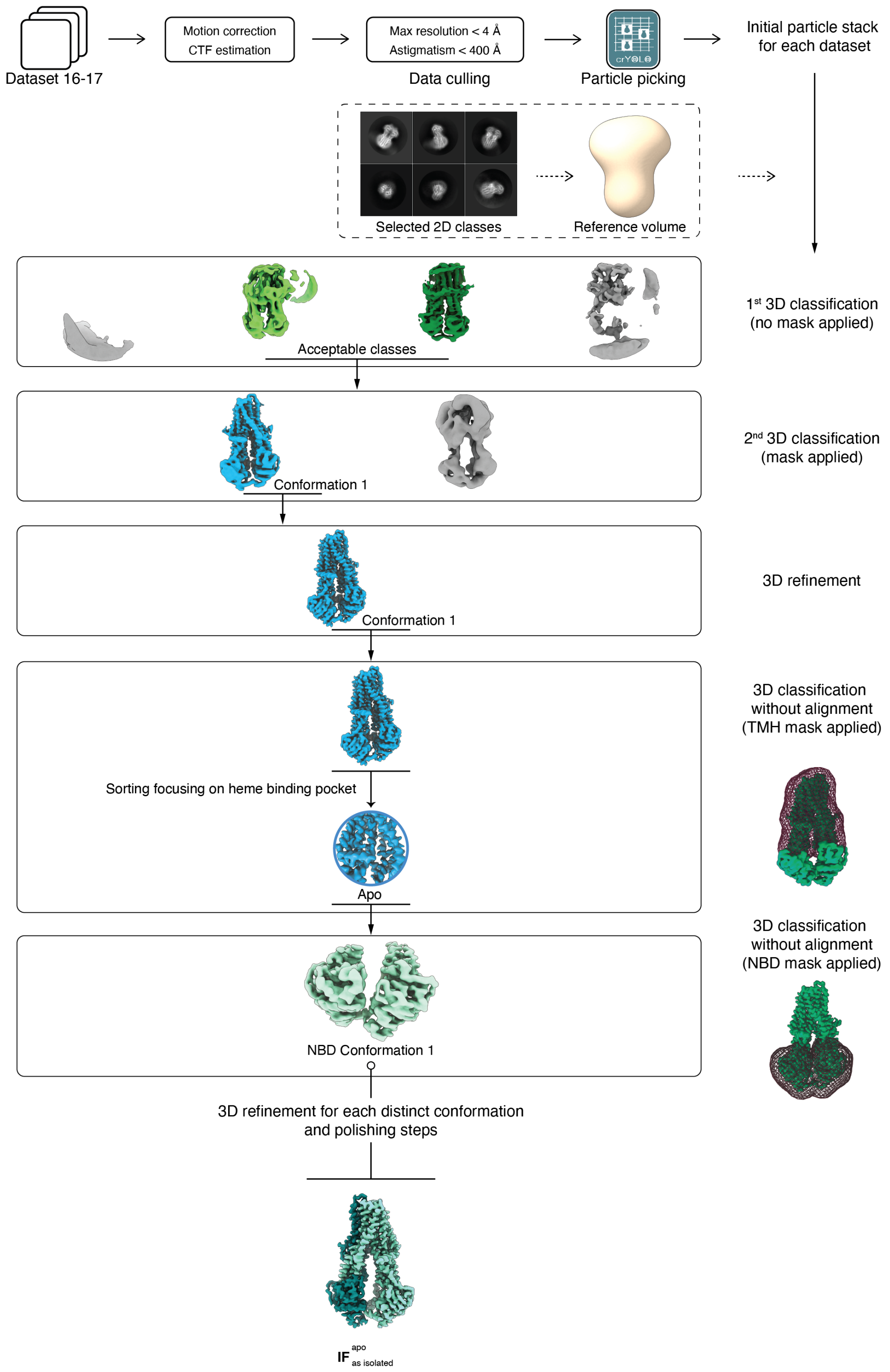

**Fig. S3 – Exemplary cryo-EM data processing workflow of H85A$^{C}$ CydDC datasets.** Datasets 16 - 17 were processed according to the depicted workflow scheme. Statistical values of datasets 16 - 17 are summarized in table S1. Initial full-frame motion correction was performed with MotionCorr2 (RELION-3.1). CTF estimation was performed using Gctf (version 1.06). Particles were picked using crYOLO and subsequently extracted in the RELION-3.1 suite. Particles contributing to 2D classes with distinct features were selected for further processing. A subset of these particles was used for initial model generation. Three-dimensional classification, CTF refinement, Bayesian polishing, and consensus 3D refinement were performed in RELION-3.1.

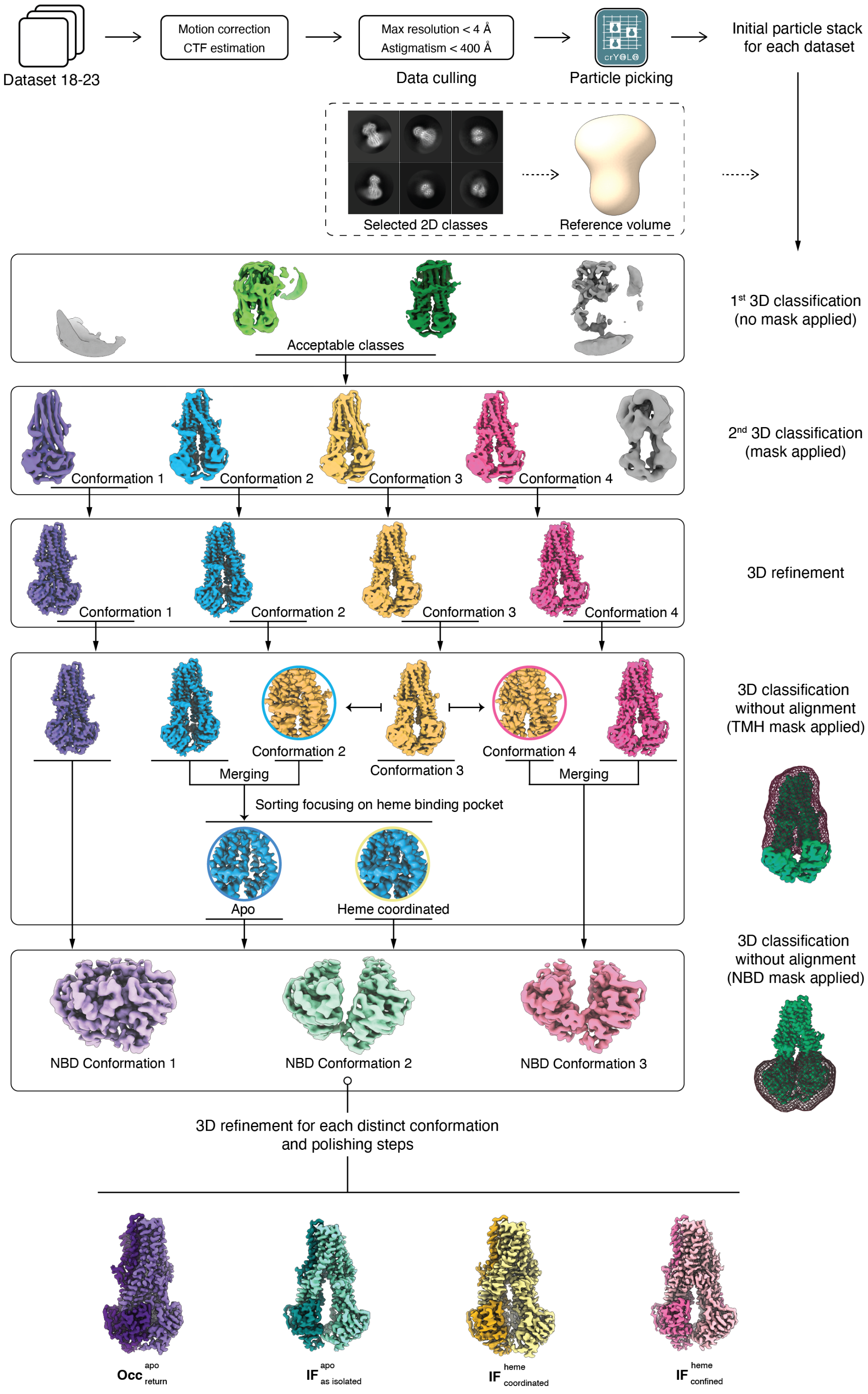

**Fig. S4 – Exemplary cryo-EM data processing workflow of E500Q$^{C}$ CydDC datasets.** Datasets 18 - 23 were processed according to the depicted workflow scheme. Statistical values of datasets 18 - 23 are summarized in table S1. Initial full-frame motion correction was performed with MotionCorr2 (RELION-3.1). CTF estimation was performed using Gctf (version 1.06). Particles were picked using crYOLO and subsequently extracted in the RELION-3.1 suite. Particles contributing to 2D classes with distinct features were selected for further processing. A subset of these particles was used for initial model generation. Three-dimensional classification, CTF refinement, Bayesian polishing, and consensus 3D refinement were performed in RELION-3.1.

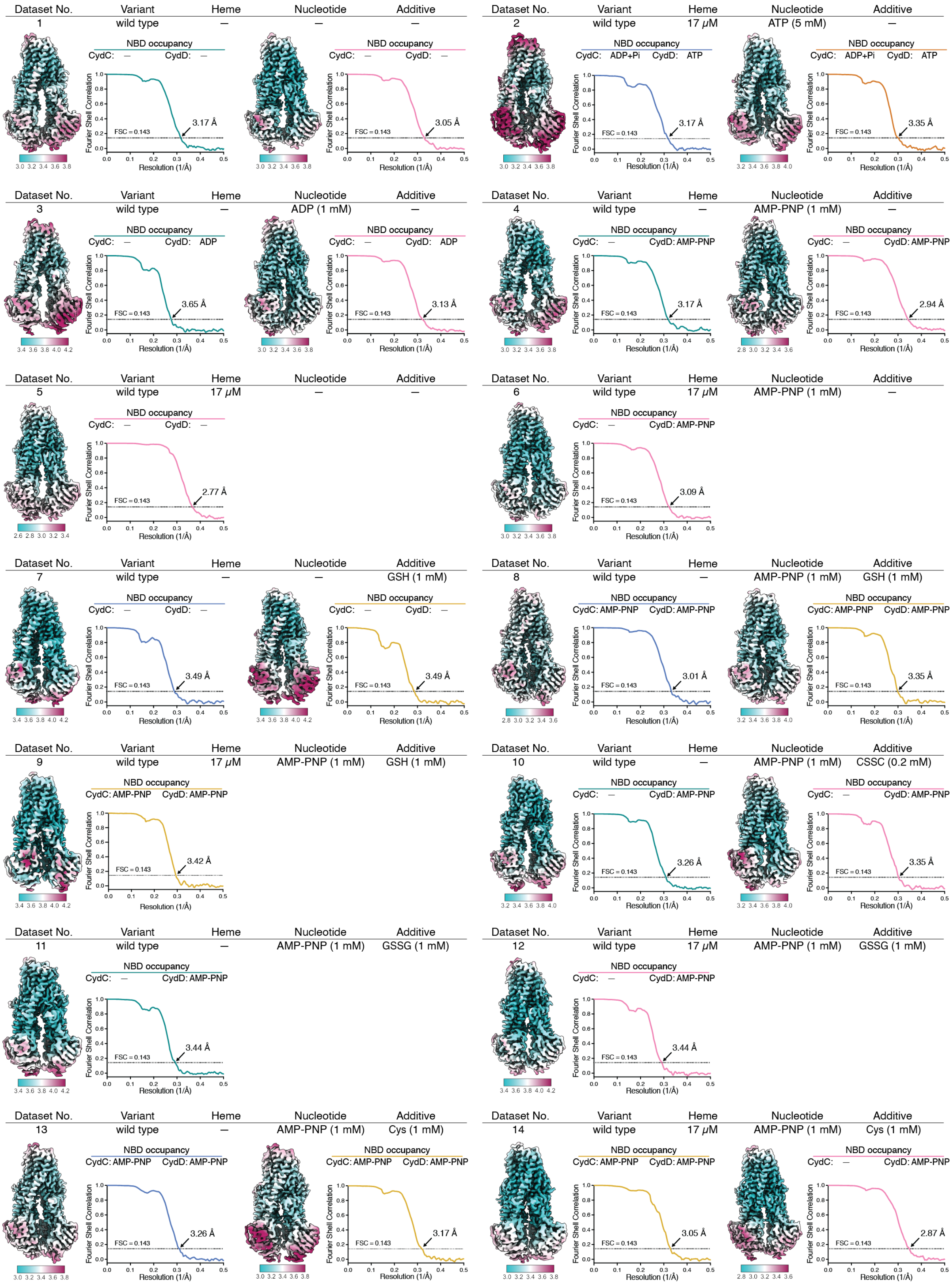

Dataset No. | Variant | Heme | Nucleotide | Additive
1 | wild type | – | – | –
NBD occupancy
CydC: – CydD: –
Fourier Shell Correlation
Resolution (1/Å)
FSC = 0.143
3.17 Å
3.05 Å
2 | wild type | 17 μM | ATP (5 mM) | –
CydC: ADP+Pi CydD: ATP
3.17 Å
3.35 Å
3 | wild type | – | ADP (1 mM) | –
CydC: – CydD: ADP
3.65 Å
3.13 Å
4 | wild type | – | AMP-PNP (1 mM) | –
CydC: – CydD: AMP-PNP
3.17 Å
2.94 Å
5 | wild type | 17 μM | – | –
CydC: – CydD: –
2.77 Å
6 | wild type | 17 μM | AMP-PNP (1 mM) | –
CydC: – CydD: AMP-PNP
3.09 Å
7 | wild type | – | – | GSH (1 mM)
CydC: – CydD: –
3.49 Å
3.49 Å
8 | wild type | – | AMP-PNP (1 mM) | GSH (1 mM)
CydC: AMP-PNP CydD: AMP-PNP
3.01 Å
3.35 Å
9 | wild type | 17 μM | AMP-PNP (1 mM) | GSH (1 mM)
CydC: AMP-PNP CydD: AMP-PNP
3.42 Å
10 | wild type | – | AMP-PNP (1 mM) | CSSC (0.2 mM)
CydC: – CydD: AMP-PNP
3.26 Å
3.35 Å
11 | wild type | – | AMP-PNP (1 mM) | GSSG (1 mM)
CydC: – CydD: AMP-PNP
3.44 Å
12 | wild type | 17 μM | AMP-PNP (1 mM) | GSSG (1 mM)
CydC: – CydD: AMP-PNP
3.44 Å
13 | wild type | – | AMP-PNP (1 mM) | Cys (1 mM)
CydC: AMP-PNP CydD: AMP-PNP
3.26 Å
3.17 Å
14 | wild type | 17 μM | AMP-PNP (1 mM) | Cys (1 mM)
CydC: AMP-PNP CydD: AMP-PNP
3.05 Å
CydC: – CydD: AMP-PNP
2.87 Å

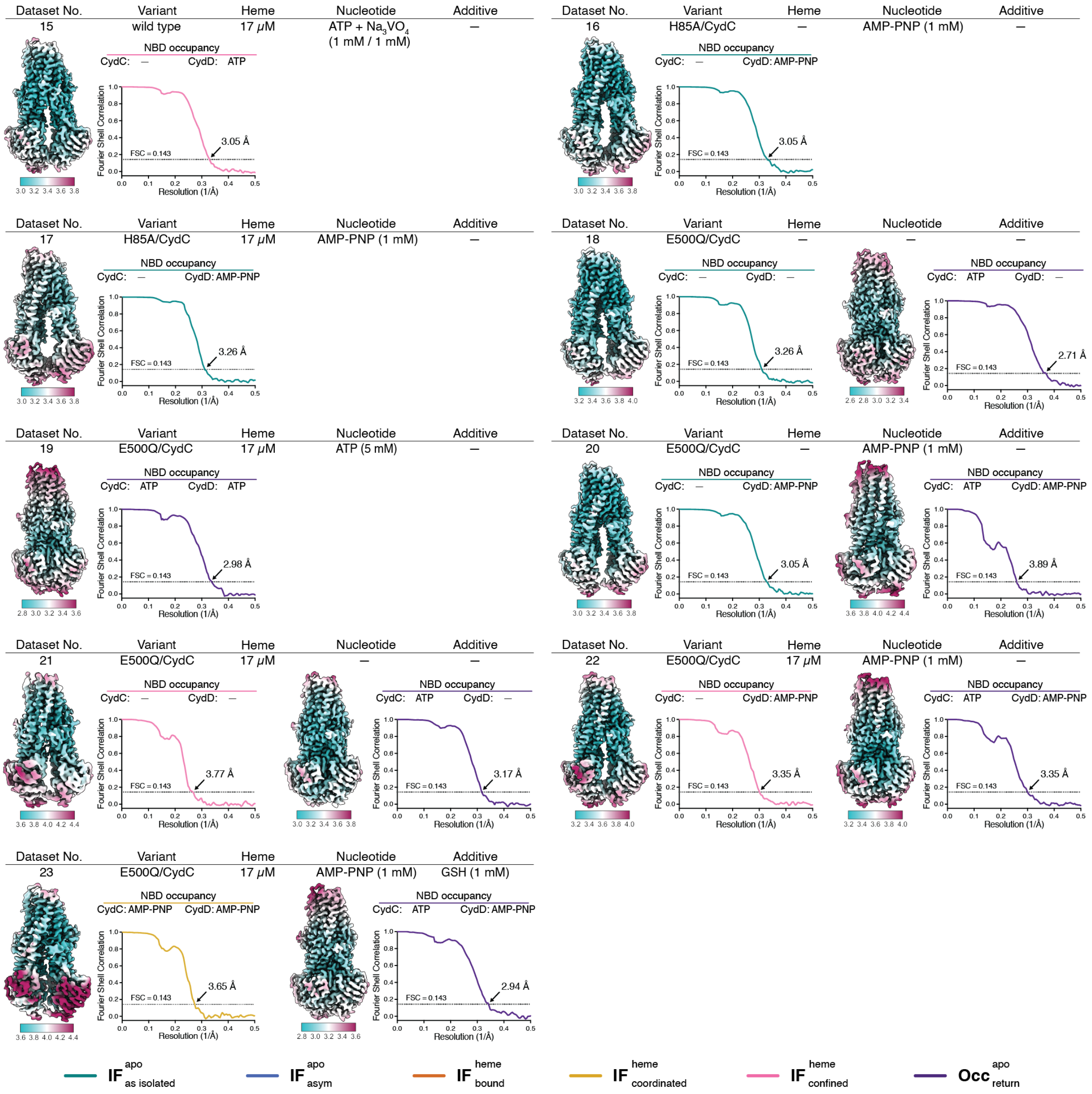

**Fig. S5 – Local resolution distributions, NBD occupancy, and Fourier shell correlations of CydDC structures.** Color-coded density maps indicate local differences in resolution. The NBD occupancy of each individual map is indicated. Fourier shell correlation plots show average resolutions at $FSC_{0.143}$. Colors of FSC curves correspond to CydDC conformations according to the key at the bottom of the figure. Sample components and their respective concentrations are given for each dataset.

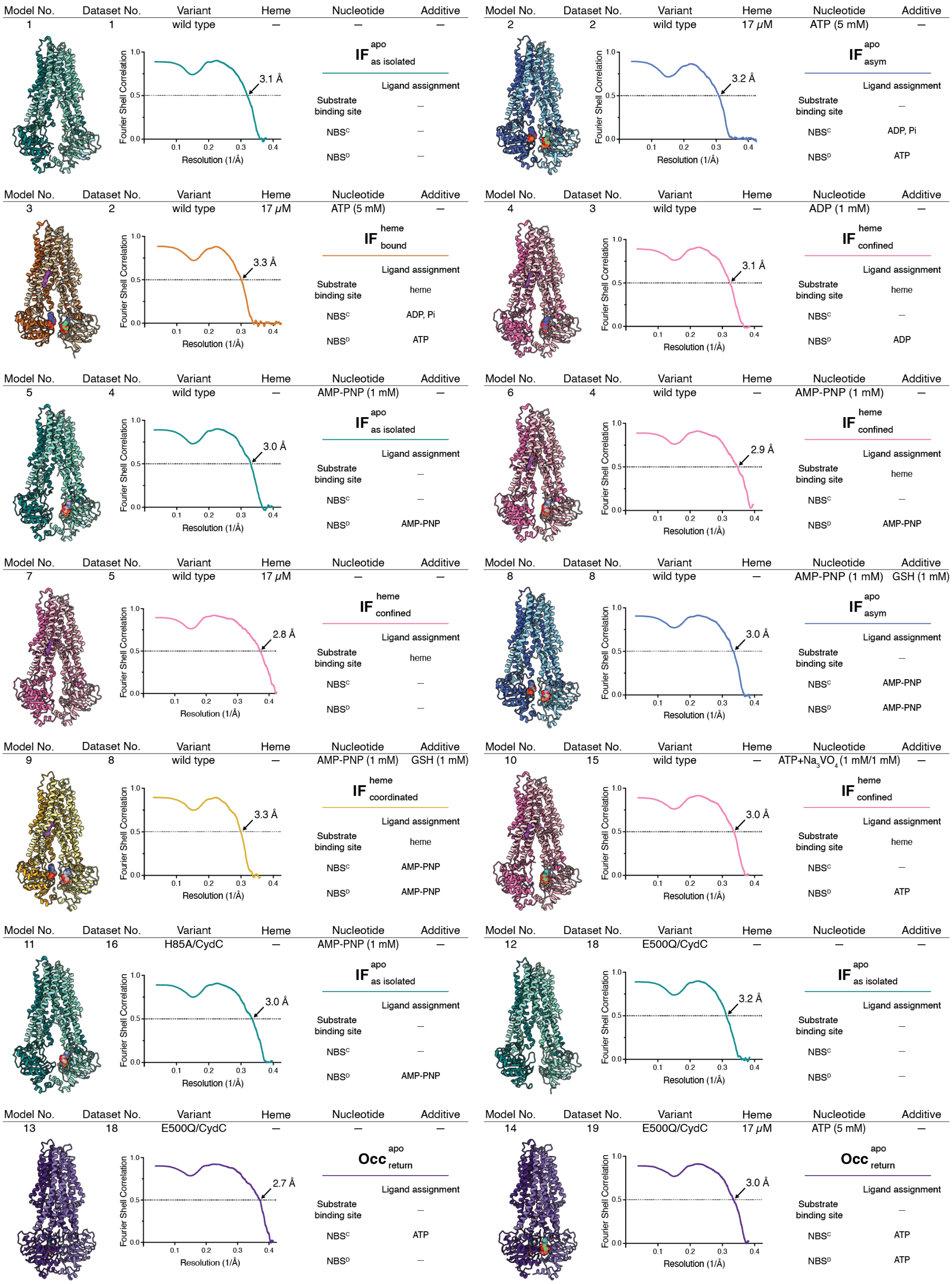
Model No. Dataset No. Variant Heme Nucleotide Additive
1 1 wild type — — —
IF apo as isolated
3.1 Å
Ligand assignment
Substrate binding site —
NBSC —
NBSD —
Model No. Dataset No. Variant Heme Nucleotide Additive
2 2 wild type 17 μM ATP (5 mM) —
IF apo asym
3.2 Å
Ligand assignment
Substrate binding site —
NBSC ADP, Pi
NBSD ATP
Model No. Dataset No. Variant Heme Nucleotide Additive
3 2 wild type 17 μM ATP (5 mM) —
IF heme bound
3.3 Å
Ligand assignment
Substrate binding site heme
NBSC ADP, Pi
NBSD ATP
Model No. Dataset No. Variant Heme Nucleotide Additive
4 3 wild type — ADP (1 mM) —
IF heme confined
3.1 Å
Ligand assignment
Substrate binding site heme
NBSC —
NBSD ADP
Model No. Dataset No. Variant Heme Nucleotide Additive
5 4 wild type — AMP-PNP (1 mM) —
IF apo as isolated
3.0 Å
Ligand assignment
Substrate binding site —
NBSC —
NBSD AMP-PNP
Model No. Dataset No. Variant Heme Nucleotide Additive
6 4 wild type — AMP-PNP (1 mM) —
IF heme confined
2.9 Å
Ligand assignment
Substrate binding site heme
NBSC —
NBSD AMP-PNP
Model No. Dataset No. Variant Heme Nucleotide Additive
7 5 wild type 17 μM — —
IF heme confined
2.8 Å
Ligand assignment
Substrate binding site heme
NBSC —
NBSD —
Model No. Dataset No. Variant Heme Nucleotide Additive
8 8 wild type — AMP-PNP (1 mM) GSH (1 mM)
IF apo asym
3.0 Å
Ligand assignment
Substrate binding site —
NBSC AMP-PNP
NBSD AMP-PNP
Model No. Dataset No. Variant Heme Nucleotide Additive
9 8 wild type — AMP-PNP (1 mM) GSH (1 mM)
IF heme coordinated
3.3 Å
Ligand assignment
Substrate binding site heme
NBSC AMP-PNP
NBSD AMP-PNP
Model No. Dataset No. Variant Heme Nucleotide Additive
10 15 wild type — ATP+Na3VO4 (1 mM/1 mM) —
IF heme confined
3.0 Å
Ligand assignment
Substrate binding site heme
NBSC —
NBSD ATP
Model No. Dataset No. Variant Heme Nucleotide Additive
11 16 H85A/CydC — AMP-PNP (1 mM) —
IF apo as isolated
3.0 Å
Ligand assignment
Substrate binding site —
NBSC —
NBSD AMP-PNP
Model No. Dataset No. Variant Heme Nucleotide Additive
12 18 E500Q/CydC — — —
IF apo as isolated
3.2 Å
Ligand assignment
Substrate binding site —
NBSC —
NBSD —
Model No. Dataset No. Variant Heme Nucleotide Additive
13 18 E500Q/CydC — — —
Occ apo return
2.7 Å
Ligand assignment
Substrate binding site —
NBSC ATP
NBSD —
Model No. Dataset No. Variant Heme Nucleotide Additive
14 19 E500Q/CydC 17 μM ATP (5 mM) —
Occ apo return
3.0 Å
Ligand assignment
Substrate binding site —
NBSC ATP
NBSD ATP
Fourier Shell Correlation
Resolution (1/Å)

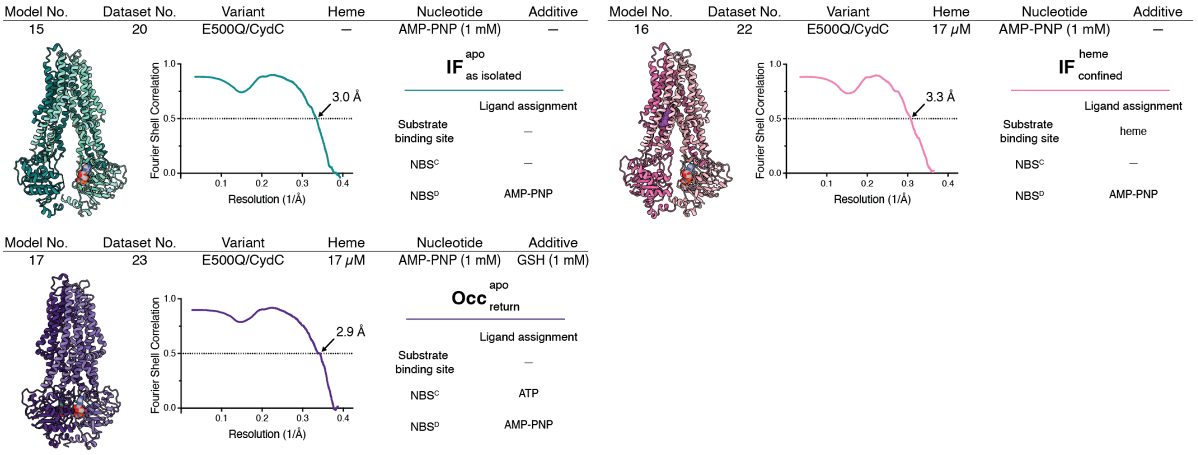

**Fig. S6 – Map-to-model correlations.** Ribbon models corresponding to conformations identified in respective cryo-EM datasets are shown. Fourier shell plots show map-to-model correlations. Resolution estimates are based on $FSC_{0.5}$ level. Ligand assignments are listed for each model and dataset. Colors of models and FSC curves correspond to CydDC conformations: $\mathbf{IF}^{\mathrm{apo}}_{\mathrm{asym}}$, blue; $\mathbf{IF}^{\mathrm{apo}}_{\mathrm{as\ isolated}}$, green; $\mathbf{IF}^{\mathrm{heme}}_{\mathrm{bound}}$, orange; $\mathbf{IF}^{\mathrm{heme}}_{\mathrm{coordinated}}$, yellow; $\mathbf{IF}^{\mathrm{heme}}_{\mathrm{confined}}$, magenta; $\mathbf{Occ}^{\mathrm{apo}}_{\mathrm{return}}$, purple. Ligand colors: ATP, green; ADP, purple; AMP-PNP, grey; heme, violet.

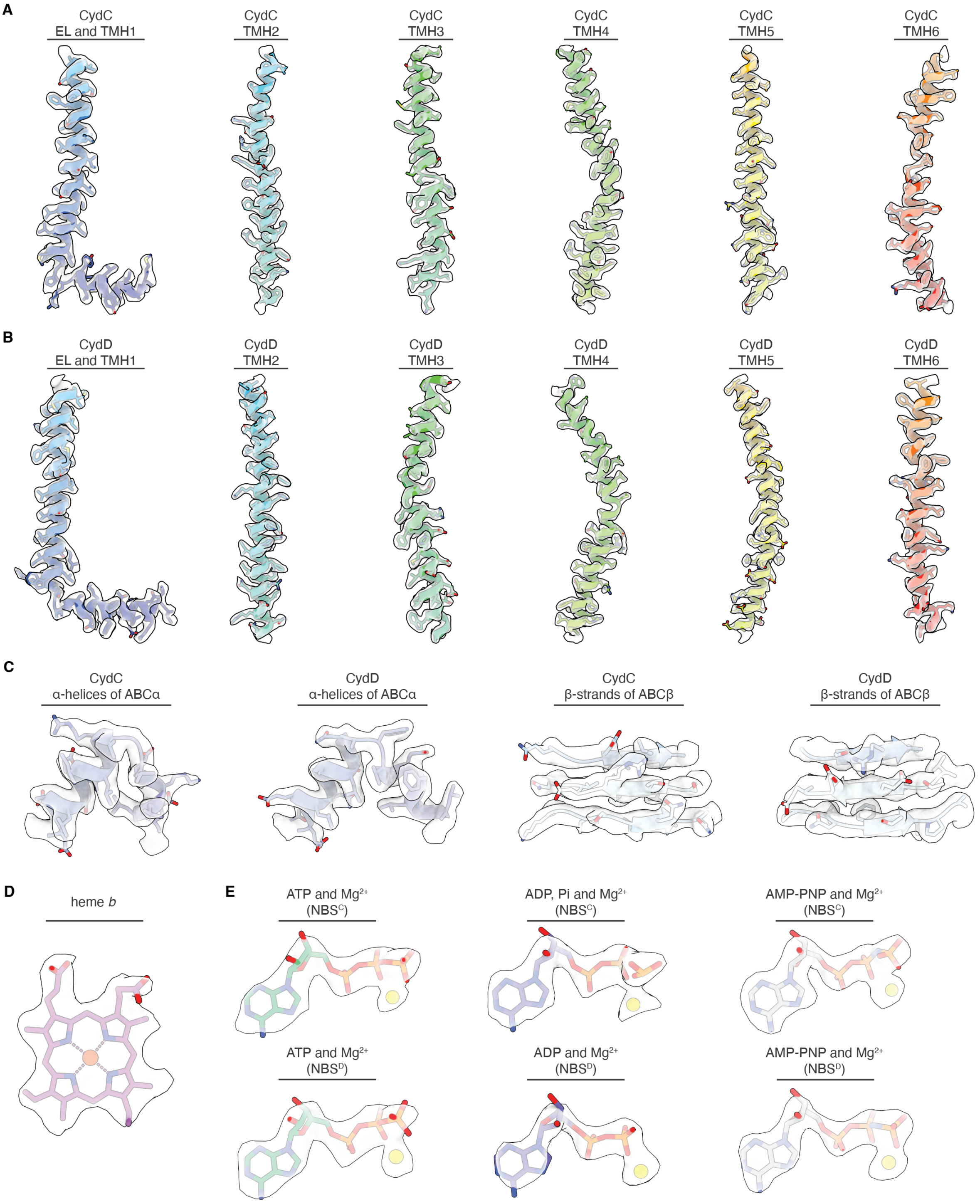

**Fig. S7 – Representative density features of CydDC.** Visual inspection of density features of **(A - B)** transmembrane helices, **(C)** nucleotide-binding domain motifs, and **(D-E)** substrates/inhibitors (heme, ATP, ADP + Pi, and AMP-PNP). Symmetry related structural elements are presented in matching color codes. Presented densities were sharpened by b-factors of between -80 and -48.

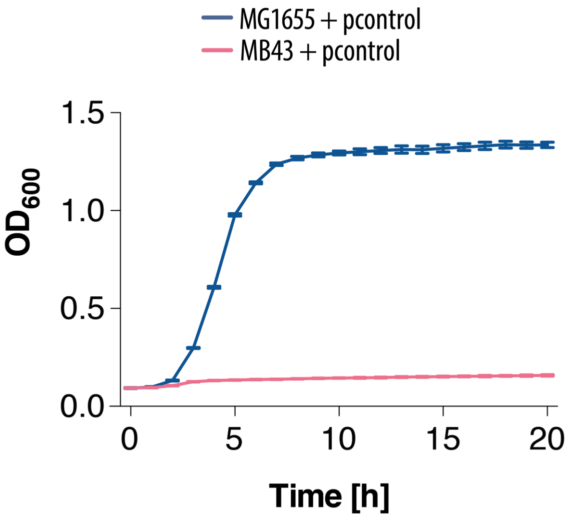

**Fig. S8 – Comparative growth of Escherichia coli strains MG1655 and MB43.** MG1655 is the wild-type strain and MB43 is the mutant strain with impaired respiration. Both strains were transformed with the empty pET control vector (pcontrol).

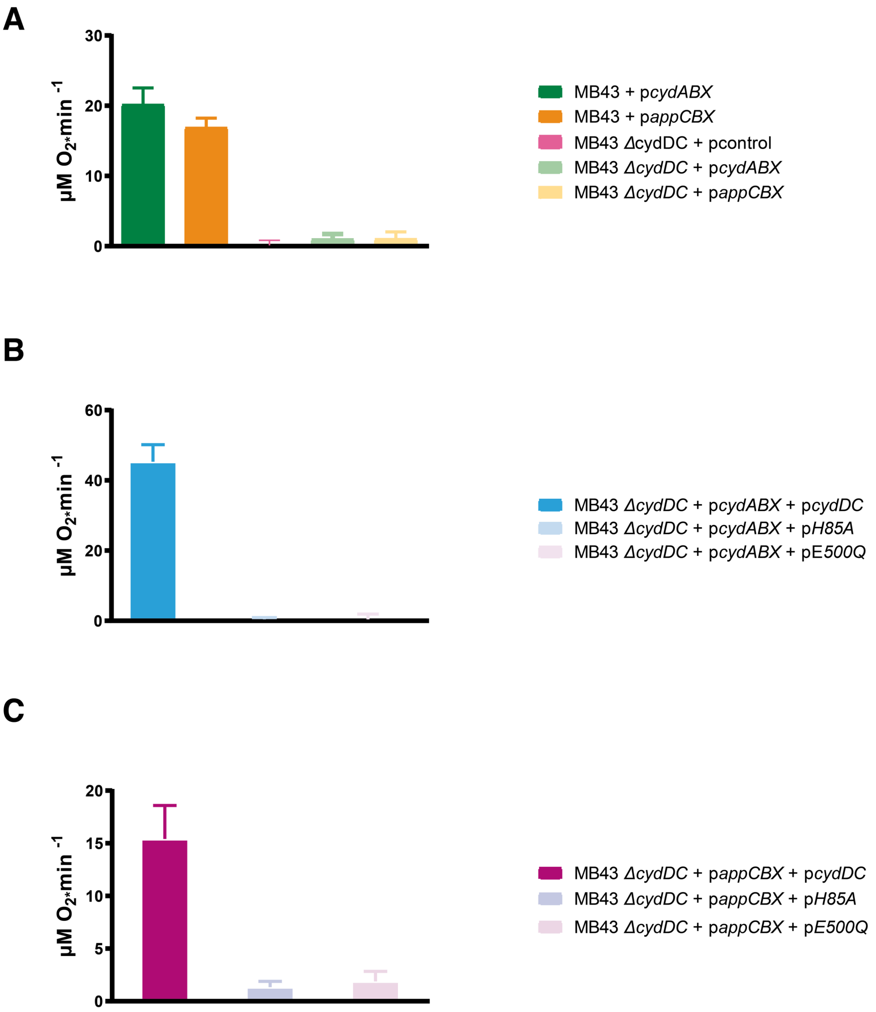

**Fig. S9 – Oxygen reductase activity of membranes isolated from bacterial strains.** (**A**) Activity of MB43 and MB43Δ*cydDC* strains complemented with structural genes for cytochrome *bd*-I (*cydABX*) or cytochrome *bd*-II (*appCBX*), (**B**) activity of MB43*cydDC* complemented with structural genes for cytochrome *bd*-I (*cydABX*) and for wild-type resp. mutant CydDC, (**C**) activity of MB43*cydDC* complemented with structural genes for cytochrome *bd*-II (*appCBX*) and for wild-type resp. mutant CydDC. Oxygen concentrations were determined with a Clark-type electrode using membranes (5 µg for panels A and B, 15 µg for panel C). Samples were preincubated in buffer and subsequently the reaction was started by addition of the DTT/ubiquinol-1 substrate mixture. All data are presented as mean of n=3 ± SD.

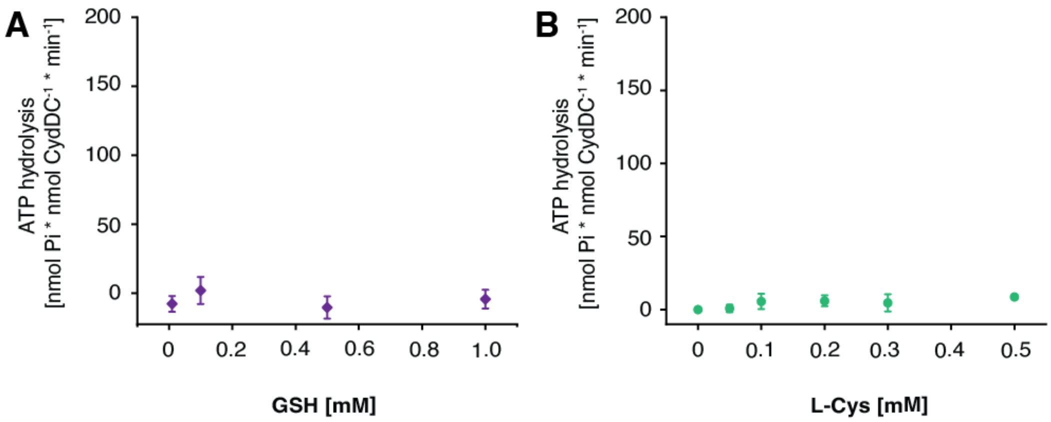

**Fig. S10 – GSH and L-Cys dependent ATPase activity of CydDC.** The concentration dependent **(A)** GSH (0.01 - 1 mM) and **(B)** L-Cys (0.05 - 0.5 mM) ATPase activity of CydDC was determined by a Malachite green phosphate assay. All presented ATP hydrolysis data are corrected for background activity in the absence of substrate candidates. All data are presented as mean of n=3 ± SD.

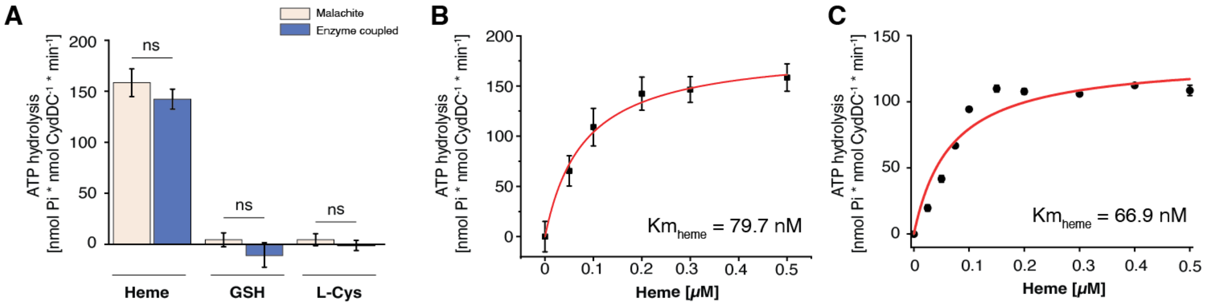

**Fig. S11 – Assessment of reductant induced ATPase activity of CydDC. (A)** ATP hydrolysis activity of CydDC was determined in the presence of heme (0.5 µM), GSH (1 mM), and L-Cys (0.5 mM) using the Malachite green phosphate and the PK/LDH coupled assays, respectively. Significance was assessed based on a paired two-tailed Student's t-test. **(B-C)** Michaelis constants ($K_M^{app}$) of CydDC for heme determined by Malachite green phosphate (79.7 nM) and PK/LDH coupled assays (66.9 nM), respectively. All presented ATP hydrolysis data are corrected for background activity in the absence of substrate candidates. All data are presented as mean of n=3 ± SD. Ns, not significant.

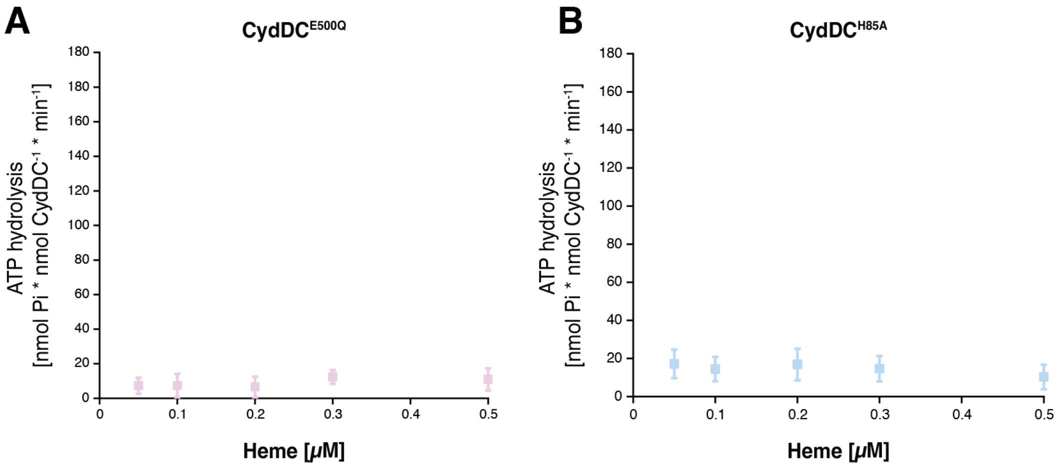

**Fig. S12 – ATPase activity of CydDC$^{E500Q}$ and CydDC$^{H85A}$**

(**A**) Heme (0.05 - 0.5 mM) dependent ATP hydrolysis activity of CydDC$^{E500Q}$ determined by Malachite green phosphate assay. **(B)** Heme (0.05 - 0.5 mM) dependent ATP hydrolysis activity of CydDC$^{H85A}$ determined by Malachite green phosphatase assay. All presented ATP hydrolysis data are corrected for background activity in the absence of heme. All data are presented as mean of n=3 ± SD.

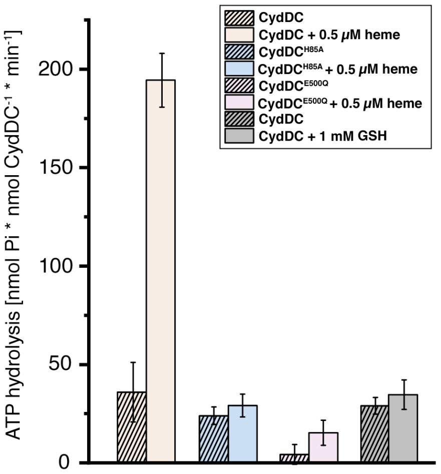

**Fig. S13 – Mutant variants influence the background ATPase activity of CydDC.** Total ATPase activities of wild-type, H85A$^{C}$, and E500Q$^{C}$ variants of CydDC show that unspecific background hydrolysis activity is slightly reduced by the histidine to alanine exchange of the characterized heme binding site, and strongly by the Walker B glutamate to glutamine exchange in NBS$^{C}$. Total ATPase activity of wild-type CydDC in the presence and absence of GSH exemplifies a non-stimulatory effect.

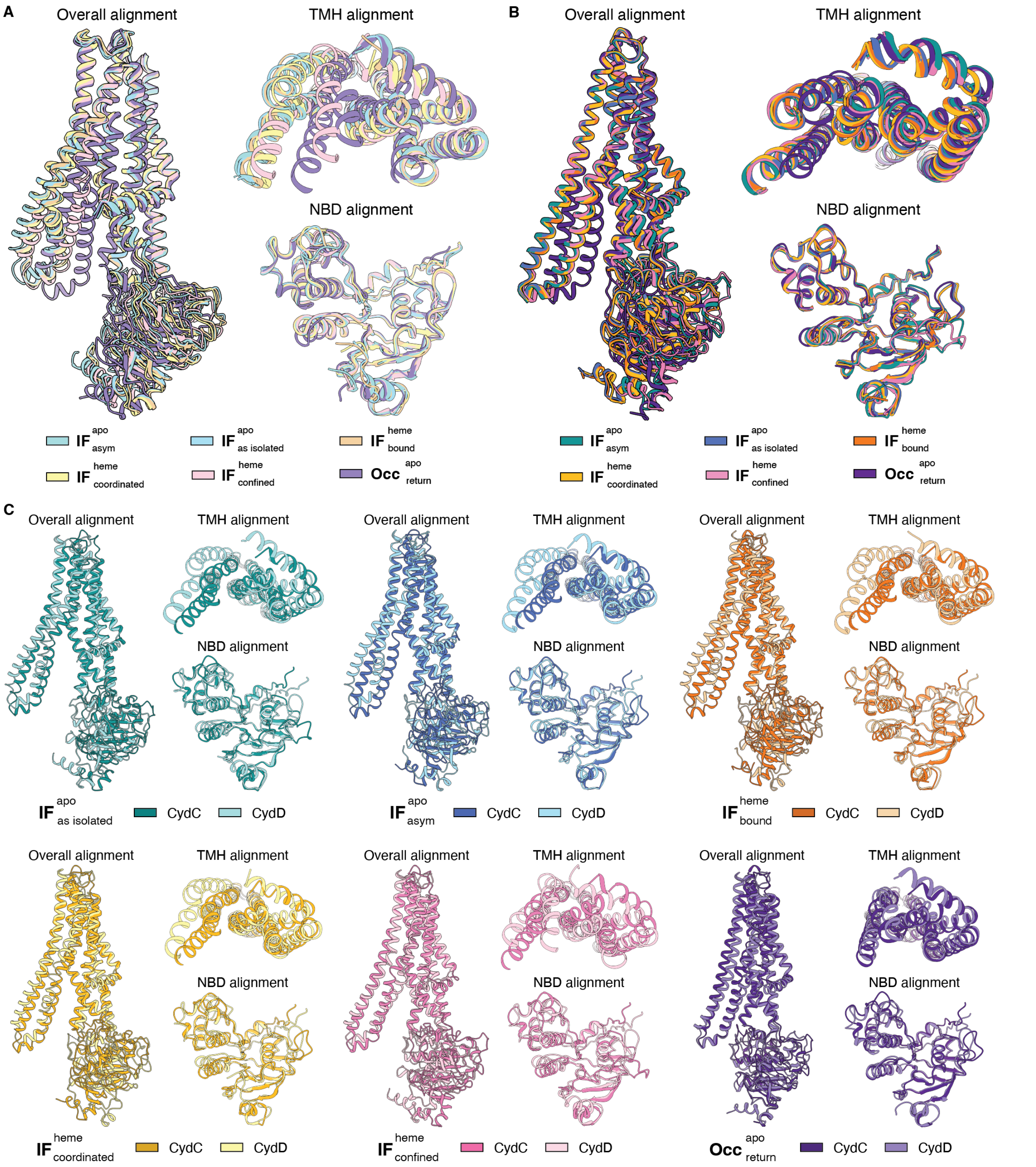

**Fig. S14 – Changing structural symmetry during the transport cycle states of CydDC**. **(A)** Structural superimposition of CydD conformations obtained by cryo-EM. **(B)** Structural superimposition of CydC conformations obtained by cryo-EM. **(C)** Structural superimposition of subunits representing each conformational state. Structural alignments are shown between subunits, TMH domains, and NB domains, respectively.

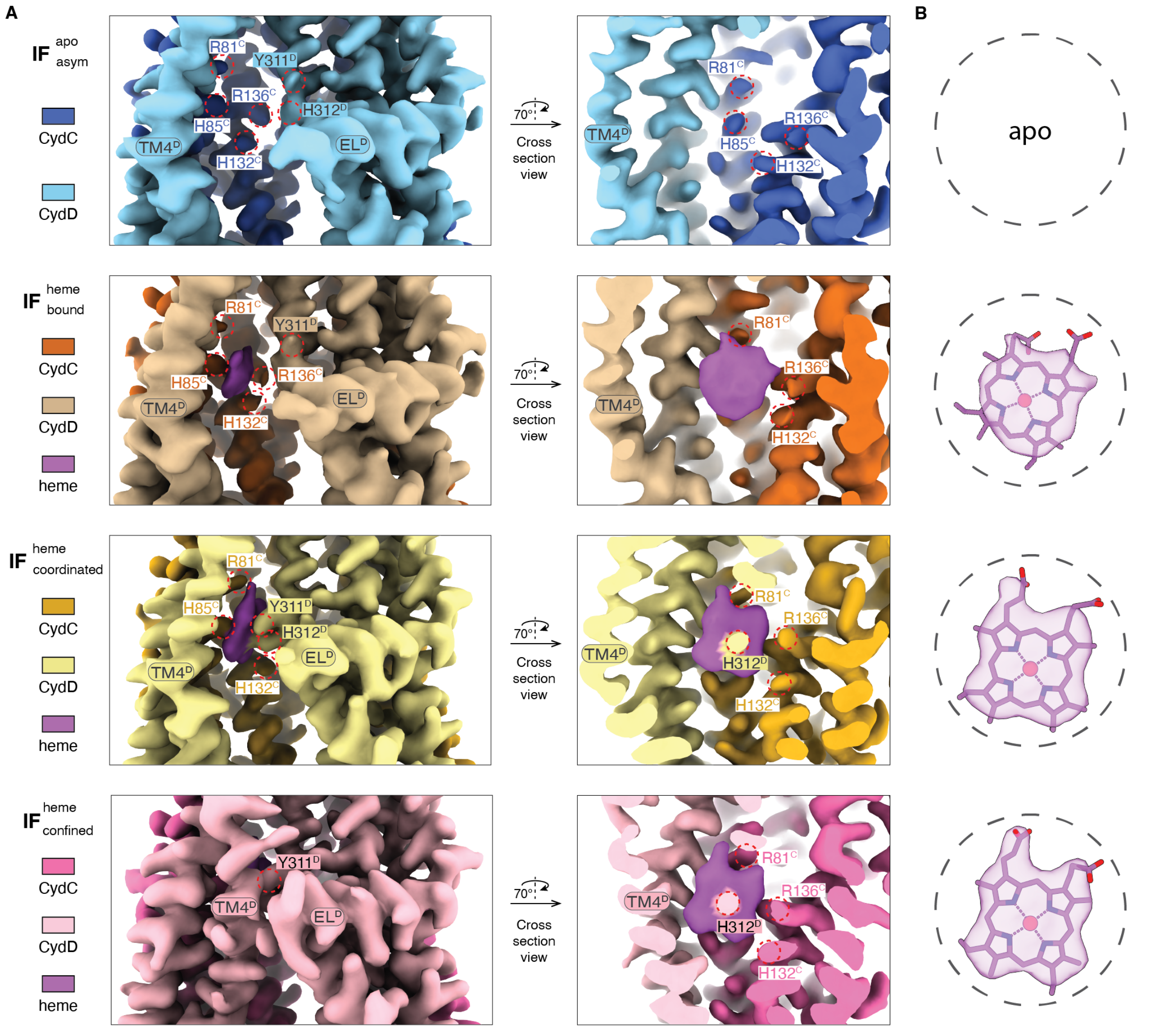

**Fig. S15 – Location and binding mode of heme in different CydDC conformations. (A)** Local density map of $\mathbf{IF}^{\mathrm{apo}}_{\mathrm{asym}}$, $\mathbf{IF}^{\mathrm{heme}}_{\mathrm{bound}}$, $\mathbf{IF}^{\mathrm{heme}}_{\mathrm{coordinated}}$, and $\mathbf{IF}^{\mathrm{heme}}_{\mathrm{confined}}$ at the heme binding site shown in side and cross-section views. **(B)** Heme models fitted into corresponding densities in maps of each conformational state. Density features of heme highlight that the heme molecule is rigidified during the process of binding and confinement.

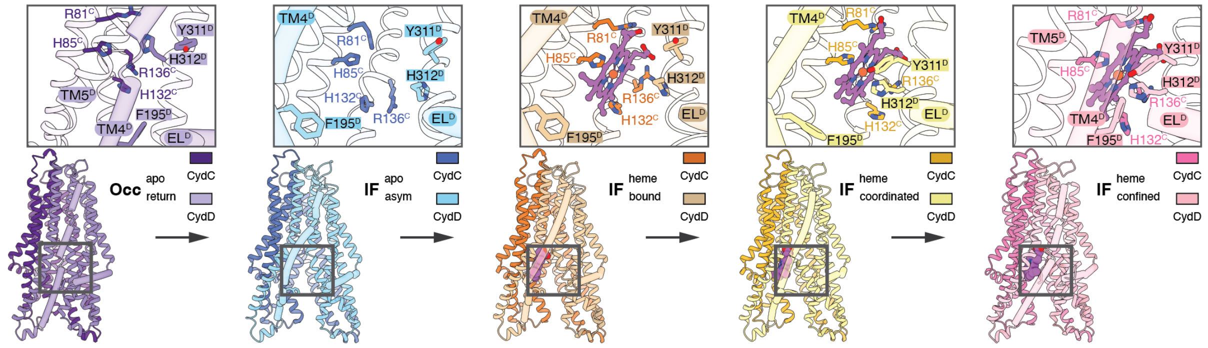

**Fig. S16 – Mechanism of heme binding.** Overview of TMH conformations during heme binding, coordination, confinement and translocation. Closeup side views show TMs and residues that form the heme binding site and the lateral entry gate. Heme is shown as purple ball-and-stick model. EL1$^{D}$ and TM4$^{D}$ are shown as tubes.

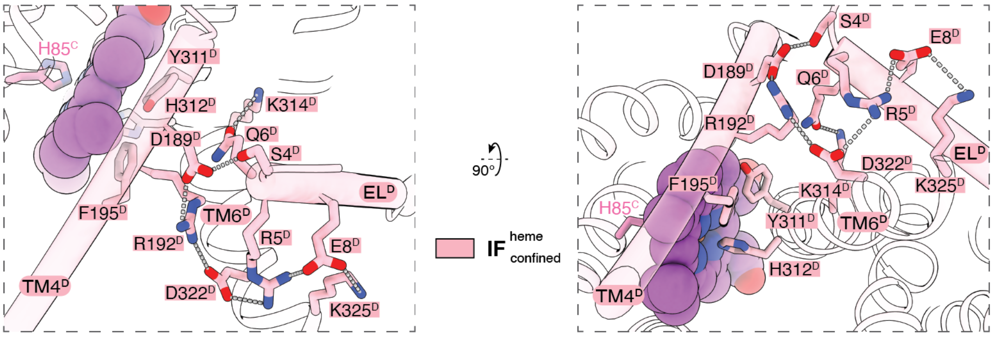

**Fig. S17 – Molecular details of the heme binding site and the membrane exposed entry gate.** Closing of the lateral substrate entry site is facilitated by the formation of an electrostatic interaction network between residues of EL$^{D}$ and TM4$^{D}$. Electrostatic interactions are indicated by dashed lines.

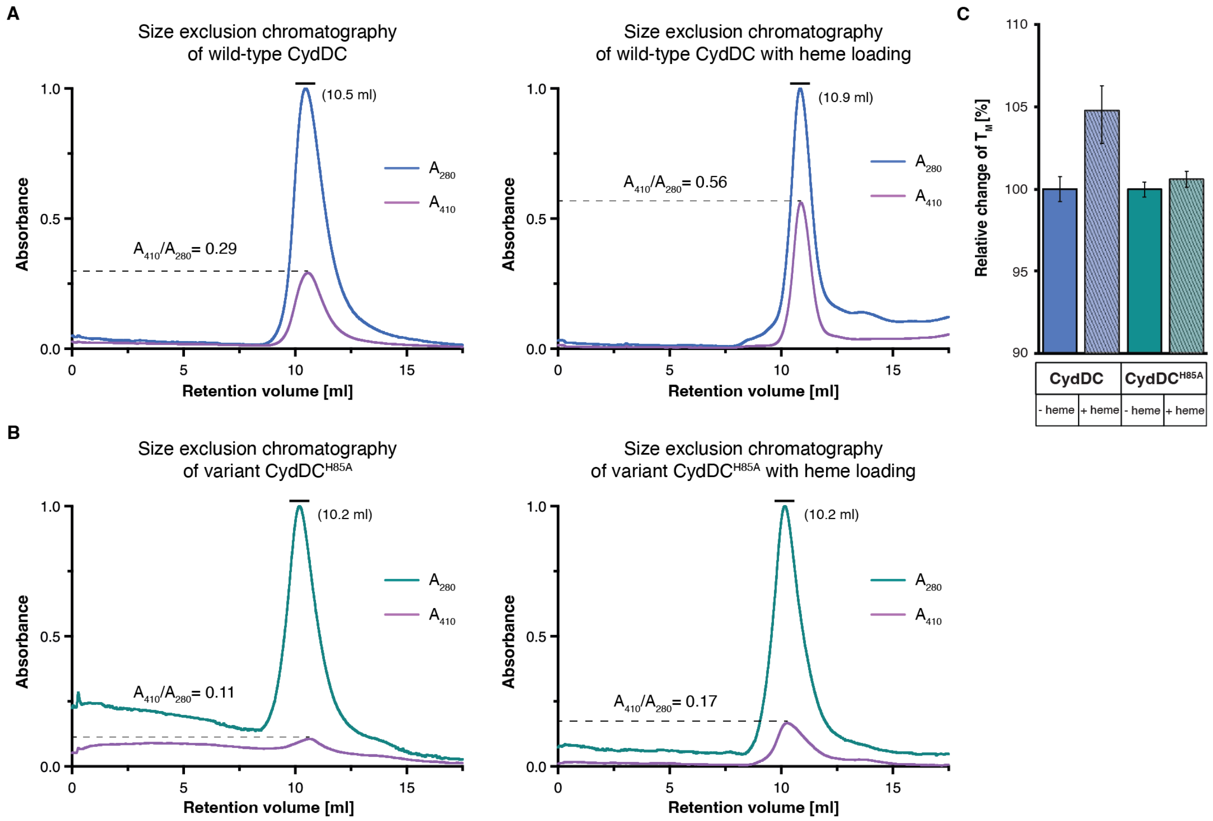

**Fig. S18 – Heme binding properties of CydDC$^{wt}$ and CydDC$^{H85A}$. (A)** Size exclusion chromatograms (SEC) of CydDC (wt) in the absence and presence of exogenously added heme (applied before the experiment). Wild-type CydDC contains bound heme from native source, which co-migrates with the peak fraction of the protein. Addition of extra heme increases the intensity of the $A_{410}$ signal indicating greater heme occupancy. **(B)** SEC of CydDC$^{H85A}$ in the absence and presence of exogenously added heme (applied before the experiment). The chromatograms show that native heme does not co-purify with the mutant variant CydDC$^{H85A}$. Furthermore, adding exogenous heme does not have the same effect observed with the wild-type CydDC. The slight increase in the $A_{410}$ signal is attributed to heme molecules migrating into hydrophobic regions of detergent micelles. Relative ratios of the peak intensities between $A_{410}$ (heme component) and $A_{280}$ (protein component) are given. **(C)** Relative change of $T_M$ in the absence and presence of exogenously added heme molecules to purified samples of CydDC$^{wt}$ and CydDC$^{H85A}$.

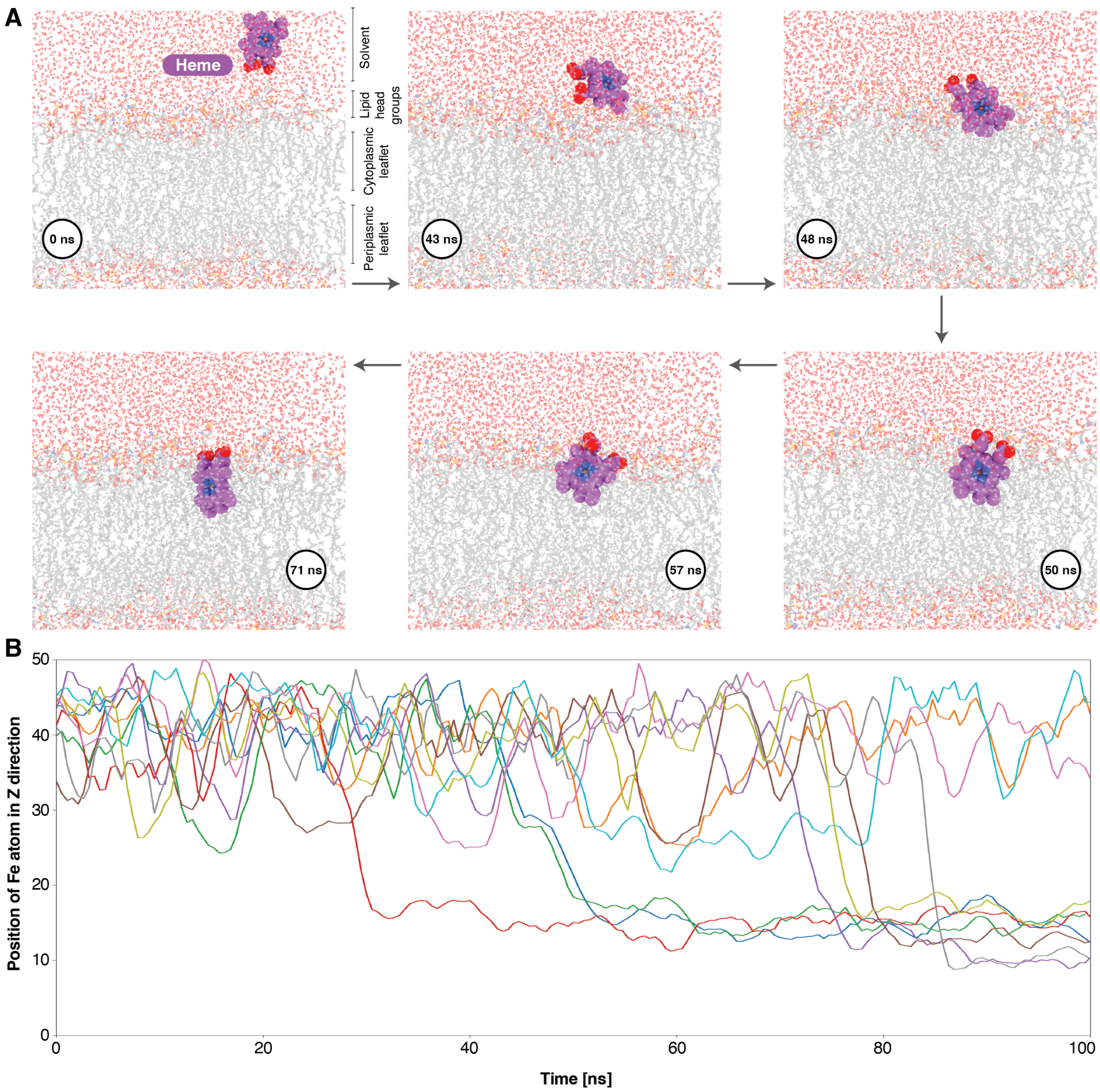

**Fig. S19 – Heme partitions spontaneously into lipid bilayers**. **(A)** Molecular dynamics simulations show spontaneous binding and insertion of heme into a lipid bilayer composed of 70 % POPE, 25% POPG, and 5% CL lipids. Heme establishes a stable position in the membrane during 100-ns simulation runs. **(B)** During MD runs, position of the heme was tracked by the distance of the Fe atom to the center of the membrane in the Z-axis direction. Graph colors represent different simulation runs.

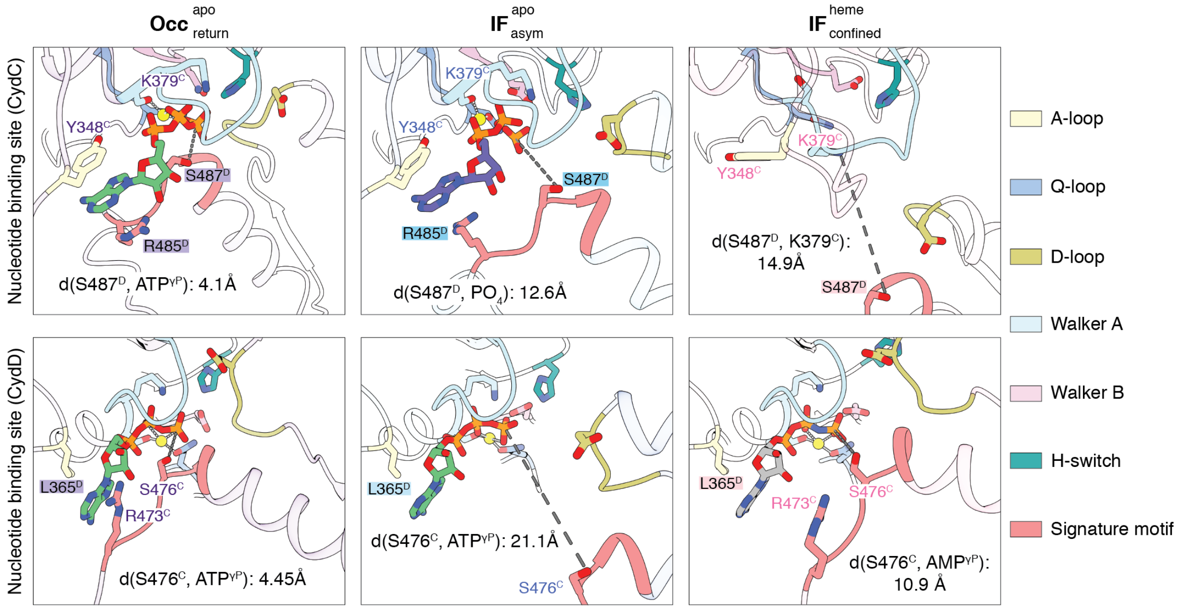

**Fig. S20 – Conformational variety of nucleotide binding sites.** Top panel: closeup view of NBS$^{C}$ conformations in the $\mathbf{Occ}^{apo}_{return}$ (ATP), $\mathbf{IF}^{apo}_{asym}$ (ADP + Pi), and $\mathbf{IF}^{heme}_{confined}$ (apo) states. Distances between the γ-phosphate and S487$^{D}$ (signature loop), and between K379$^{C}$ of the Walker A domain and S487$^{D}$ are indicated by dashed lines. Bottom panel: closeup view of NBS$^{D}$ conformations in the $\mathbf{Occ}^{apo}_{return}$ (ATP), $\mathbf{IF}^{apo}_{asym}$ (ATP), and $\mathbf{IF}^{heme}_{confined}$ (AMP-PNP) states. Distances between the γ-phosphates of ATP/AMP-PNP and S476$^{C}$ of the signature loop are indicated by dashed lines. Conserved structural motifs of nucleotide binding and hydrolysis sites are highlighted. ATP, green; ADP, purple; AMP-PNP, grey; phosphate, orange; $Mg^{2+}$, yellow.

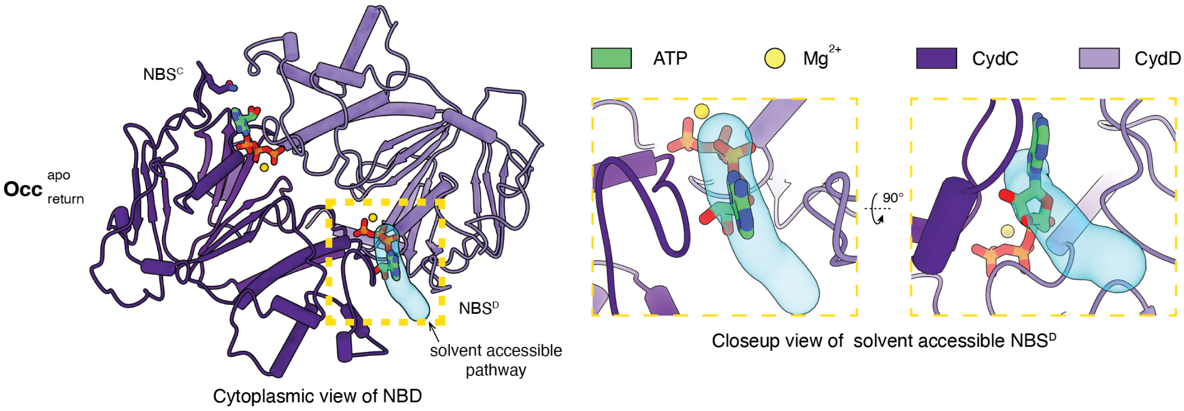

**Fig. S21 – Local environment and accessibility of NBS$^{D}$ and NBS$^{C}$.** Left: ribbon model of the interlocked NBD dimer in the $\mathbf{Occ}^{apo}_{return}$ state from turnover dataset of CydDC$^{E500Q}$. A predicted pathway between the aqueous environment and NBS$^{D}$ is depicted in blue. Right: closeup views of NBS$^{D}$ in side and top view orientation show that bound ATP is not shielded from the local environment. Weaker atomic interactions and exposure to the surrounding environment explain the observed reduced occupancy and possibility of exchange of nucleotide at NBS$^{D}$ in the $\mathbf{Occ}^{apo}_{return}$ state. ATP, green; $Mg^{2+}$, yellow; CydC, light purple; CydD, dark purple.

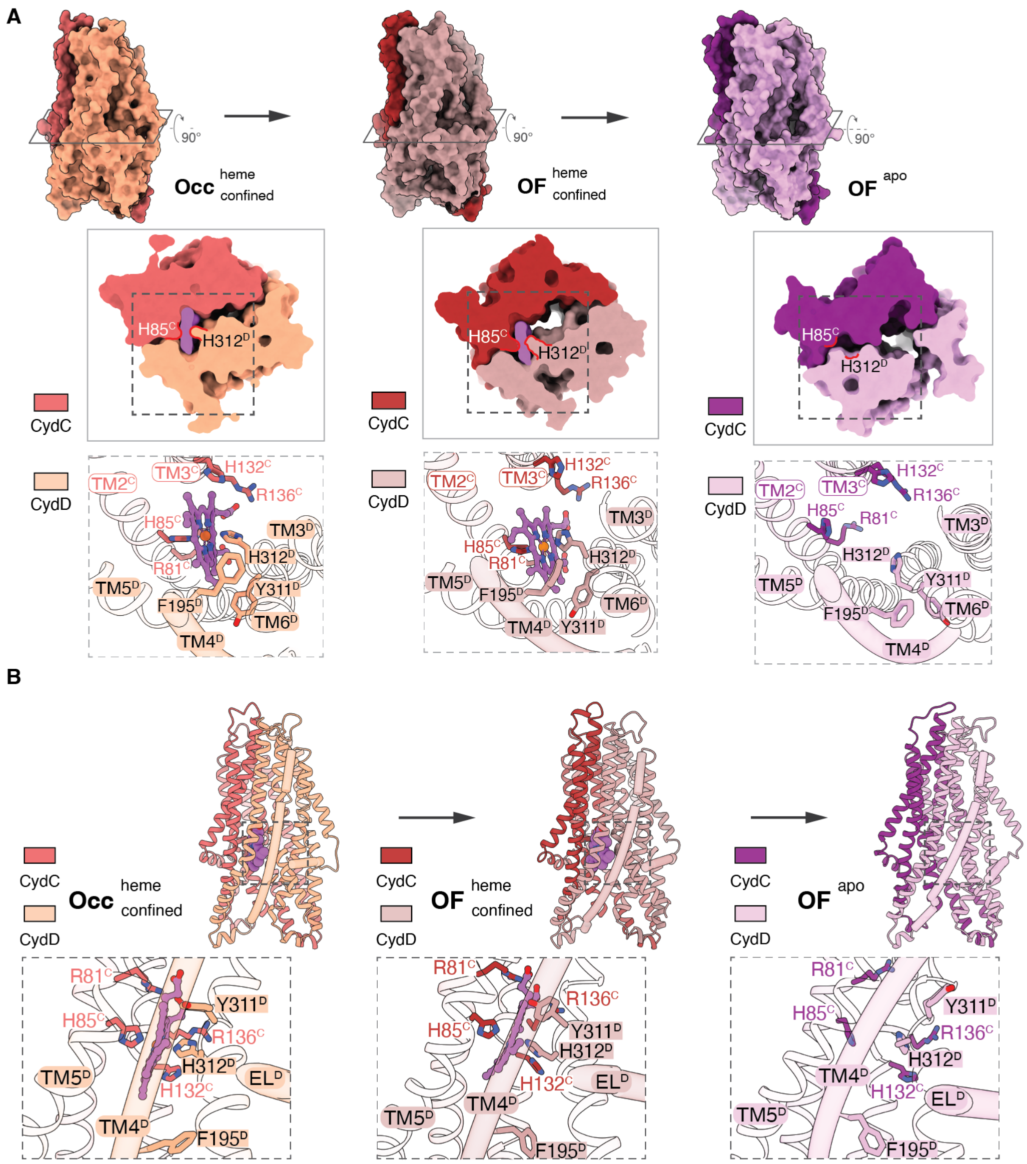

**Fig. S22 – Simulated outward-facing (OF) conformations of CydDC.** (**A**) Closeup views show residues involved in heme binding. Surface model cross sections illustrate the changing shape and size of the internal cavity in different conformational states. Surface model side views highlight the change in overall shape of the TMH region in different conformational states. (**B)** Closeup side views show changing orientations of TMs and heme-binding residues during conformational transitions of the predicted outward-facing states. Heme is shown as purple ball-and-stick model. EL1$^{D}$ and TM4$^{D}$ are shown as tubes.

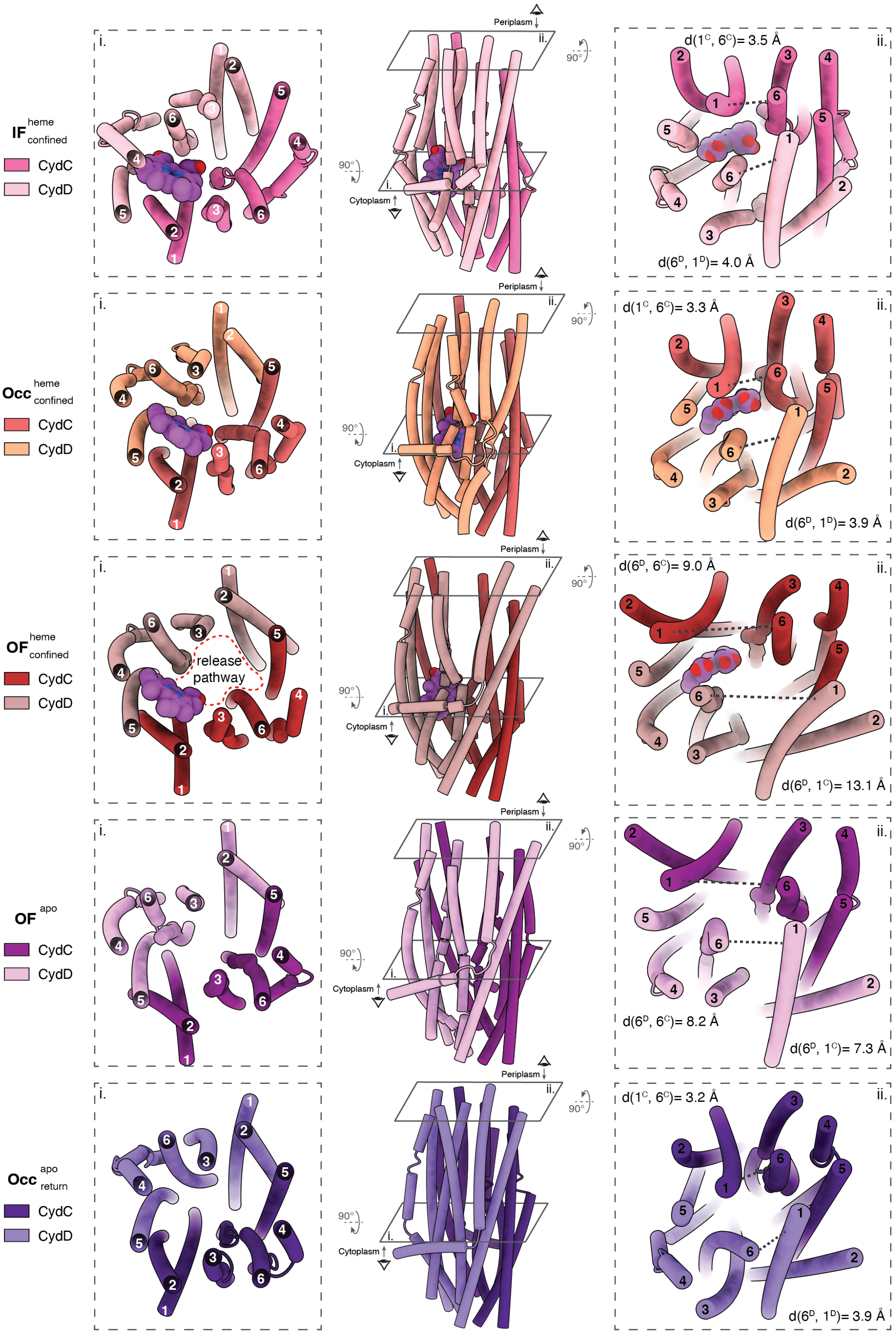

**Fig. S23 – Transmembrane helix rearrangement during the proposed heme release mechanism.** Center: side views of CydDC TMH regions during heme confinement and release toward the periplasm. Left: cross section views of TMH conformations at the level of the heme-binding site. Right: cross-section views of TMH conformations at the level of the periplasmic gate. Distances between TMs 1 and 6 of CydD and CydC (critical for gate opening and closing) are indicated. Periplasmic loop regions are omitted for clarity.

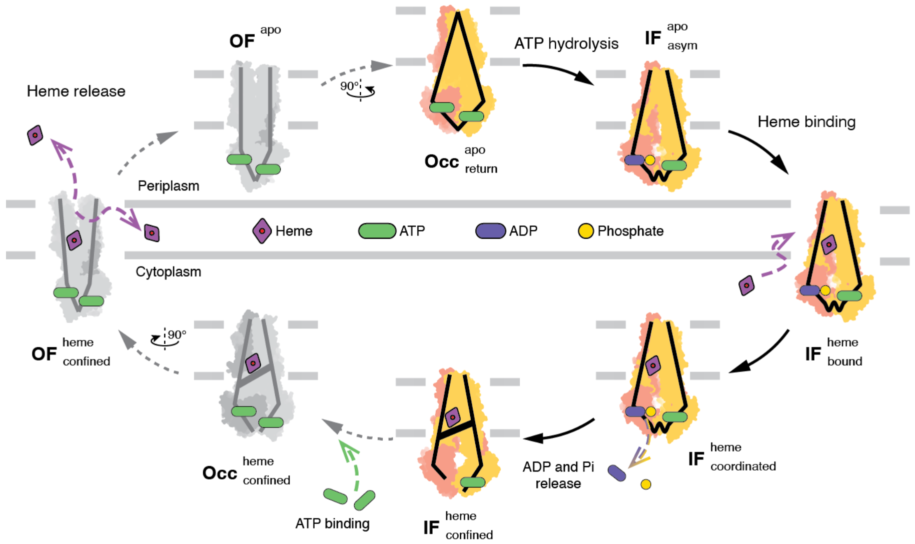

**Fig. 24 – Proposed model of heme transport facilitated by CydDC.** Energy of ATP hydrolysis at $NBD^{C}$ converts CydDC from the $\mathbf{Occ}^{\mathrm{apo}}_{\mathrm{return}}$ state to the $\mathbf{IF}^{\mathrm{apo}}_{\mathrm{asym}}$ state. Binding of heme to its dedicated pocket causes closing of the lateral substrate gate and triggers the release of ADP + Pi caused by conformational changes of the nucleotide-binding site of CydC. Release of heme occurs independent of ATP hydrolysis but requires binding of ATP. Separation of TM lobes and rearrangement of residues around the heme-binding pocket cause the release of heme either towards the periplasmic space or to the periplasmic bilayer leaflet of the membrane. Conformations obtained by cryo-EM are shown as yellow and red schematic models. Conformations predicated by MD simulations are shown as light and dark grey models.

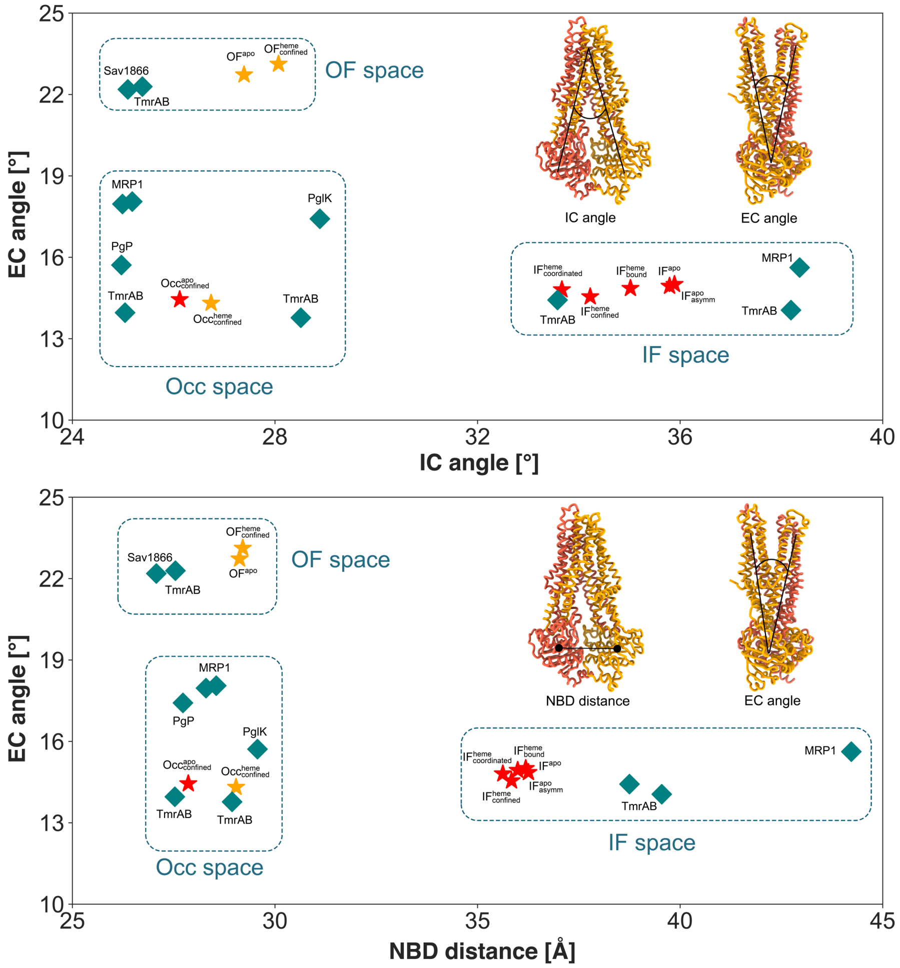

**Fig. S25 – Conformational space of ABC transporters.** Conformation space of type IV ABC transporters defined based on three parameters: the intracellular gate angle (IC angle), the extracellular gate angle (EC angle), and the distance between two nucleotide-binding domains (NBD distance). Cryo-EM structures from this study are displayed as red stars, the three modeled conformations are shown as orange stars, previously published structures of ABC transporters are presented as teal diamonds. The clustering of structures in three major conformations [(inward-facing (IF), outward-facing (OF), and occluded (Occ)] is highlighted by dashed rectangles. The IC angle is described as the angle between two axes: axis 1 between the center of mass of the extracellular part of the TMH domain and the center of mass of the intracellular parts of $TM1^{C}$, $TM2^{C}$, $TM3^{C}$, $TM6^{C}$, $TM4^{D}$ and $TM5^{D}$; and axis 2 between the center of mass of the whole extracellular part of the NBD region and the center of mass of the intracellular parts of $TM1^{D}$, $TM2^{D}$, $TM3^{D}$, $TM6^{D}$, $TM4^{C}$, and $TM5^{C}$. We define the EC angle as the angle between two axes: axis 3 between the centers of mass of the NBDs and the extracellular parts $TM1^{C}$, $TM2^{C}$, $TM3^{C}$, $TM4^{C}$, $TM5^{D}$, and $TM6^{D}$; and axis 4 between the center of mass of the NBDs and of the extracellular parts $TM1^{D}$, $TM2^{D}$, $TM3^{D}$, $TM4^{D}$, $TM5^{C}$, and $TM6^{C}$. The NBD distance is between the centers of mass of two NBDs (N-terminal loops and C-terminal helices were excluded). PDB IDs: TmrAB: 6RAG (IF wide), 6RAF (IF narrow), 6RAM (Occ), 6RAK (Occ), and 6RAJ (OF); MRP1, 5UJA (IF), 6BHU (Occ), and 6UY0 (Occ); Pgp, 6C0V (Occ); PglK, 5C73 (Occ); and Sav1866, 2ONJ (OF).
